# Can We Extract Physics-like Energies from Generative Protein Diffusion Models?

**DOI:** 10.1101/2025.11.28.690021

**Authors:** Sudeep Sarma, Harrison Truscott, Da Xu, Kendall Reid, Lee-Shin Chu, Jacky Chen, Jeffrey J. Gray

**Affiliations:** Department of Chemical and Biomolecular Engineering, Johns Hopkins University, Baltimore, MD 21218, USA; Bioinformatics Program, Center for Biotechnology Education, Johns Hopkins University, Baltimore, MD 21218, USA; CMU-Pitt Computational Biology, Dept. of Computational & Systems Biology, University of Pittsburgh, Pittsburgh, PA 15260, USA

**Author notes:** These authors contributed equally (Sudeep Sarma, Harrison Truscott). Computational Engineering Division, Lawrence Livermore National Laboratory, Livermore, CA 94550, USA;. Also at: Program in Molecular Biophysics, Johns Hopkins University, MD 21218, USA; Data Science and AI Institute, Johns Hopkins University, MD 21218, USA.

## Abstract

Diffusion models have emerged as the state-of-the-art method in generative artificial intelligence (AI) and have shown great success in image synthesis, video generation, molecular design, and protein structure prediction. For biophysical problems, such as protein folding and association, a fundamental question in diffusion-based methods is how their learned functions correspond to thermodynamics. In this paper, we study diffusion models through the lens of theoretical biophysics, analyzing their underlying formulation of potentials and exploring their applications in scoring protein interactions. We develop simple theories rooted in statistical physics that relate thermodynamic potentials to the negative log of the probability of observing a system in a particular state. We include dimensional analysis of diffusion model equations and a table mapping AI and physics jargon. We then test a diffusion model’s ability to capture learned energies as negative log-likelihood values, −log *p*_0_(*x*_0_), by integrating over the diffusion-generated path or a probability flow path. We test these integrals on a simple 1D Gaussian mixture diffusion model and a protein-docking diffusion model, DFMDock. In the 1D case, we find that integration over both diffusion and flow paths can accurately recover ground truth probabilities. When we extract the learned docking energies for cases where DFMDock succeeds, we observe energy funnels with the minimum energy near the experimental docked structure, like those we observe with Rosetta, an empirically tuned physics-based biomolecular modeling suite. The learned energy performs comparably or outperforms Rosetta interface energy in 9 out of 25 cases at ranking the correctness of docked poses. These data show that we can extract a relevant learned energy function from a diffusion model and compare it to physical energy functions.

## I. Introduction

Diffusion models are a class of deep-generative models that have shown great promise in image [1–3] and video generation [4], protein design [5, 6], and protein structure prediction [7–9]. Diffusion models were originally formulated by Sohl-Dickstein *et al*. [10] in analogy with nonequilibrium thermodynamics. They work in two stages. In forward diffusion, a data distribution is transformed to a prior (typically a Gaussian) over a series of time steps to learn an effective force field (the score), which is the time-dependent gradient of the logarithmic probability density function [11]. The score is learned by a deep neural network matching the noise added in training. One can then use the learned score to sample the underlying probability distribution of the data during a backward diffusion process starting from the prior. Leading image generation tools such as DALL-E [1], Stable Diffusion [2], and Midjourney [3] and text-to-video models like Sora [4] are all based on diffusion models. In protein design, breakthroughs such as RFDiffusion [5] and Chroma [6] have led to novel biomolecules that fold and bind in the lab. RFDiffusion represents amino acid residues as rigid frames and generates protein backbones from an ideal-gas-like prior. Chroma diffuses in polymer backbone torsion space, respecting conformational statistics of polymer ensembles.

In 2024, Deepmind released AlphaFold3 (AF3) [7], replacing AlphaFold2’s [12] structure module with a diffusion module capable of predicting structures of proteins, protein complexes and post-translational modifications. AF3 drew from previous pioneering studies of diffusion for biomolecules [5, 6, 13], and has since inspired several derivative models (Chai-1 [8], Boltz-2 [14], and OpenFold3 [15]). The incredible success of AF3 and other protein deep learning models is in some ways surprising because these methods outperform physics-based methods like Rosetta [16, 17], which is based on decades of research into conformational sampling and scoring that follow energy functions derived from or inspired by physical principles. It raises the question of whether these high-performing deep-learning methods have implicitly learned a potential akin to nature’s free energy function [18].

The generative model training objective is to learn to sample from a distribution that matches known protein structures. In other words, the sampling probability should be maximized at real protein structures, similar to Anfinsen’s dogma [19], which states that naturally folded protein structures are at their lowest free energy state (i.e., in equilibrium). There are many functional forms that can have minima at the same (folded) states, so an AI protein model’s learned energy function does not need to match nature’s energy function. Additionally, diffusion models do not typically allow direct access to the probability of a given state. Here, we seek general approaches to extract information from a diffusion model in the form of its learned potential. In this way, we can compare this potential to physics-based energy functions and consider how AI and physics-based methods might be combined or improved.

To better understand the inherent potentials that diffusion models learn, we focus on a tractable and interpretable model for the protein-protein docking problem [20]. Previously, we introduced DFMDock (Denoising Force Matching Dock), a diffusion model for rigid-body protein-protein docking [21]. It is trained by adding translational and rotational noise to experimentally determined, bound protein complexes, which the model then learns to reverse through a denoising force-matching objective. During inference, the model inputs two unbound monomers and generates the structure of the bound complex. For our model study, we use a DFMDock version trained only on the translation docking space. In this way, we can also make comparison with a 1D translational diffusion model of a Gaussian mixture.

We formulate diffusion models in the language of statistical thermodynamics and explore how protein diffusion models like DFMDock score biomolecular structures and complexes. Statistical thermodynamics provides a framework for predicting the static and dynamic properties of a many-body system from its microscopic constituents and their interactions [22]. For systems in equilibrium, the Boltzmann distribution relates the probability of finding a system in a particular configuration *x* to its energy *E*(*x*) as:

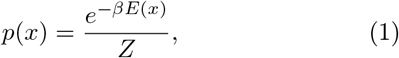

where *β* is the system’s inverse temperature and the partition function *Z* is integrated over the space of all possible system configurations:

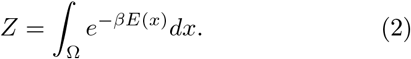

We assess a diffusion model’s ability to capture energies by introducing a framework to extract negative log-likelihood values (NLL) of a sample *x*_0_ at diffusion time *t* = 0, −log *p*_0_(*x*_0_). Utilizing the Boltzmann relation, we establish a relationship between this learned energy, −log *p*_0_(*x*_0_), and the thermodynamic free energy of the configuration, *E*_0_(*x*_0_).

This relationship connects deep learning models with physics by mapping learned potentials with fundamental thermodynamic quantities. We explore multiple approaches to calculating *E*_0_(*x*_0_) by integrating the implicitly learned energy gradient using ∇_*x*_ log *p* over both noisy diffusion paths and the smooth deterministic path generated by the probability flow ordinary differential equation (ODE) as described in [11]. As a test for our approach, we train a simple 1D diffusion model to learn a Gaussian mixture and compare generated samples’ −log *p*_0_(*x*_0_) values using our integral methods to the ground truth values. We then apply the same approach to a protein docking diffusion system and compare the learned energies with Rosetta energies (**Fig. 1**).

**FIG. 1.**
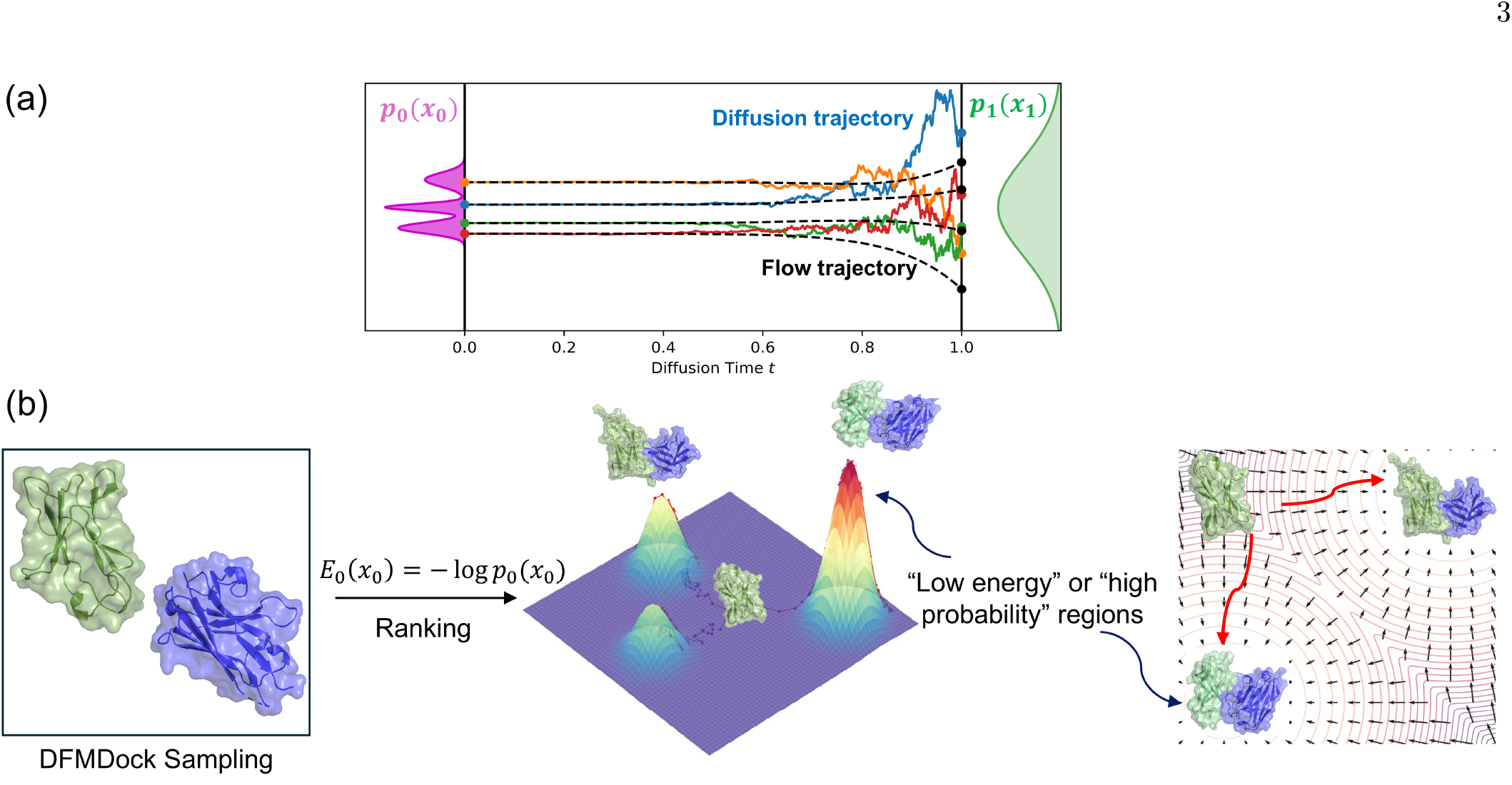
(a) Diffusion (SDE paths, solid colors) and flow (ODE paths, dashed black) trajectories from a noise (prior) distribution, *p*_1_(*x*_1_), at time *t* = 1 to the data distribution, *p*_0_(*x*_0_), at time *t* = 0. (b) Negative log-likelihood values of docked protein complexes are a measure of learned protein-protein interaction energy.

To our knowledge this is the first attempt at extracting potentials from a biomolecular score-based diffusion model and comparing with a physical energy function. We believe this is also the first exploration of using paths other than the probability flow ODE for likelihood computation. We aspire to improve the interpretability of deep-learning models for protein-protein interactions and open the door to building more robust physics informed tools for biomolecular engineering.

## II. Related Work

Protein-protein interactions (PPIs) drive a wide range of biological and chemical processes. They are involved in most cellular functions in living organisms, such as signaling, regulation, and recognition. Understanding their three-dimensional structures provides atomic-level insight into the mechanisms of these functions [23]. Traditional protein docking methods use sampling algorithms such as local-shape matching and Monte Carlo search algorithms to generate plausible docked conformations [24, 25]. Scoring functions, such as the Rosetta interface score function, can then be used to evaluate the physical energy of these docked conformations [26]. However, these search and scoring protocols are computationally intensive. Recently, new generative deep learning methods, especially diffusion models, have shown promise in addressing the protein-docking problem.

The first diffusion model applied to molecular docking, DiffDock, learned small-molecule–protein docking with denoising score matching on the translation, rotation, and torsion spaces of small molecules [13]. The developers subsequently introduced DiffDock-PP, a diffusion model for rigid protein docking [27] that learned to translate and rotate unbound protein structures to their bound conformations. DiffMaSIF is a diffusion framework for protein docking that uses an encoder-decoder architecture to learn physical surface patch complementarity [28]. LatentDock first trains a variational autoencoder on protein sequences and structures and then diffuses in the latent space to produce the final conformations of the protein complex [29].

These models have shown promising empirical results. Other recent work has focused on the theoretical underpinnings of diffusion models. Diffusion models derive from non-equilibrium physics [10], but several papers have also conceptualized diffusion models using tools of equilibrium statistical mechanics. Ambrogioni *et al*. [30] showed that generative diffusion models undergo second-order phase transitions as described by mean-field theory. Sclocchi *et al*. [31] found that during the backward diffusion process, the system undergoes a phase transition at particular noise levels after which high-level features like the class of an image are irreversibly decided. Biroli *et al*. [32] showed that the backward diffusion process has three dynamical regimes, and they characterized the cross-overs between them as speciation (analogous to symmetry-breaking phenomena) and collapse (analogous to glass transition). There have been several attempts to extract meaningful energy scores from protein structure models. One strategy is to use an additional proxy network trained on a downstream task. Roney and Ovchinnikov [18] showed that AlphaFold2’s confidence module, trained solely on training-time accuracy of the structure module’s predictions, nonetheless worked as such a proxy. They demonstrated that the confidence score from AlphaFold2 (even deprived of the evolutionary signal from multiple sequence alignment) is capable of discriminating between native and non-native structures and is correlated with Rosetta energies. Liu *et al*. [33] extended the methodology to AlphaFold3’s confidence module, which allowed ranking protein-ligand and protein-nucleic acid interactions. Zaidi *et al*. [34] trained a neural network to predict molecular properties, including experimentally determined energies, using embeddings from a pre-trained molecular structure generative diffusion model as input. These results attest to the usefulness of leveraging high-quality pretrained generative protein structure models for energy and ranking, but the use of an auxiliary network limits our ability to interrogate what, if any, physics is being learned from and used for structure generation by the original model. Energy-based models (EBMs) are a class of generative models that directly learn a scalar energy function *E*_*t*_(*x*) = −log(*p*(*x*_*t*_, *t*)) −log(*Z*), rather than the energy gradient ∇_*x*_ log *p*(*x*_*t*_, *t*) learned by score-based diffusion models [35]. Thus, they have been promising candidates for physically-informed simultaneous inference and energy prediction for biophysical generative models. DSMBind [36] is an EBM whose energy function is optimized by matching its gradient to that of the forward diffusion process to predict binding energies of protein-protein interactions. DockGame [37] learned an energy function via supervision from physics-based models and self-supervision via score-matching with diffusion models for rigid multimeric protein docking. EBMDock [38] uses an energy-based learning framework and Langevin dynamics sampling for docking pose prediction. Borisiak *et al*. [39] trained an energy-based diffusion network to score mutations on peptides and CDR3 loops within TCR-pMHC interfaces. Roney *et al*. [40] trained an energy-based protein diffusion model, ProteinEBM, and demonstrated correlation between its energy function and both Rosetta and experimental energies. These results have all shown great promise, yet standard EBMs do not trivialize the problem of free energy computation, as they have difficulty determining the relative normalization of sparse data regions [41, 42] and tend to have noisy *t* = 0 energies [40, 42].

The connection between sampling likelihood and energy (eq. (1)) has led several groups to explore the connection between physical force, *F*(*x*) = ∇_*x*_*E*(*x*), and the diffusion-model learned score, ∇_*x*_ log *p*(*x*_*t*_, *t*), near time *t* = 0. Arts *et al*. [43] used an energy-based model trained on coarse-grained molecular dynamics (MD) equilibrium data to compute the score at time *t* = 0 for use as an approximate coarse-grained force field. Raja *et al*. [44] used the score at *t* = 0 as an atomistic force-field to define a stochastic process exploring molecular conformations, then sampled high-probability transition paths between states by minimizing the Onsager-Machlup functional of this process over the space of possible paths. He *et al*. [45] developed a framework for free energy estimation on stochastic interpolant models, a class of models that builds upon and extends score-based diffusion models [46]. Using a trained stochastic interpolant model, consisting of two energy models and a learned velocity field transforming between the two, they developed work integrals along stochastic paths connecting the two distributions to estimate the difference in total free energy (*k*_*B*_*T* log *Z*) between the two distributions. Their method, FEAT, is in many ways the equivalent of diffusion model likelihood estimation for stochastic interpolant models, and there are many overlaps in strategy and findings (which we will return to in **Section V**) though their use of interpolated energy models makes their work integrals differ from ours. Together, these methods highlight the benefits of utilizing the score for downstream tasks: it is accessible from both score-based diffusion models and energy models via automatic differentiation, it has direct statistical meaning (and physical meaning, at time *t* = 0), and score-based analysis can be applied directly to any existing diffusion and energy model with no additional training or downstream fine-tuning.

The principle of extracting learned likelihood from score-based generative diffusion models was first introduced by Song *et al*. (2020) [11] using their probability flow ODE. By converting the diffusion process into an equivalent process using stochasticity-free ODEs, they could use the instantaneous change-of-variables formula from [47] to integrate the probability change along a flow ODE trajectory. By contrast, we formulate likelihood recovery in terms of the Fokker-Planck equation describing the diffusion process. The instantaneous change-of-variables formula is a special case of this Fokker-Planck formulation when integrated over the flow trajectory, thanks to the equivalence between the ODE and diffusion process [11, 47], meaning our formulation generalizes the original likelihood formula to any (piecewise-differentiable) path. Our paper extends the existing literature in three ways.

First, we build on the interpretation of diffusion models as capturing physical phenomena by extending the notion of score-as-force to times beyond *t* = 0 in concert with a (less-physical) time-dependent energy function. Inspired by this thermodynamic interpretation, we then derive a novel method of likelihood recovery from diffusion models by integrating the spatiotemporal log-probability gradient, generalizing existing likelihood recovery methods to arbitrary paths. Finally, we compare this learned energy to physical energy potentials for protein docking, thus contributing to the interpretability of diffusion models in protein structure prediction and design.

## III. Theory

### A. Background: Stochastic Differential Equations (SDEs) in Diffusion Models

The goal of generative modeling is to learn to transform a distribution of noise into samples that closely resemble a particular data distribution, such as the distribution of naturally occurring protein structures. In diffusion modeling, we first describe a transformation from data to noise—the *forward* diffusion process—and train a neural network to undo this transformation during the *reverse* diffusion process.

The forward diffusion process is obtained by incrementally noising a distribution of samples, *p*_0_, until the samples converge to a known prior distribution, *p*_1_. To describe the transformation of data to noise, the continuous variable *t* ∈[0, 1] is introduced, which indexes a continuum of probability distributions smoothly interpolating from *p*_0_ to *p*_1_. The variable *t* is referred to as “time” in the machine learning literature, but for a protein or other physical system, it actually represents an alchemical transformation. For example, RFDiffusion diffuses protein structures into an oriented, ordered ideal gas. We can draw parallels with alchemical free energy methods that gradually change one molecule into another along a transformation variable, allowing the calculation of the excess chemical potential through a nonphysical perturbation [48, 49]. To clarify the jargon between the AI and physical chemistry communities, we summarize the interpretation of the diffusion model terms in **Table I**, where we differentiate the alchemical time dimension [*τ* ] from the physical time dimension [*T* ].

**TABLE I.**
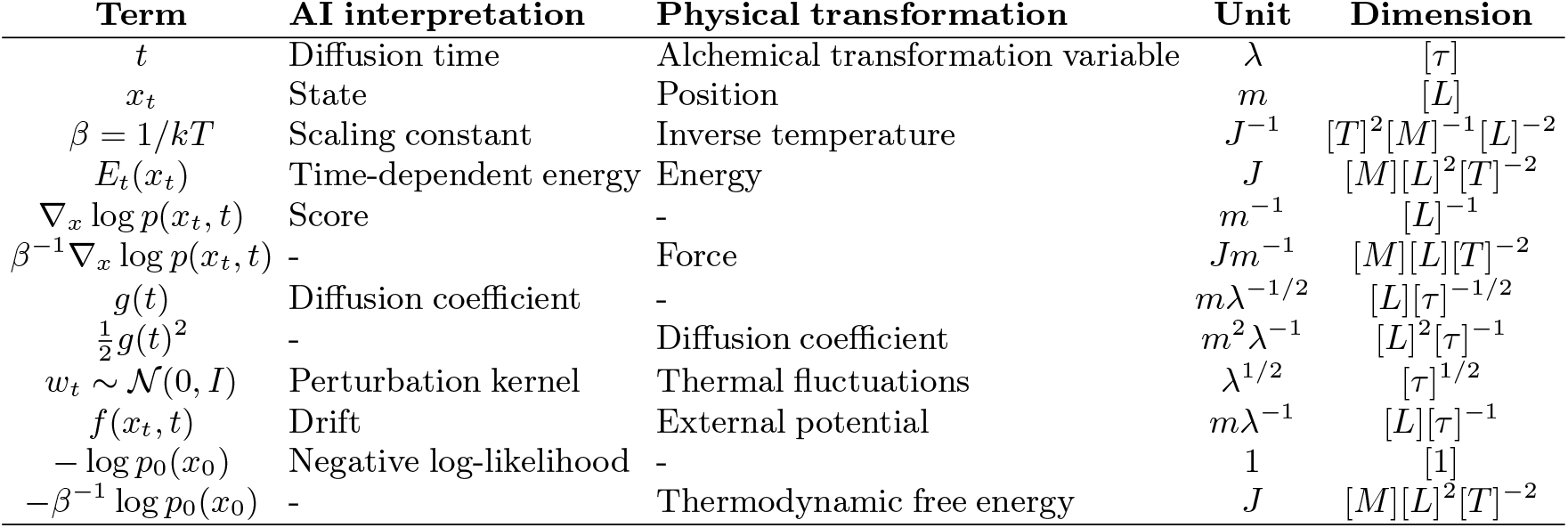
Mathematical terms of diffusion models and their interpretations as AI or physics jargon. Example units: *m*, meters; *J*, Joules. *λ* is used as an arbitrary unit of diffusion “time.” Dimensions: *T*, physical time; *τ*, diffusion/alchemical time; *L*, length; *M*, mass. Dimensional analysis deriving these units from the major equations is provided in **Section A** of the Supplementary Material.

| Term | AI interpretation | Physical transformation | Unit | Dimension |
| --- | --- | --- | --- | --- |
| $t$ | Diffusion time | Alchemical transformation variable | $\lambda$ | $[\tau]$ |
| $x_t$ | State | Position | $m$ | $[L]$ |
| $\beta = 1/kT$ | Scaling constant | Inverse temperature | $J^{-1}$ | $[T]^2[M]^{-1}[L]^{-2}$ |
| $E_t(x_t)$ | Time-dependent energy | Energy | $J$ | $[M][L]^2[T]^{-2}$ |
| $\nabla_x \log p(x_t, t)$ | Score | - | $m^{-1}$ | $[L]^{-1}$ |
| $\beta^{-1} \nabla_x \log p(x_t, t)$ | - | Force | $Jm^{-1}$ | $[M][L][T]^{-2}$ |
| $g(t)$ | Diffusion coefficient | - | $m\lambda^{-1/2}$ | $[L][\tau]^{-1/2}$ |
| $\frac{1}{2}g(t)^2$ | - | Diffusion coefficient | $m^2\lambda^{-1}$ | $[L]^2[\tau]^{-1}$ |
| $w_t \sim \mathcal{N}(0, I)$ | Perturbation kernel | Thermal fluctuations | $\lambda^{1/2}$ | $[\tau]^{1/2}$ |
| $f(x_t, t)$ | Drift | External potential | $m\lambda^{-1}$ | $[L][\tau]^{-1}$ |
| $-\log p_0(x_0)$ | Negative log-likelihood | - | 1 | [1] |
| $-\beta^{-1} \log p_0(x_0)$ | - | Thermodynamic free energy | $J$ | $[M][L]^2[T]^{-2}$ |

Dimensional analysis of key equations using physical units can be found in **Supplementary Material A**. Mathematically, the incremental noising of the data is formulated with a stochastic differential equation (SDE) known as the forward SDE:

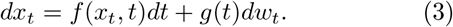

In the forward diffusion process, a given point *x*_0_ ∈ℝ_*n*_ sampled from the data distribution, *x*_0_ ∼*p*_0_, follows this SDE from time *t* = 0 to *t* = 1, arriving at some destination *x*_1_ in the prior distribution, *x*_1_∼ *p*_1_ [11]. The process is parameterized by two functions: the drift function, *f*(*x*_*t*_, *t*): ℝ_*n*_× ℝ→ℝ_*n*_, and the diffusion coefficient, *g*(*t*): ℝ →ℝ. While the AI jargon typically names *g*(*t*) as the diffusion coefficient, in statistical physics, the quantity 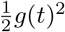 is usually given that name and has the units one would expect (**Table I**). For the purposes of this paper, we use the AI convention of referring to *g*(*t*) as the diffusion coefficient. *w*_*t*_ is a Wiener process describing stochastic Brownian motion, where *dw*_*t*_ is an infinitesimal random perturbation sampled at each time *t* from a Gaussian with mean 0 and variance *dt*. In DFMDock, *x*_*t*_ represents the relative position of the ligand relative to the receptor, and *dw*_*t*_ represents random translations of the ligand protein.

The forward SDE is constructed with specific choices *f* and *g* so that the resulting *marginal* probability distribution of this process, *p*(*x*_*t*_, *t*)—the probability of observing a point *x*_*t*_ at time *t*, marginalized over all initial points *x*_0_ and stochastic trajectories from *x*_0_ to *x*_*t*_—converges to the desired prior distribution *p*_1_ at *t* = 1. By definition, *p*(*x*_0_, 0) also matches our data distribution *p*_0_(*x*_0_). Moreover, *p*(*x*_*t*_, *t*) defines a continuum of distributions *p*(·, *t*), smoothly interpolating from *p*_0_ to *p*_1_.

In our 1D diffusion model and DFMDock, we use a “variance exploding” forward diffusion process [11] with 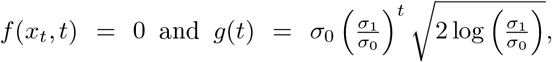, with *σ*_0_ and *σ*_1_ as hyperparameters. Because *f*(*x*_*t*_, *t*) = 0, the variance exploding diffusion process continuously adds pure zero-centered white noise to the data. As *t* increases, this causes the marginal distribution *p*(*x*_*t*_, *t*) to approach a Gaussian distribution, motivating the use of a zero-centered Gaussian prior for our choice of *p*_1_. However, as we cannot add an infinite amount of noise, we must assume that by *t* = 1, sufficient noise has been added that *p*(*x*_1_, *t*) is still approximately Gaussian. As a sum of zero-centered Gaussians is also Gaussian, we can compute the total added noise from time 0 to 1 by integrating the instantaneous variance, *g*(*t*)^2^*dt*:

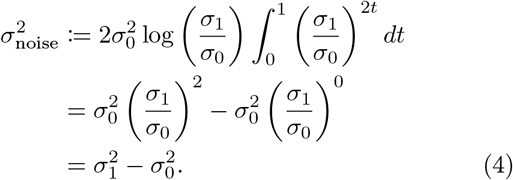

We can therefore compute the mean *µ*_prior_ = *µ*_data_ and variance 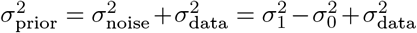 of the prior as the sum of two random variables, where *µ*_data_ and 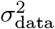 are the mean and variance of the data distribution respectively. If we have an estimate of our data variance, we can then set *σ*_0_ = *σ*_data_, which simplifies the prior to have variance 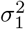.

The hyperparameters *σ*_0_ and *σ*_1_ thus respectively describe the assumed initial and final standard deviation of the marginal distribution *p*(*x*_*t*_, *t*). DFMDock uses *σ*_0_ = 0.1Å and *σ*_1_ = 30 Å, with 0.1 Å generally approximating uncertainty of atomic coordinates in a crystal structure. Our 1D diffusion model uses *σ*_0_ = 30 and *σ*_1_ = 70, which we selected based on experimental data on prior accuracy and likelihood recovery (**Supp. Fig. S3, Supplementary Material B**). For both priors, we use a Gaussian: *p*_1_(*x*_1_) = 𝒩 (*x*_1_; *µ*_prior_ = 0, *σ*_prior_ = *σ*_1_). For DFMDock, *µ*_prior_ = 0 means the center of the distribution has the centers of mass of both proteins at the same position.

During inference, we use the reverse SDE [50]:

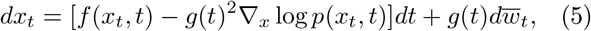

where time flows backwards from 1 to 0, *dt* is an infinitesimal negative timestep and 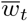 is a reverse-time Wiener process with variance −*dt*. This SDE models the reverse diffusion process: starting from *t* = 1 and a sample *x*_1_∼ *p*_1_, the marginal distribution of the reverse SDE transforms the prior distribution *p*_1_ into the data distribution *p*_0_ as *t* decreases from 1 to 0. ∇_*x*_ log *p*(*x*_*t*_, *t*), known as the *score*, is the spatial gradient of the log of the marginal probability of the process at time *t*.

By construction, this reverse SDE has an identical marginal distribution to the forward SDE [50]. The only unknown in eq. (5) is the score, so in score-based generative modeling, the diffusion model consists of a neural network, *s*_*θ*_(*x*_*t*_, *t*), parameterized by weights *θ* and trained to approximate ∇_*x*_ log *p*(*x*_*t*_, *t*) given a sample *x*_*t*_ and a time *t* (potentially conditioned on other aspects of the process, such as image class or protein sequence identity). Since the marginal probability *p*(*x*_0_, 0) of arriving at a point *x*_0_ via the reverse SDE should match the data distribution *p*_0_(*x*_0_), sampling a point *x*_1_ from the prior and following the reverse SDE using the learned score should produce samples *x*_0_ with the same probability as sampling from the data distribution directly.

The marginal distribution is generally intractable to solve at training time, as we only have a finite dataset of samples from *p*_0_ and thus cannot marginalize over the entire distribution. However, we can still evaluate the conditional probability *q*(*x*_*t*_, *t*| *x*_0_) of reaching a point *x*_*t*_ at time *t* when starting from a single point in the data distribution, *x*_0_, instead of the entire *p*_0_ distribution. Vincent (2011) [51] proved that matching the learned score with *q* conditioned on each of the data points in turn will converge to the marginal distribution as the size of the dataset grows. This results in the so-called denoising score matching objective that is used to train score-based diffusion models:

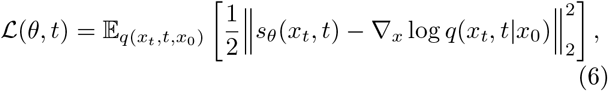

where the expectation is evaluated as a weighted sum over samples from the forward diffusion process starting at the points in the training dataset. In this way, score-based diffusion models implicitly incorporate information about all possible diffusion trajectories into the score. We reason that it should therefore be possible to recover a trained model’s understanding of the ground-truth probability distribution—and thus, its understanding of the ground truth *energy*—without needing to explicitly marginalize over all paths through diffusion space.

### B. Forces and Energies in Diffusion Models

In the case of protein docking or folding, our data distribution *p*_0_ is assumed to sample from the distribution of system states at thermal equilibrium, corresponding to stably folded or docked proteins. Thus, we can define an energy *E*_0_(*x*_0_) for any equilibrium state *x*_0_ based on *p*_0_:

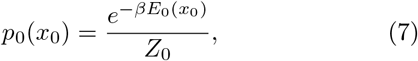

where *Z*_0_ is the partition function associated with the system at equilibrium, calculated by integrating over the space of all system states:

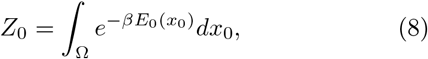

and *β* corresponds to the system’s inverse-temperature,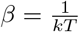.

Diffusion models are trained to allow sampling from *p*_0_, meaning they must have an implicit understanding of this energy function at equilibrium. However, they cannot output *p*_0_(*x*_0_) directly. Instead, as discussed earlier, score-based diffusion models learn the *score* of the marginal distribution *p*(*x*_*t*_, *t*) of the diffusion process:

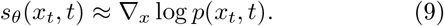

We can, however, analyze what this score means in terms of the energy formulation. If we also assume that the system of states *x*_*t*_ described by *p*(*x*_*t*_, *t*) at constant time *t* is at pseudo-equilibrium (in other words, the alchemical transformation described by the diffusion process is quasi-static, which is guaranteed if the diffusion process follows the Fokker-Planck Equation [52]), we can define an energy *E*_*t*_ for each stage *t* in the diffusion process (alchemical transformation) by generalizing eq. (7):

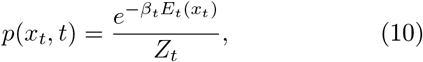

where *β*_*t*_ and *Z*_*t*_ are the inverse temperature and partition functions respectively at time *t*. Converting probability into energy for a physical system requires *β* and *Z* for the, physical units and reference energy, respectively. However, as *E*_*t*_ does not describe a physical system for *t >* 0, the values of *Z*_*t*_ and *β*_*t*_ are mostly arbitrary, and critically, our assumption of a particular temperature or partition function at time *t* does not affect our calculation of the energy at time 0 using *β* and *Z*_0_. Thus, for the purposes of this paper, we assume a constant *β*_*t*_ = *β* and *Z*_*t*_ = *Z*_0_ = *Z*:

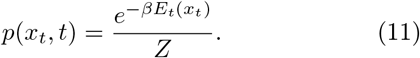

Taking the log of both sides isolates the energy term (with a coefficient of −*β*):

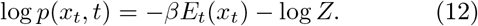

And since *Z* is independent of *x*_*t*_, taking the spatial gradient of both sides yields the score solely in terms of energy:

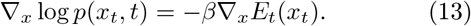

In classical physics, the negative gradient of energy with respect to position, −∇_*x*_*E*_*t*_(*x*_*t*_), is a measure of force, so we define it as *F*_*t*_(*x*_*t*_). Thus, output of a score-based diffusion model can be understood to be a scaled dimensionless force describing a learned energy function, *E*_*t*_(*x*_*t*_), at time *t* (see dimensions in **Table I**):

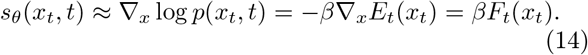

For samples generated by DFMDock, the data distribution *p*_0_ aims to capture the distribution of all protein-protein docking systems at equilibrium. *x*_0_ denotes a docked complex and 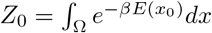 would then be the partition function of two associating proteins. For rigid docking between a receptor protein and a ligand protein, the search space Ω spans all possible translations and rotations of the ligand, with the receptor fixed. Thus, the negative log-likelihood of a docked sample generated at *t* = 0 is, in principle, proportional to the thermodynamic free energy of a protein-protein complex.

### C. Negative Log-Likelihood or “Learned Energy” Estimation

As score-based diffusion models do not produce *p*_0_ directly, we must find some way to recover it from the neural network’s output: the spatial gradient of the marginal probability distribution, ∇_*x*_ log *p*(*x*_*t*_, *t*). Following [11], we postulate that to recover *p*_0_(*x*_0_), one may integrate the change in marginal likelihood along a path from *x*_1_ in the known prior distribution *p*_1_ to the sample *x*_0_ in the unknown data distribution *p*_0_. We formulate this integral in two ways: first, we use the physical intuition of force and energy to try to construct a work integral over a path through data space *x*. Second, we use the probability theory behind the marginal distribution of our SDE to formulate an integral of 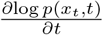 along a path from noise to data. We find that both approaches reduce to solving the same problem: integrating the spatiotemporal gradient field in both space and time of the marginal probability *p*(*x*_*t*_, *t*) over a path.

### D. Physics-based Energy Recovery: The Work Integral

In classical mechanics, calculation of the energy accumulated by an object acted on by an external force *F*_*t*_(*x*_*t*_) over a path *S* can be achieved using the work integral:

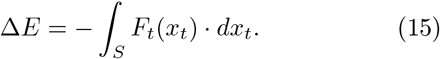

From eq. (14), *F*_*t*_(*x*_*t*_, *t*) can be replaced by 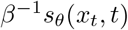 to measure the change in learned energy:

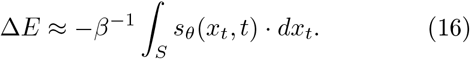

However, this integral is flawed, and leads to incorrect results both theoretically and numerically. To see why, we simply need to return to the derivation of the work integral:

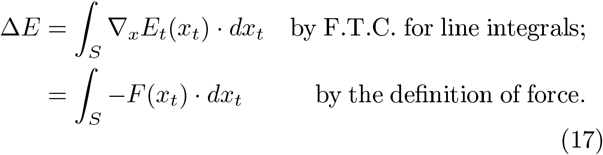

This derivation makes a fundamental assumption that the energy function, *E*_*t*_, is solely a function of space, *x*, and therefore independent of time, *t*. However, the connection between learned probability and learned energy gives no such guarantee, as the marginal probability *p*(*x*_*t*_, *t*) changes significantly with time. This corresponds to our understanding of the diffusion process as an alchemical transformation, during which the free energy of the system itself may change with alchemical time even when no work is done within the system. Thus, to compute ∆*E*, we need to incorporate both the spatial *and* temporal derivatives of *E*_*t*_:

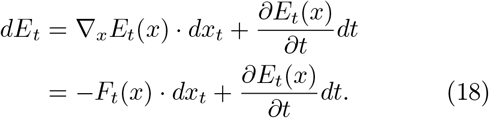

Along a given path *x*_*t*_, we can change variables by replacing *dx*_*t*_ with 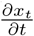*dt* to formulate ∆*E* for a space- and time-dependent *E*_*t*_(*x*_*t*_):

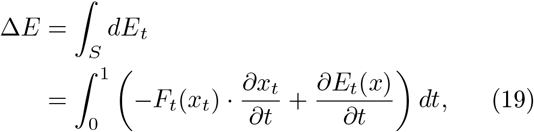

where we now define the integral range from 0 to 1 to correspond to the connection between the data distribution and the noise distribution.

This integral relies on both the spatial and temporal gradients of *E*_*t*_(*x*). While the learned spatial gradient is easily accessible from the score, how to determine the learned temporal energy gradient associated with the alchemical transformation is not obvious. To understand further, we need to examine the probability theory behind the marginal distribution *p*(*x*_*t*_, *t*)—so let’s briefly reframe the problem of learned energy recovery to that of likelihood computation.

### E. Negative Log-Likelihood Recovery

Our primary goal is to capture the energy of docked complexes *E*_0_(*x*_0_). Since we can directly relate the energy at *t* = 0 with the marginal log-likelihood log *p*_0_(*x*_0_) via eq. (12), if we can compute this log-likelihood directly, we can subsequently recover the energy up to a constant offset. Replacing *E* in eq. (19) with log *p* (and canceling the resulting *β*^*−*1^ terms) we can consider the line integral of the gradient of this log-likelihood over some path *x*_*t*_ through time and space:

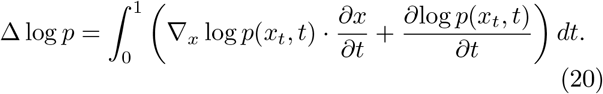

The learned score *s*_*θ*_(*x*_*t*_, *t*) can be used to approximate ∇_*x*_ log *p*(*x*_*t*_, *t*). The time-derivative 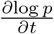can be addressed using the Fokker-Planck equation (see **Supplementary Material C**), which gives us the change in the marginal probability distribution of an SDE in terms of the diffusion coefficient *g*(*t*) and spatial derivatives of the distribution at time *t* (assuming the drift *f*(*x*_*t*_, *t*) = 0):

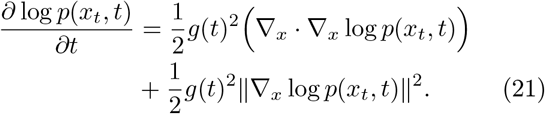

Now that we can write eq. (20) solely in terms of ∇_*x*_ log *p*(*x*_*t*_, *t*), we can once again substitute the learned score function *s*_*θ*_ from a trained diffusion model. Computing the divergence of the learned score requires taking its derivative; helpfully, modern neural network libraries (in the case of DFMDock, PyTorch) have built-in automatic differentiation capabilities used for backpropagation, so computing this derivative is tractable. Specifically, the divergence of the score is the sum of its partial derivatives along each dimension *i*, which can be rewritten more compactly as a trace:

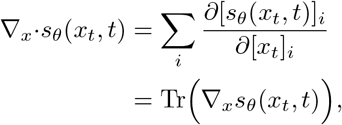

where 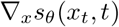 is the Jacobian matrix of *s*_*θ*_, whose diagonal elements are precisely the derivatives in the sum. Using this identity, we can rewrite eq. (21) in terms of *s*_*θ*_:

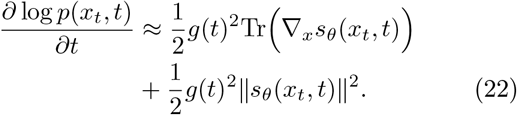

Plugging this approximation back into eq. (20), we have an expansion for the change in log probability over some path *x*_*t*_ in terms of the neural network score:

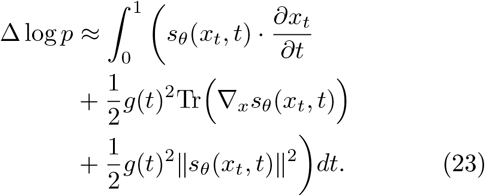

Returning to the energy formulation, eq. (12) tells us the energy field *E*_*t*_(*x*_*t*_) at time *t* is proportional to the log marginal probability, log *p*(*x*_*t*_, *t*), plus the time-independent constant *Z*, meaning its time derivative is simply

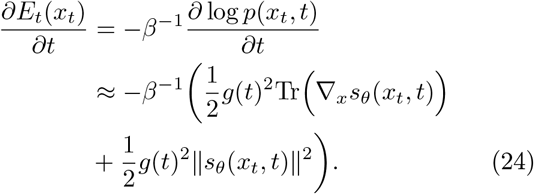

This completes the energy integral as well:

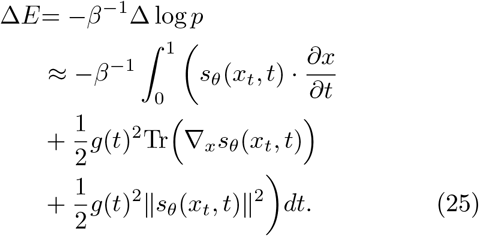

Using either the physical intuition or the underlying probability produces a line integral which evaluates the change in learned energy (log likelihood) over some path *x*_*t*_. Thus, if we want to compute the learned energy (log likelihood) of a point *x*_0_ in the unknown data distribution, it is sufficient to construct a path starting at *x*_0_ at *t* = 0 and ending at some point *x*_1_ at *t*_1_ in the known fully noised distribution *p*_1_. Then, using our integral, we can find the log likelihood of *x*_0_ as

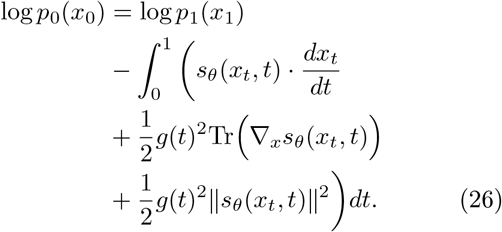

The negative sign on the integral stems from the fact that we are integrating from *t* = 0 (data) to *t* = 1 (noise), while the relevant delta is from noise to data. Later, we will be integrating over an ODE trajectory, where the data end-point is known but the noised endpoint is not, so we ensure consistency by always integrating from *t* = 0 to *t* = 1.

### F. DiffLikelihood: Extracting Log-Likelihood by Integrating Over Sampled Diffusion Paths

The most natural choice of path over which to evaluate our integral is the reverse diffusion trajectory used to generate a given sample, as calculated during model inference. These trajectories are computed by evaluating the reverse SDE equation (5) and using the diffusion model’s output *s*_*θ*_(*x*_*t*_, *t*) to approximate ∇_*x*_ log *p*(*x*_*t*_, *t*), resulting in a discrete trajectory of points in data-space 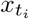 and diffusion time *t*_*i*_. For numerical accuracy while only sampling points on this discrete trajectory, we can use the trapezoid rule of numerical integration:

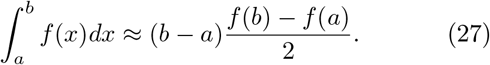

Applying this to each time step from 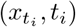 to 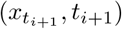 produces a discrete equation for log *p*_0_(*x*_0_)

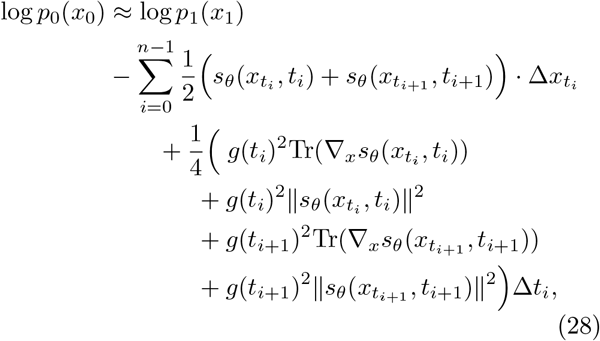

where 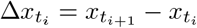 and 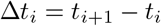.

The reverse diffusion path is convenient because it allows likelihood integration concurrent with inference by reusing the score computed by the network. Likewise, the trapezoid rule is the highest order numerical method that doesn’t require sampling additional intermediate points (and thus additional evaluations of the score model). Alternatively, trapezoid integration can be interpreted as a discrete implementation of the Stratonovich integral of our likelihood integrand, which is required instead of standard Itô integration to enable the use of the chain rule, and thus eq. (20) [53]. In either case, by using the trapezoid rule once per pair of vertices, we effectively approximate the integral of a piecewise-differentiable path of line segments. The pseudocode algorithm for trapezoidal integration of diffusion trajectories (**Supp. Algorithm 3**) can be found in **Supplementary Material E**.

If need be, one could integrate each segment more accurately by further interpolating to sample the score at more points along the trajectory (**Supp. Algorithm 4**), or by using a black-box ODE integrator to intelligently sample along each piecewise-differentiable segment (**Supp. Algorithm 5**). We find that this extra precision does not significantly change the results when used to integrate our Gaussian toy model’s diffusion trajectories compared to trapezoidal integration (**Supp. Fig. S1**). However, piecewise ODE integration of DFMDock diffusion trajectories meaningfully decreases noise (**Supp. Fig. S2**), so we use the higher precision method for DFMDock trajectories. A detailed breakdown and comparison of diffusion trajectory integration method performance can be found in **Supplementary Material F**.

Although we are able to achieve good numerical precision on diffusion trajectories, we will see the noise term in eq. (5) produces paths with large displacements ∆*x*_*t*_ (**Fig. 2a**) and noisy likelihoods (**Fig. 3a**), motivating us to explore integration over other paths.

**FIG. 2.**
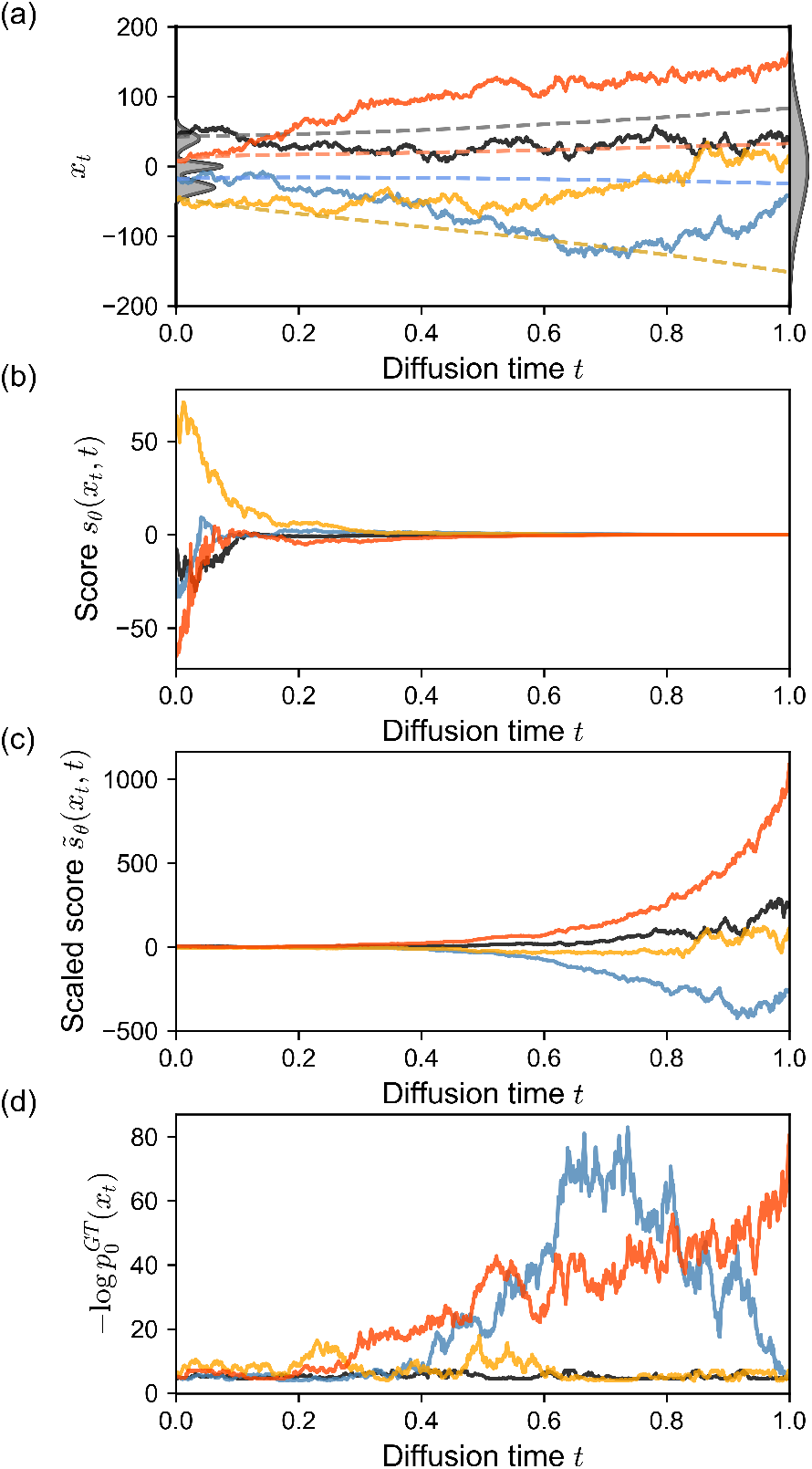
Trajectories for four individual 1D samples. (a) Diffusion trajectories *x*_*t*_ from the Gaussian mixture diffusion model. Flow paths that result in the same ending points as the diffusion paths are shown as dashed lines. Ground-truth distribution (left) and prior distribution (right) shown in gray. (b) The learned score function *s*_*θ*_(*x*_*t*_, *t*) = *∇*_*x*_ log *p*(*x*_*t*_, *t*). (c) The scaled score function 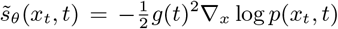 with the exponential noise schedule 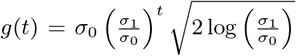 (d) Ground truth energy 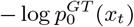 evaluated over 1000 diffusion steps from *t* = 1 to *t* = 0.

**FIG. 3.**
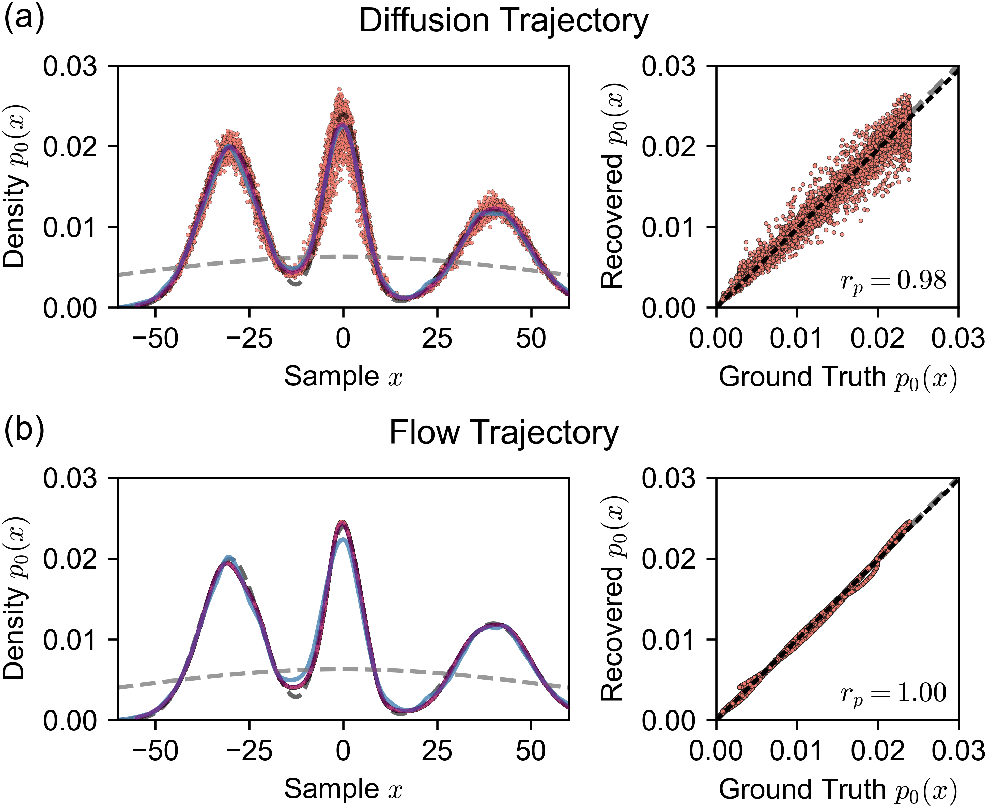
Recovered probabilities for samples generated from a 1D Gaussian mixture diffusion model integrated over (a) diffusion trajectories and (b) flow trajectories (red dots, left panels), computed as *p*_0_(*x*_0_) = exp(log *p*_1_(*x*_1_)− ∆ log *p*). For the diffusion trajectories in (a), the binned mean of recovered probabilities is shown as a purple curve. The dashed black curve represents the data distribution 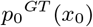,the dashed gray curve shows the unimodal Gaussian prior distribution *p*_1_(*x*_1_), and the blue curve is a kernel density estimate of samples (*N* = 10, 000) generated from the diffusion model. The right panels show correlation plots of recovered probability values from both trajectories versus the ground truth probability values. A *y* = *x* line is shown in gray dashes, and a dotted linear model is shown in black with Pearson coefficients *r*_*p*_.

### G. FlowLikelihood: Extracting Log-Likelihood by Integrating over Flow-Equivalent ODEs

Since log *p*(*x*_*t*_, *t*) is a state function, we are free to integrate over alternative paths to the same sample *x*_0_ to compute log *p*_0_(*x*_0_). A less noisy alternative to the diffusion trajectories is to use a flow-equivalent ODE as described by [11], where paths are generated from the following ODE:

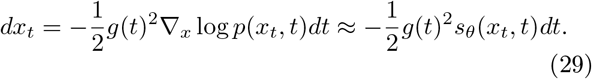

The likelihood integral in eq. (26) is then evaluated over these flow trajectories from *t* = 0 to *t* = 1 with initial condition *x*_0_, the point in the data distribution whose learned likelihood we seek.

The explicit description of the path in terms of an ODE confers two major benefits over a discretely sampled diffusion trajectory: First, both path and integrand can be computed using existing black-box ODE solvers, rather than the limited discrete methods used for the diffusion trajectories. Second, since we have an explicit form for *dx*_*t*_, we can substitute it directly into our integrand’s 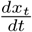 term. When we do so, the first and third terms cancel (see **Supplementary Material D**), leaving us with the much simplified:

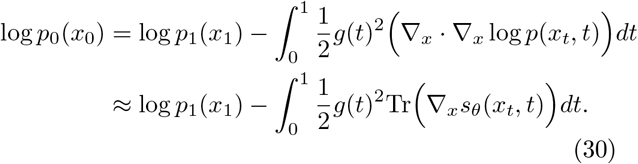

This simplification is no coincidence; when initialized from the same prior distribution at *t* = 1, the probability flow ODE shares the same marginal distribution as the forward and reverse diffusion SDEs in eqs. (3) and (5) (and is approximated using the same score as used for diffusion inference), which means it also functions as a continuous normalizing flow (CNF) model [11, 47]. As CNF trajectories smoothly follow the probability flow of their own marginal, they have a natural formula for computing the change in likelihood along an ODE solution: the instantaneous change-of-variables formula, simply the divergence of the underlying vector field, which tracks the dispersion or concentration of probability along the path [47].

Integrating over the flow trajectory simplifies the integral and provides a smoother, easier to integrate path than the noisy, discrete diffusion trajectory. Unlike diffusion trajectory integration, since a flow trajectory can be computed for any point in the data space, we are no longer limited to assessing the likelihood of points generated during inference. However, flow integration comes at the computational cost of having to generate a path *de novo* for each sample. Pseudocode for integration over flow trajectories can be found in **Supplementary Material E, Supp. Algorithm 6**.

## IV. Results

### A. Temporal Analysis of Diffusion Trajectories from 1D Diffusion Model

To validate our theoretical analysis, we trained a 1D diffusion model to learn a simple Gaussian mixture distribution (see **Section VI.A** for training procedure), generated samples from the trained model, and evaluated the recovery of learned energy over both reverse diffusion and flow trajectories.

**Fig. 2a** shows four sampled reverse diffusion trajectories from the toy model. The diffusion trajectories from the model are noisy at early times (*t* = 1) and become progressively smoother as the model gradually transforms the points to the data distribution (*t* →0) and the amount of added noise per diffusion step decreases. By solving the probability flow ODE (eq. 29), we also compute and show the flow paths that result in the same sample point at *t* = 0 (dashed lines). **Fig. 2b** shows the learned score function *s*_*θ*_(*x*_*t*_, *t*) = ∇_*x*_ log *p*(*x*_*t*_, *t*) along those four diffusion trajectories. The score has low magnitude near *t* = 1 which indicates low confidence of steering towards the high-likelihood regions of the data. As time decreases, the magnitude of the score increases, and the model shows stronger guidance for *x*_*t*_ towards high-likelihood regions of the data distribution.

In **Fig. 2c**, we visualize the scaled score-term used in the reverse SDE, 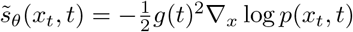, where *g*(*t*) is the noise-schedule (diffusion coefficient). As we use an exponential noise schedule, which exhibits strong weighting, 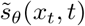shows large fluctuations near *t* = 1, and its magnitude decays as *t* → 0. To better interpret how the system evolves from noise to data, we use the ground truth distribution of the trimodal Gaussian to plot 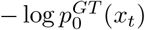 evaluated over the 1000 reverse diffusion steps (**Fig. 2d**). The ground truth energies are high and noisy at early times, but the fluctuations gradually decay and converge to a stable value as *x*_*t*_ approaches the high-likelihood regions of the data distribution *p*_0_(*x*_0_).

### B. Integration Over Diffusion and Flow Trajectories Recover Probabilities that Approximate the True Data Distribution

In **Fig. 3a-b**, we plot the 1D system ground truth (training) distribution *p*_0_(*x*_0_) (dashed black curve) and the prior distribution *p*_1_(*x*_1_) (dashed gray curve). To check the 1D diffusion model’s ability to generate samples from the data distribution, we plot the kernel density estimate (KDE) of 10,000 sample data points generated from the model (blue curve), showing that the model has learned the ground truth distribution well (Kolmogorov-Smirnov distance of 0.011).

Next, we examined the energies recovered from our integral formulations. We computed negative log-likelihood values (NLL) of samples generated by our Gaussian mixture diffusion model by integrating over both diffusion and flow trajectories. To evaluate the quality of the NLL values, we plot the recovered probabilities *p*_0_(*x*_0_)= exp(log *p*_1_(*x*_1_) −∆ log *p*) (red dots in **Fig. 3a-b**). The distribution recovered by integrating over diffusion trajectories shows noisy deviation from the training data distribution, but the binned average probability (purple line) matches the ground-truth training distribution almost exactly. In contrast, integrating over flow trajectories yields a noise-free smooth curve that closely follows the ground-truth distribution.

To quantify the consistency between the integrals, **Fig. 3** shows scatter plots of the recovered probability versus ground truth probability for the three integral methods (right panels). The Pearson correlation coefficients (*r*_*p*_ = 0.98 for diffusion and *r*_*p*_ = 1.00 for flow) indicate that learned likelihoods from both diffusion and flow trajectories approximate the underlying data distribution very well.

### C. Temporal Analysis of Protein Docking Diffusion Trajectories from DFMDock

Following the 1D diffusion model analysis, we now apply the same approaches to a 3D, rigid-body, translational diffusion model for protein docking, where the diffusion process occurs over *x*_*t*_ = (*x, y, z*) in 3D Euclidean space. We performed our analysis on the DFMDock model [21], which learns to dock protein complexes given two unbound protein monomers. We limited the docking space to translation only (see **Section VI.B**), as the version of the Fokker-Planck equation used in eq. (21) assumes R_*n*_ Euclidean space and is incompatible with general Riemannian manifolds like the SE(3) space used in rotational diffusion. We performed DFMDock inference on 25 targets from Docking Benchmark 5.5 (DB5.5) [54, 55] with 120 samples for each target.

We first examined the score and the learned energies over the 40-step reverse diffusion trajectories starting from the unbound monomers at *t* = 1 to the bound complex at *t* = 0 for two example targets. As an example of a complex for which DFMDock generated accurate samples, we selected the complex of subtilisin BPN’ with an inhibitor (PDB: 2SIC). DFMDock-generated samples of docked 2SIC structures exhibited DockQ scores of up to 0.96 (high quality). As an example of a complex for which DFMDock generated diverse but low-accuracy structures, we chose PKR kinase domain-eIF2*α* (PDB: 2A1A); the best generated docked 2A1A structure only reached a DockQ score of 0.72, a medium quality ranking. Two denoising trajectories from *t* = 1 to *t* = 0 representing the most successful docking attempts for each target are shown in **Fig. 4a,e**. Similar to 1D cases, we observe increasing score *s*_*θ*_(*x*_*t*_, *t*) (**Fig. 4b,f**) and decreasing scaled score 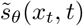 (**Fig. 4c,g**) moving towards *t* = 0.

**FIG. 4.**
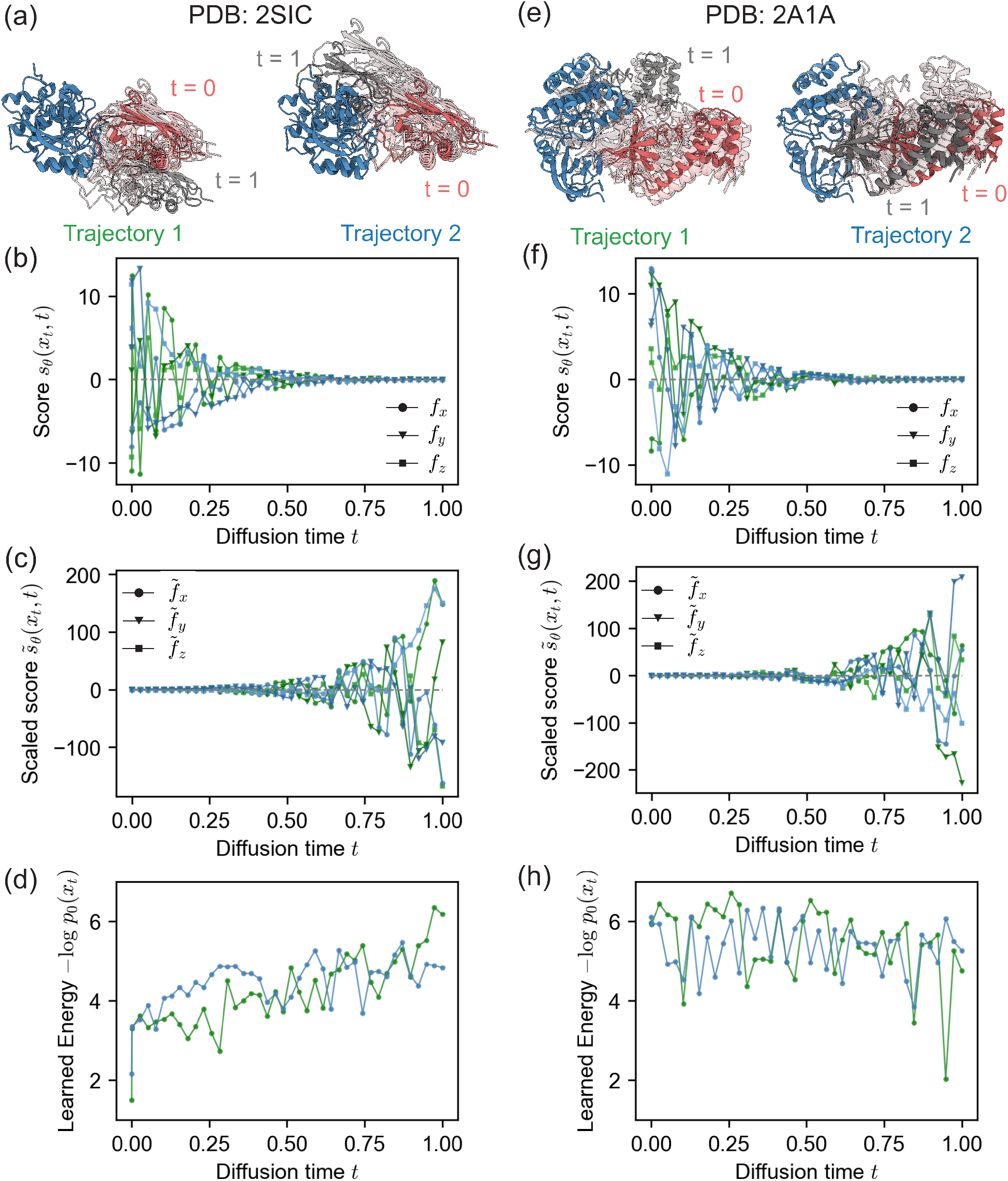
DFMDock reverse diffusion trajectories for two successful docking samples of two protein complexes, (a-d) 2SIC and (e-h) 2A1A. (a,e) Top two DockQ structures. Receptor protein, blue; ligand protein, gray (*t* = 1) to red (*t* = 0). (b,f) Score function learned by DFMDock *s*_*θ*_(*x*_*t*_, *t*) = *∇*_*x*_ log *p*(*x*_*t*_, *t*) along *x, y*, and *z* directions (*f*_*x*_,*f*_*y*_,*f*_*z*_). (c,g) Scaled score function 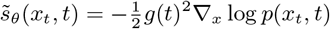 with 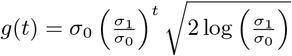 along *x, y*, and *z* directions 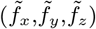. (d,h) Learned energy, *−* log *p*_0_(*x*_*t*_), recovered from integrating flow trajectories terminating at each point along the reverse diffusion trajectory.

As flow trajectories can be used to retrieve the energy of a pose without needing its diffusion trajectory, we used them to compute the learned energy, − log *p*_0_(*x*_*t*_), of intermediate poses *x*_*t*_ in the learned data distribution *p*_0_ (**Fig. 4d,f**). For the 2SIC trajectories, the learned energy shows a general downward trend over the course of 40 diffusion steps, suggesting that the reverse diffusion process increasingly favored more energetically plausible protein-protein interactions learned by the model. This decrease in energy can be correlated with improved docking quality as the model samples more refined conformations. The 2A1A trajectories, on the other hand, display deep and sharp oscillations and few general trends as the diffusion time approaches zero, notably *not* decreasing as time goes to zero.

### D. Learned Energies Compare to Rosetta Energy for Scoring Protein-Protein Interactions

We then computed the learned energy −log *p*_0_(*x*_0_) of 120 DFMDock generated structures for all 25 protein-protein complexes. As in the 1D case, we extracted potentials for each sample by integrating eqs. (26) and (30) over diffusion and flow trajectories respectively (**Supp. Fig. S4-S11**); we used multiple numerical integration strategies for diffusion trajectories, namely trapezoidal integration (**Supp. Algorithm 3, Supp. Fig. S4, S5**), interpolated trapezoidal integration (**Supp. Algorithm 4, Supp. Fig. S6, S7**), and piecewise ODE integration (**Supp. Algorithm 5, Supp. Fig. Supp. Fig. S8, S9**). Our experiments suggest that piecewise ODE integrated likelihoods are the most accurate (**Supplementary Material F**), so here we focus on diffusion likelihoods integrated by piecewise ODE. To analyze whether these potentials are low for near-native structures, we used two measures of similarity to the ground truth structures, interface-residue RMSD (**Supp. Fig. S4, S6, S8, S10**) and DockQ, a composite measure for docking quality [56] (**Supp. Fig. S5, S7, S9, S11**). In **Fig. 5**, we examine the learned and Rosetta energies for the same two complexes analyzed in **Section IV.C**, 2SIC and 2A1A. For both 2SIC and 2A1A, the Rosetta energy reveals a funnel-like curve with the global minimum near the ground truth (native) structure. For 2SIC, the learned energies from both flow and diffusion also display a near-native global minimum (**Fig. 5a**, rows 1 and 2). DFMDock’s 2SIC energy landscape seems to have a sharp funnel centered near the native structure, with a relatively flat distribution outside of it. That is, many samples lie at approximately the same non-minimal energy (*βE*_0_ ≈4 for flow, 0 for diffusion), including several samples ranked well by DockQ or interface RMSD. The inability to distinguish between medium- and low-quality structures might indicate that DFMDock has learned little about physical long-range interactions that might guide a protein towards its docked conformation, instead learning that structures without proper short-range interactions are unlikely to be native. In the case of 2A1A, the DFMDock learned energies fail to capture the correct binding energetics, with learned energies from flow integration presenting what may be a non-native energy funnel consisting of incorrect structures (iRMSD≈ 20, DockQ ≈0). Interestingly, learned energies from diffusion do not show this minimum, which might be due to the greater variance of integrating stochastic energies or could reflect a consistent bias along paths produced by flow integration. In any case, both methods rank near-native structures with similar or higher energies as non-native, while Rosetta energies correctly rank near-native poses best. The contrast between DFMDock’s ability to sample high-quality poses and its inability to properly rank poses with its energy for certain targets indicates that the DFMDock learned energy does not reflect the physical principles of protein-protein interactions in all cases.

**FIG. 5.**
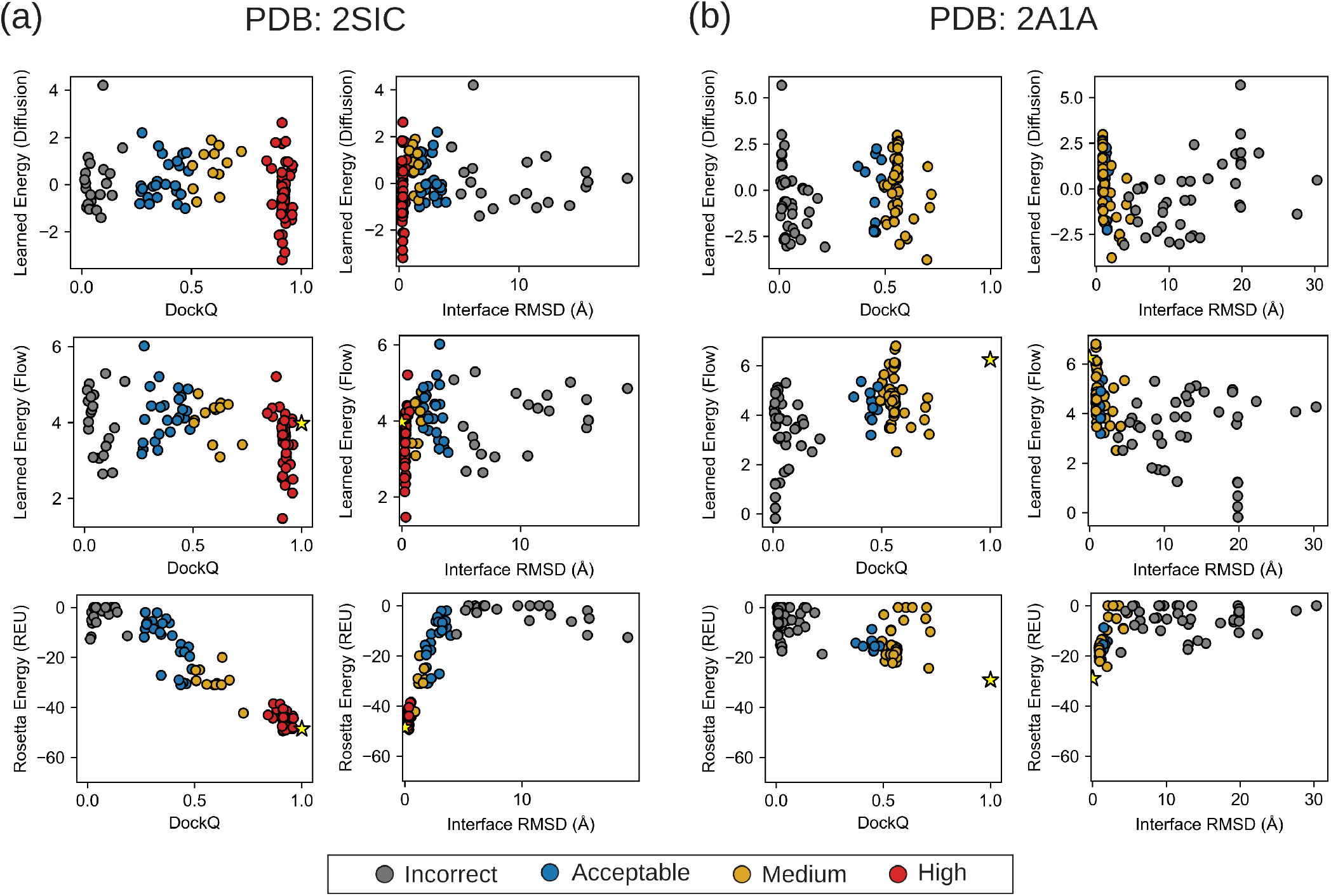
Energy vs Docking accuracy for 120 DFMDock-generated docking poses for each of (a) PDB ID: 2SIC and (b) PDB ID: 2A1A. Energies used are (1st row) learned energy from integrating over DFMDock diffusion trajectories by piecewise ODE integration, (2nd row) learned energy from integrating DFMDock flow trajectories, and (3rd row) Rosetta interface energy. Energies are plotted against DockQ score (1st and 3rd columns) and interface RMSD (2nd and 4th columns). Individual points are colored by the quality of the corresponding docking pose based on the CAPRI classification [57]: incorrect, gray; acceptable, blue; medium quality, gold; high quality, red. Energies computed for the ground truth structures are shown as yellow stars. As there is no diffusion trajectory for ground truth structures, no star is plotted for learned diffusion energies.

With flow integration, we can also compute the DFMDock learned energy for ground-truth poses without a diffusion trajectory. Learned energies of ground truth poses (**Fig. 5, Supp. Fig. S10, S11**, yellow stars) are often higher than those of the best DFMDock-generated docking pose, and fall within two standard deviations of lower quality ones. The high learned energies of ground truth poses suggest that DFMDock has not sufficiently learned to capture protein-protein interactions.

Finally, we explored whether the funnel-like behavior of the DFMDock learned energies is useful for ranking docking poses. **Fig. 6a** compares the top ranked model quality (DockQ) for the 25 protein-protein complexes with learned energy from flow integration (blue), learned energy from diffusion integration (green), Rosetta energy (red cross), and an oracle setting (black star, the sampled pose with the highest DockQ score). The learned energy from flow integration performs comparably or outperforms Rosetta interface energy in 9 out of 25 cases in identifying correct docking poses among the sampled poses. Diffusion-trajectory learned energy performs similarly, matching or outperforming Rosetta on 8 structures, but surprisingly performs well on *different* structures than flow. Combined, flow and diffusion energies together rank 13 structures as well as Rosetta ranking. Examining the per-sample energies (**Supp. Fig. S8-S11**), it is unclear how much of the success of diffusion-trajectory energy ranking of lower quality structures is purely due to random noise. Nonetheless, the result implies an orthogonality and potential synergy between diffusion and flow, as well as the value of exploring multiple paths through diffusion space to the same sample, which we elaborate on in **Section V**.

**FIG. 6.**
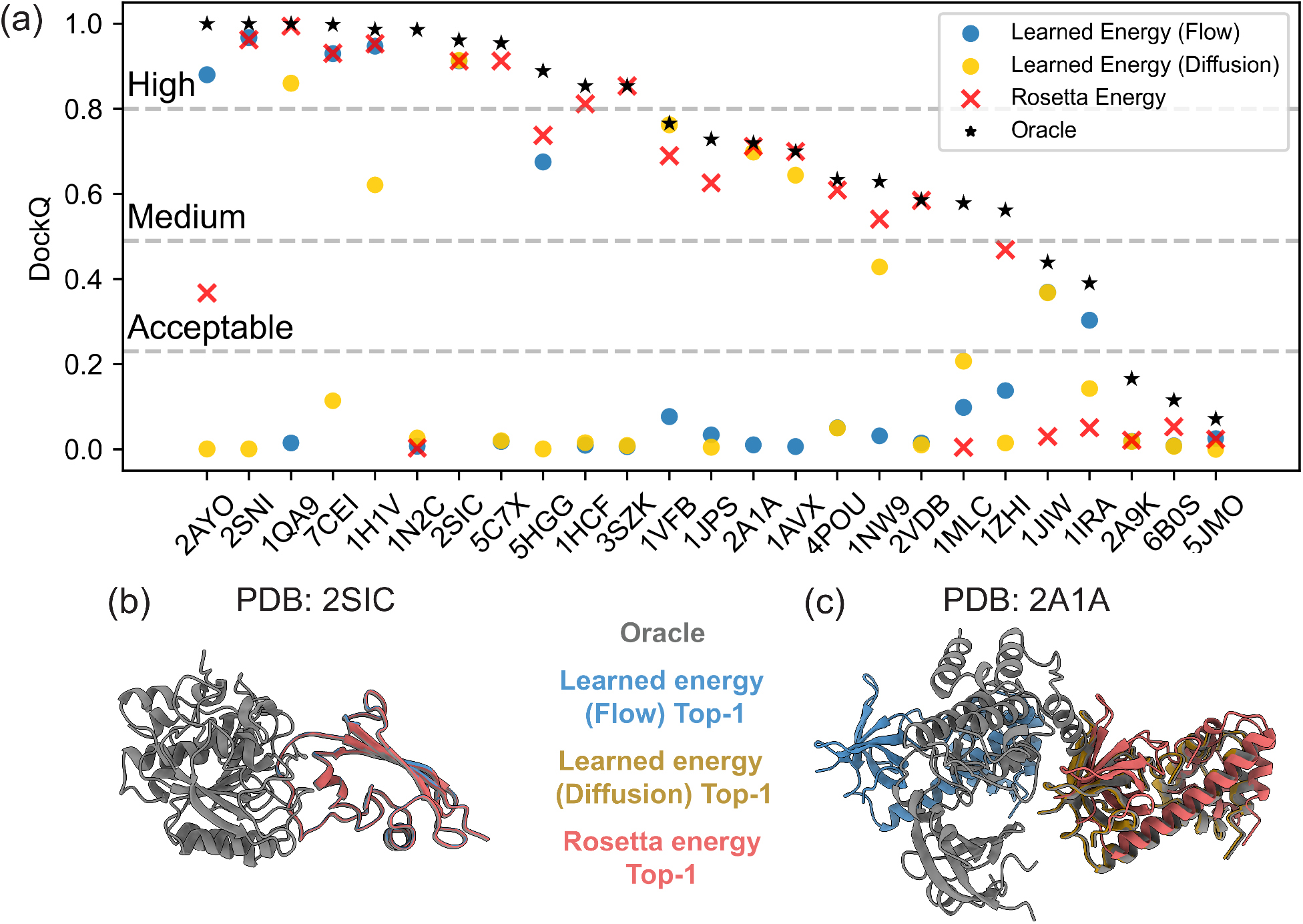
(a) Comparison of top-ranked model quality (DockQ) for 25 targets from the DB5.5 test set with learned energy over flow trajectories (blue circles), diffusion trajectories (gold circles), Rosetta energy (red crosses), or in the oracle setting (black stars). Top predictions ranked by the learned energy over flow trajectories (blue), diffusion trajectories (gold), and Rosetta energy (red) are compared against the highest-DockQ generated structure (gray) for (b) 2SIC (Oracle DockQ: 0.96) and (c) 2A1A (Oracle DockQ: 0.72).

**Fig. 6b,c** shows a structural comparison between the learned energy and Rosetta interface energy’s top-ranked predictions for two targets. For 2SIC, both of the learned energies (flow and diffusion) and Rosetta interface energies identify (distinct) high quality poses, with DockQ = 0.91 in all three cases. However, for 2A1A, the learned energy fails to identify even an acceptable quality structure (DockQ = 0.01). Interestingly, diffusion-integrated learned energy identifies a medium quality pose (DockQ = 0.69) nearly matching the docking quality of Rosetta (DockQ = 0.71).

## V. Discussion and Conclusion

The goal of this work was to interpret diffusion models through the lens of statistical thermodynamics by analyzing the underlying learned potential function (at *t* = 0) and exploring its applications in scoring protein complexes. We developed theory rooted in statistical thermodynamics to relate the probability of observing a system in a particular state to the energy of that state. We defined the energy function that a diffusion model implicitly learns as the *learned energy* and showed that it is equal to the negative log-likelihood, −log *p*_0_(*x*_0_), up to a constant reference energy log *Z* and scaled by an inverse temperature *β*.

Since diffusion models do not explicitly learn the data distribution *p*_0_(*x*_0_), but rather the score function, ∇_*x*_ log *p*(*x*_*t*_, *t*), we developed methods to evaluate *p*_0_(*x*_0_). We postulated that to recover *p*_0_(*x*_0_), we can integrate along a path from the known prior distribution *p*_1_(*x*_1_) to the target data distribution *p*_0_(*x*_0_). We constructed a path integral over *x*_*t*_ to calculate the likelihood, and evaluated it over two paths: (1) discretized diffusion trajectories from the reverse SDE, and (2) flow trajectories, which follow a smooth, deterministic path defined by an ODE equivalent in marginal probability to the reverse SDE.

We initially tested our approach on a simple 1D diffusion model of a Gaussian mixture distribution. Using our integral methods, we evaluated −log *p*_0_(*x*_0_) and *p*_0_(*x*_0_) values of samples generated by the model and compared them to the known analytical solution. While diffusion trajectory integrals are convenient since they can be calculated from model inference points, their integration results in noisy likelihood recovery, even as averaged diffusion likelihoods well approximate the ground truth distribution. Fortunately, flow trajectories are both smooth and integrable using black-box ODE solvers. Flow trajectories effectively and continuously approximate the probabilities of the true data distribution at the cost of an additional path calculation.

Building on these results, we applied the same methodology to a protein-protein diffusion docking model, DFMDock, which generates protein complex structures given two unbound protein monomer structures. We extracted the learned potentials from DFMDock to score protein complexes and compared them to Rosetta energies. The learned energies from DFMDock-generated docking poses reveal binding energy funnels that sometimes match Rosetta’s interface energy funnels in that the near-native structures have lower energies. But in other cases, the learned energy funnels have minima at non-native structures, and the native structures have higher energies. This work shows that we can examine ensembles of diffusion-generated protein structures in a similar manner as in physics-based energy approaches.

Our method for likelihood calculation integrates the spatiotemporal gradient of the marginal probability *p*(*x*_*t*_, *t*) over two paths: the probability flow ODE, and a discrete sampling of the reverse-time SDE. Interestingly, this gradient is (by definition) a conservative vector field over the combined diffusion space ℝ^*n*+1^ (combining data-space *x*_*t*_∈ ℝ_*n*_ and time *t* ∈ℝ), which means that the integral of this gradient, ∆ log *p*, should be dependent solely on the endpoints of the path, not on the trajectory we use for integration. Since we have an explicit formula for log *p*_1_, this means the calculated value of log *p*_0_ = log *p*_1_ −∆ log *p* should be the same for any path we use during integration.

In practice, however, we find this is not the case: flow trajectories do not yield the same value as diffusion trajectories, and different diffusion paths terminating at near-identical samples can have significantly different likelihoods. This suggests that one or more of our assumptions must not hold in practice, leading us to identify two main sources of error: 1) inaccuracy in our explicit formula for the prior, and 2) spatiotemporal non-conservativity of the learned score.

First, a critical assumption of diffusion models is that the marginal distribution at time *t* = 1 is a Gaussian with known mean and variance. In practice, reaching a true Gaussian requires adding infinite noise and accurately estimating the mean and variance of the data distribution. As the former is impossible and the latter is difficult, especially for low-data conditional diffusion like protein docking, the assumed prior will always deviate from the true *t* = 1 marginal, introducing error. In fact, diffusion models in the image generation space have been shown to have decidedly inaccurate priors [58, 59]; thus, it is likely to be a concern for likelihood recovery even for well-performing models like AlphaFold3. We explored the effects of this error in **Supplementary Material B**, and used the experiments to select optimal hyperparameters for our 1D toy model. Interestingly, while prior inaccuracy introduces bias to deterministic flow integration, its effect on diffusion trajectory integration is to add significant noise while maintaining the accuracy of the average of stochastic trajectories terminating at the same sample. Averaging the learned energy integrated along multiple paths to the same *x*_0_ is similar in principle to the ensembling techniques used by He *et al*. in their FEAT method [45]; formalizing this approach may yield better results than either flow or diffusion trajectories.

Second, there is evidence that diffusion models do not generally learn spatially-conservative [60] or Fokker-Planck consistent [61] scores, introducing further path-dependence. Spatial and temporal conservativity are likely to vary across diffusion space and time, meaning certain paths from noise to data might yield more accurate reflections of the underlying marginal probability than others. The regions with highest marginal probability are sampled more during both training and inference, potentially justifying the use of SDE/ODE paths whose marginals match that of the diffusion process. Still, there may be benefits to engineering alternative paths. For example, as inference can follow many possible paths through diffusion space, using an ensemble as described above might yield a more representative sample of inference time likelihoods than any single path. Further insight may be gained by exploring the conservativity in the score learned by networks like DFMDock and AlphaFold3 and the impact of score inaccuracies on different trajectories and methods of likelihood computation. An open question in the field of protein structure prediction and design is whether and how AI models might learn a thermodynamic function of protein folding and association. Better interpretability of AI models in biology is also important from a biosecurity perspective. There are other investigations of these questions in the literature. Ahdritz *et al*. [62] released OpenFold, an open-source version of AF2 and carefully investigated model’s learning process and the model’s capacity to generalize to unseen regions of fold space. Roney *et al*. [63] hypothesized that AF2 has learned an implicit energy function through its confidence module, and demonstrated that AF2 can be used to rank the quality of candidate protein structures without needing coevolution data.

While we explored here how physical energies can be accessed from learned diffusion models, another approach is to inject biophysical knowledge into AI models. Kulyte *et al*. [64] used molecular dynamics (MD) force-fields to guide diffusion models for antibody design. Similarly, Wang *et al*. [65] used MD force-fields to guide diffusion models for protein conformation generation. Lewis *et al*. [66] trained a diffusion model on MD conformational ensembles to improve realistic diversity of conformational sampling. Analyses like ours could help probe the effects of adding physical information into deep-learning based biomolecular design and structure prediction tools.

To our knowledge, this work is the first attempt at extracting potentials from a biomolecular score-based diffusion model and comparing them with physical energy functions. For feasibility, we restricted ourselves to a low-dimensional rigid body docking model and tested on a small validation set as a proof of concept. More work is needed to interrogate the scalability of the method and our findings to more complex models and larger, more diverse protein datasets. An exciting future direction is to apply our methodology to state-of-the-art diffusion-based structure prediction models such as AlphaFold3 [7] or Boltz-2 [14] to interrogate their learned energy functions associated with protein folding stability. Another important direction is to generalize likelihood calculation to be compatible with diffusion models over arbitrary Riemannian manifolds such as the SE(3) space used in the original DFMDock with rotations [21] and DiffDock-PP [27]. As we are not limited only to computing the likelihood of generated samples, exploring the learned energy landscape around generated structures might reveal important details about the shape of the learned energy funnel. For example, whether the structure is at a local minimum could be used as a ranking metric, and learned energy surfaces could be used for local refinement.

## VI. Data

### A. Gaussian Mixture Model Training

To train a 1D diffusion model to learn a simple Gaussian mixture distribution, we generated 60,000 training data points from a trimodal Gaussian, defined as a weighted sum of three individual Gaussian components. The probability density function is

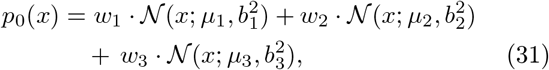

where the means are *µ*_1_ = −30, *µ*_2_ = 0, and *µ*_3_ = 40; the standard deviations are *b*_1_ = 8.0, *b*_2_ = 5.0, and *b*_3_ = 10.0; and the mixture weights are *w*_1_ = 0.4, *w*_2_ = 0.3, and *w*_3_ = 0.3. For testing, we sampled 10,000 synthetic points with 1,000 diffusion steps from the learned model and compared the probability density function of the analytical formulation with the kernel density estimate of the generated samples (**Fig. 3**).

### B. DFMDock Inference

We performed inference using the DFMDock model [21], which is trained on DIPS-hetero [67], a subset of DIPS with approximately 11,000 heterodimers. Although DFMDock was originally designed to reverse both translational and rotational noise, here we restricted the added rigid-body noise during both training and inference to be translation-only, such that all processes occur in the 3D Euclidean space R_3_. Additionally, the original formulation of DFMDock uses a random sampling method to construct the interaction graph as input to the neural network, whereas here the graph is generated deterministically using k-nearest neighbors. This ensures that all stochasticity during inference comes solely from added perturbations of the diffusion process. Using the same training parameters as [21], we retrained the model with these modifications and observed slightly degraded performance. As in [21], we use Docking Benchmark 5.5 (DB5.5) [54, 55], a widely used dataset for assessing docking performance, and sampled 120 poses with 40 diffusion steps for each of 25 targets from the set. For Rosetta energies computed for the sampled poses, we first performed the `docking_local_refine` protocol before computing the *I*_*sc*_ score with the REF15 energy function.

### C. ODE Integration

Black-box ODE integration was performed using the TorchDiffeq library, https://github.com/rtqichen/torchdiffeq [68]. For both flow and piecewise ODE trajectories, we use the ‘RK4’ method (fourth-order Runge-Kutta with 3/8 rule) with a step size equal to that of diffusion inference; our Gaussian 1D trajectories are sampled with 100 diffusion steps, so the ODE step size is 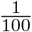,and DFMDock is sampled with 40 diffusion steps, so the ODE step size is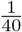. We explore the tradeoffs between precision and computation time of smaller DFMDock step sizes in **Supplementary Material F**.

## Code Availability

The training, inference, and likelihood evaluation code for the 1D diffusion model and the likelihood evaluation code for DFMDock are available at https://github.com/Graylab/DiffEnergy.

## Acknowledgments

This work was supported by National Institutes of Health grant R35-GM141881, Moderna, and AstraZeneca. Computational resources were provided by the Advanced Research Computing at Hopkins (ARCH). The authors thank Jeremias Sulam and Jacopo Teneggi for suggesting that we explore an analytic, 1D model to gain better insight into our DockQ values.

## References

[1] A. Ramesh, M. Pavlov, G. Goh, S. Gray, C. Voss, A. Radford, M. Chen, and I. Sutskever, Zero-Shot Text-to-Image Generation (2021), 2102.12092 [cs].

[2] R. Rombach, A. Blattmann, D. Lorenz, P. Esser, and B. Ommer, High-Resolution Image Synthesis with Latent Diffusion Models (2022), 2112.10752 [cs].

[3] A. Borji, Generated Faces in the Wild: Quantitative Comparison of Stable Diffusion, Midjourney and DALL-E 2 (2023), 2210.00586 [cs].

[4] Y. Liu, K. Zhang, Y. Li, Z. Yan, C. Gao, R. Chen, Z. Yuan, Y. Huang, H. Sun, J. Gao, L. He, and L. Sun, Sora: A Review on Background, Technology, Limitations, and Opportunities of Large Vision Models (2024), 2402.17177 [cs].

[5] J. L. Watson, D. Juergens, N. R. Bennett, B. L. Trippe, J. Yim, H. E. Eisenach, W. Ahern, A. J. Borst, R. J. Ragotte, L. F. Milles, B. I. M. Wicky, N. Hanikel, S. J. Pellock, A. Courbet, W. Sheffler, J. Wang, P. Venkatesh Sappington, S. V. Torres, A. Lauko, V. De Bortoli, E. Mathieu, S. Ovchinnikov, R. Barzilay, T. S. Jaakkola, F. DiMaio, M. Baek, and D. Baker, De novo design of protein structure and function with rfdiffusion, Nature 620, 10.1038/s41586-023-06415-8 (2023).

[6] J. B. Ingraham, M. Baranov, Z. Costello, K. W. Barber, W. Wang, A. Ismail, V. Frappier, D. M. Lord, C. Ng-Thow-Hing, E. R. Van Vlack, S. Tie, V. Xue, S. C. Cowles, A. Leung, J. V. Rodrigues, C. L. Morales-Perez, A. M. Ayoub, R. Green, K. Puentes, F. Oplinger, N. V. Panwar, F. Obermeyer, A. R. Root, A. L. Beam, F. J. Poelwijk, and G. Grigoryan, Illuminating protein space with a programmable generative model, Nature 623, 1070–1078 (2023).

[7] J. Abramson, J. Adler, J. Dunger, R. Evans, T. Green, A. Pritzel, O. Ronneberger, L. Willmore, A. J. Ballard, J. Bambrick, S. W. Bodenstein, D. A. Evans, C.-C. Hung, M. O’Neill, D. Reiman, K. Tunyasuvunakool, Z. Wu, A. Žemgulytė, E. Arvaniti, C. Beattie, O. Bertolli, A. Bridgland, A. Cherepanov, M. Congreve, A. I. Cowen-Rivers, A. Cowie, M. Figurnov, F. B. Fuchs, H. Gladman, R. Jain, Y. A. Khan, C. M. R. Low, K. Perlin, A. Potapenko, P. Savy, S. Singh, A. Stecula, A. Thillaisundaram, C. Tong, S. Yakneen, E. D. Zhong, M. Zielinski, A. Žídek, V. Bapst, P. Kohli, M. Jaderberg, D. Hassabis, and J. M. Jumper, Accurate structure prediction of biomolecular interactions with alphafold 3, Nature 630, 10.1038/s41586-024-07487-w (2024).

[8] J. Boitreaud, J. Dent, M. McPartlon, J. Meier, V. Reis Rogozhnikov, and K. Wu, Chai-1: Decoding the molecular interactions of life (2024), bioRxiv doi:10.1101/2024.10.10.615955.

[9] J. Wohlwend, G. Corso, S. Passaro, M. Reveiz, K. Leidal, W. Swiderski, T. Portnoi, I. Chinn, J. Silterra, T. Jaakkola, and R. Barzilay, Boltz-1: Democratizing biomolecular interaction modeling (2024), bioRxiv doi:10.1101/2024.11.19.624167.

[10] J. Sohl-Dickstein, E. A. Weiss, N. Maheswaranathan, and S. Ganguli, Deep unsupervised learning using nonequilibrium thermodynamics (2015), 1503.03585.

[11] Y. Song, J. Sohl-Dickstein, D. P. Kingma, A. Kumar, S. Ermon, and B. Poole, Score-based generative modeling through stochastic differential equations (2020), 2011.13456.

[12] J. Jumper, R. Evans, A. Pritzel, T. Green, M. Figurnov, O. Ronneberger, K. Tunyasuvunakool, R. Bates, A. Žídek, A. Potapenko, et al., Highly accurate protein structure prediction with alphafold, Nature 596, 583 (2021).

[13] G. Corso, H. Stärk, B. Jing, R. Barzilay, and T. Jaakkola, DiffDock: Diffusion steps, twists, and turns for molecular docking (2022), 2210.01776.

[14] S. Passaro, G. Corso, J. Wohlwend, M. Reveiz, S. Thaler, V. R. Somnath, N. Getz, T. Portnoi, J. Roy, H. Stark, D. Kwabi-Addo, D. Beaini, T. Jaakkola, and R. Barzilay, Boltz-2: Towards accurate and efficient binding affinity prediction (2025), bioRxiv doi:10.1101/2025.06.14.659707.

[15] The OpenFold3 Team, Openfold3-preview (2025).

[16] A. Leaver-Fay, M. Tyka, S. M. Lewis, O. F. Lange, J. Thompson, R. Jacak, K. W. Kaufman, P. D. Renfrew, C. A. Smith, W. Sheffler, I. W. Davis, S. Cooper, A. Treuille, D. J. Mandell, F. Richter, Y.-E. A. Ban, S. J. Fleishman, J. E. Corn, D. E. Kim, S. Lyskov, M. Berrondo, S. Mentzer, Z. Popović, J. J. Havranek, J. Karanicolas, R. Das, J. Meiler, T. Kortemme, J. J. Gray, B. Kuhlman, D. Baker, and P. Bradley, Rosetta3, in Computer Methods, Part C (Elsevier, 2011) p. 545–574.

[17] R. F. Alford, A. Leaver-Fay, J. R. Jeliazkov, M. J. O’Meara, F. P. DiMaio, H. Park, M. V. Shapovalov, P. D. Renfrew, V. K. Mulligan, K. Kappel, J. W. Labonte, M. S. Pacella, R. Bonneau, P. Bradley, R. L. Dunbrack, R. Das, D. Baker, B. Kuhlman, T. Kortemme, and J. J. Gray, The rosetta all-atom energy function for macromolecular modeling and design, Journal of Chemical Theory and Computation 13, 3031–3048 (2017).

[18] J. P. Roney and S. Ovchinnikov, State-of-the-Art Estimation of Protein Model Accuracy Using AlphaFold, Physical Review Letters 129, 238101 (2022).

[19] C. B. Anfinsen, Principles that govern the folding of protein chains, Science 181, 223 (1973), 10.1126/science.181.4096.223.

[20] I. A. Vakser, Protein-protein docking: From interaction to interactome, Biophysical Journal 107, 1785–1793 (2014).

[21] L.-S. Chu, S. Sarma, and J. J. Gray, Unified sampling and ranking for protein docking with dfmdock (2024), bioRxiv doi:10.1101/2024.09.27.615401.

[22] D. J. Wales, Exploring energy landscapes, Annual Review of Physical Chemistry 69, 401–425 (2018).

[23] S. Sledzieski, R. Singh, L. Cowen, and B. Berger, D-script translates genome to phenome with sequence-based, structure-aware, genome-scale predictions of protein-protein interactions, Cell Systems 12, 969 (2021).

[24] S.-Y. Huang, Search strategies and evaluation in protein–protein docking: principles, advances and challenges, Drug Discovery Today 19, 1081–1096 (2014).

[25] N. A. Marze, S. S. Roy Burman, W. Sheffler, and J. J. Gray, Efficient flexible backbone protein–protein docking for challenging targets, Bioinformatics 34, 3461–3469 (2018).

[26] G. Lemmon and J. Meiler, Rosetta ligand docking with flexible xml protocols, in Computational Drug Discovery and Design (Springer New York, 2011) p. 143–155.

[27] M. A. Ketata, C. Laue, R. Mammadov, H. Stärk, M. Wu, G. Corso, C. Marquet, R. Barzilay, and T. S. Jaakkola, DiffDock-PP: Rigid protein-protein docking with diffusion models (2023), 2304.03889.

[28] F. Sverrisson, M. Akdel, D. Abramson, J. Feydy, A. Goncearenco, Y. Adeshina, D. Kovtun, C. Marquet, X. Zhang, D. Baugher, et al., Diffmasif: Surface-based protein-protein docking with diffusion models, in Machine Learning in Structural Biology workshop at NeurIPS 2023 (2023).

[29] M. McPartlon, C. Marquet, T. Geffner, D. Kovtun, A. Goncearenco, Z. Carpenter, L. Naef, M. Bronstein, and J. Xu, Latentdock: Protein-protein docking with latent diffusion, MLSB (2023).

[30] L. Ambrogioni, The statistical thermodynamics of generative diffusion models: Phase transitions, symmetry breaking, and critical instability, Entropy 27, 291 (2025).

[31] A. Sclocchi, A. Favero, and M. Wyart, A phase transition in diffusion models reveals the hierarchical nature of data, Proceedings of the National Academy of Sciences 122, 10.1073/pnas.2408799121 (2025).

[32] G. Biroli, T. Bonnaire, V. de Bortoli, and M. Mézard, Dynamical regimes of diffusion models, Nature Communications 15, 10.1038/s41467-024-54281-3 (2024).

[33] Y. Liu, Q. Yu, D. Wang, and M. Chen, Af3score: A score-only adaptation of alphafold3 for biomolecular structure evaluation, Journal of Chemical Information and Modeling 65, 8207 (2025), pMID: 40671257, 10.1021/acs.jcim.5c00653.

[34] S. Zaidi, M. Schaarschmidt, J. Martens, H. Kim, Y. W. Teh, A. Sanchez-Gonzalez, P. Battaglia, R. Pascanu, and J. Godwin, Pre-training via Denoising for Molecular Property Prediction 10.48550/arXiv.2206.00133 (2022), 2206.00133 [cs].

[35] Y. Song and D. P. Kingma, How to Train Your Energy-Based Models (2021), 2101.03288 [cs].

[36] W. Jin, X. Chen, A. Vetticaden, S. Sarzikova, R. Raychowdhury, C. Uhler, and N. Hacohen, DSMBind: SE(3) denoising score matching for unsupervised binding energy prediction and nanobody design (2023), bioRxiv doi:10.1101/2023.12.10.570461.

[37] V. R. Somnath, P. G. Sessa, M. R. Martinez, and A. Krause, DockGame: Cooperative Games for Multimeric Rigid Protein Docking (2023), 2310.06177 [cs].

[38] H. Wu, W. Liu, Y. Bian, J. Wu, N. Yang, and J. Yan, EBM-Dock: Neural probabilistic protein-protein docking via a differentiable energy model, in The Twelfth International Conference on Learning Representations (2024).

[39] K. Borisiak, G. M. Visani, and A. Nourmohammad, Loop-Diffusion: An equivariant diffusion model for designing and scoring protein loops (2024), 2409.18201 [physics].

[40] J. P. Roney, C. Ou, and S. Ovchinnikov, Protein diffusion models as statistical potentials (2026), bioRxiv doi: 10.1101/2019.12.11.123456.

[41] F. Guth, Z. Kadkhodaie, and E. P. Simoncelli, Learning normalized image densities via dual score matching (2026), 2506.05310 [cs.LG].

[42] M. Plainer, H. Wu, L. Klein, S. Günnemann, and F. Noé, Consistent sampling and simulation: Molecular dynamics with energy-based diffusion models (2025), 2506.17139 [cs.LG].

[43] M. Arts, V. G. Satorras, C.-W. Huang, D. Zuegner, M. Federici, C. Clementi, F. Noé, R. Pinsler, and R. van den Berg, Two for One: Diffusion Models and Force Fields for Coarse-Grained Molecular Dynamics 10.48550/arXiv.2302.00600 (2023), 2302.00600 [cs].

[44] S. Raja, M. Šípka, M. Psenka, T. Kreiman, M. Pavelka, and A. S. Krishnapriyan, Action-Minimization Meets Generative Modeling: Efficient Transition Path Sampling with the Onsager-Machlup Functional (2025), 2504.18506 [cs].

[45] J. He, Y. Du, F. Vargas, Y. Wang, C. P. Gomes, J. M. Hernández-Lobato, and E. Vanden-Eijnden, FEAT: Free energy Estimators with Adaptive Transport (2026), 2504.11516 [stat].

[46] M. S. Albergo, N. M. Boffi, and E. Vanden-Eijnden, Stochastic Interpolants: A Unifying Framework for Flows and Diffusions (2025), 2303.08797 [cs].

[47] R. T. Q. Chen, Y. Rubanova, J. Bettencourt, and D. Duvenaud, Neural ordinary differential equations 2 (2019), 1806.07366 [cs.LG].

[48] D. L. Beveridge and F. M. DiCapua, Free energy via molecular simulation: Applications to chemical and biomolecular systems, Annual Review of Biophysics and Biophysical Chemistry 18, 431–492 (1989).

[49] R. Abel, L. Wang, E. D. Harder, B. J. Berne, and R. A. Friesner, Advancing drug discovery through enhanced free energy calculations, Accounts of Chemical Research 50, 1625–1632 (2017).

[50] B. D. Anderson, Reverse-time diffusion equation models, Stochastic Processes and their Applications 12, 313 (1982).

[51] P. Vincent, A connection between score matching and denoising autoencoders, Neural Computation 23, 1661 (2011).

[52] B. Øksendal, Stochastic differential equations, in Stochastic Differential Equations: An Introduction with Applications (Springer Berlin Heidelberg, Berlin, Heidelberg, 2003) pp. 65–84.

[53] B. Øksendal, Stochastic differential equations, in Stochastic Differential Equations: An Introduction with Applications (Springer Berlin Heidelberg, Berlin, Heidelberg, 2003) pp. 35–37.

[54] T. Vreven, I. H. Moal, A. Vangone, B. G. Pierce, P. L. Kastritis, M. Torchala, R. Chaleil, B. Jiménez-García, P. A. Bates, J. Fernandez-Recio, A. M. Bonvin, and Z. Weng, Updates to the integrated protein–protein interaction benchmarks: Docking benchmark version 5 and affinity benchmark version 2, Journal of Molecular Biology 427, 3031 (2015).

[55] J. D. Guest, T. Vreven, J. Zhou, I. Moal, J. R. Jeliazkov, J. J. Gray, Z. Weng, and B. G. Pierce, An expanded benchmark for antibody-antigen docking and affinity prediction reveals insights into antibody recognition determinants, Structure 29, 606 (2021).

[56] S. Basu and B. Wallner, DockQ: A Quality Measure for Protein-Protein Docking Models, 11, e0161879.

[57] J. Janin, K. Henrick, J. Moult, L. T. Eyck, M. J. E. Sternberg, S. Vajda, I. Vakser, and S. J. Wodak, CAPRI: A Critical Assessment of PRedicted Interactions, Proteins: Structure, Function, and Bioinformatics 52, 2 (2003).

[58] S. Lin, B. Liu, J. Li, and X. Yang, Common Diffusion Noise Schedules and Sample Steps are Flawed, 2024 IEEE/CVF Winter Conference on Applications of Computer Vision (WACV), 5392 (2024).

[59] G. Franzese, S. Rossi, L. Yang, A. Finamore, D. Rossi, M. Filippone, and P. Michiardi, How Much Is Enough? A Study on Diffusion Times in Score-Based Generative Models, Entropy 25, 633 (2023).

[60] T. Salimans and J. Ho, Should EBMs model the energy or the score?, in Energy Based Models Workshop-ICLR 2021 (2021).

[61] C.-H. Lai, Y. Takida, N. Murata, T. Uesaka, Y. Mitsufuji, and S. Ermon, Fp-diffusion: Improving score-based diffusion models by enforcing the underlying score fokker-planck equation (2023), 2210.04296 [cs.LG].

[62] G. Ahdritz, N. Bouatta, C. Floristean, S. Kadyan, Q. Xia, W. Gerecke, T. J. O’Donnell, D. Berenberg, I. Fisk,N. Zanichelli, B. Zhang, A. Nowaczynski, B. Wang, M. M. Stepniewska-Dziubinska, S. Zhang, A. Ojewole, M. E. Guney, S. Biderman, A. M. Watkins, S. Ra, P. R. Lorenzo, L. Nivon, B. Weitzner, Y.-E. A. Ban, S. Chen, M. Zhang, C. Li, S. L. Song, Y. He, P. K. Sorger, E. Mostaque, Z. Zhang, R. Bonneau, and M. AlQuraishi, OpenFold: Retraining AlphaFold2 yields new insights into its learning mechanisms and capacity for generalization, Nature Methods 21, 1514 (2024).

[63] J. P. Roney and S. Ovchinnikov, State-of-the-art estimation of protein model accuracy using alphafold, Physical Review Letters 129, 10.1103/physrevlett.129.238101 (2022).

[64] P. Kulytė, F. Vargas, S. V. Mathis, Y. G. Wang, J. M. Hernández-Lobato, and P. Liò, Improving antibody design with force-guided sampling in diffusion models (2024), 2406.05832 [q-bio.QM].

[65] Y. Wang, L. Wang, Y. Shen, Y. Wang, H. Yuan, Y. Wu, and Q. Gu, Protein conformation generation via force-guided SE(3) diffusion models, in Proceedings of the 41st International Conference on Machine Learning, Proceedings of Machine Learning Research, Vol. 235, edited by R. Salakhutdinov, Z. Kolter, K. Heller, A. Weller, N. Oliver, J. Scarlett, and F. Berkenkamp (PMLR, 2024) pp. 56835–56859.

[66] S. Lewis, T. Hempel, J. Jiménez-Luna, M. Gastegger, Y. Xie, A. Y. Foong, V. G. Satorras, O. Abdin, B. S. Veeling, I. Zaporozhets, Y. Chen, S. Yang, A. E. Foster, A. Schneuing, J. Nigam, F. Barbero, S. Vincent, A. Campbell, J. Yim, M. Lienen, Y. Shi, S. Zheng, H. Schulz, U. Munir, R. Sordillo, R. Tomioka, C. Clementi, and F. Noé, Scalable emulation of protein equilibrium ensembles with generative deep learning, Science, eadv9817 (2025).

[67] A. Morehead, C. Chen, A. Sedova, and J. Cheng, DIPS-Plus: The enhanced database of interacting protein structures for interface prediction, 10, 509.

[68] R. T. Q. Chen, torchdiffeq (2018).

