## Supplementary Material for "Can We Extract Physics-like Energies from Generative Protein Diffusion Models?"

#### A. Dimensional Analysis of Key Equations

We perform dimensional analysis on the fundamental equations underlying diffusion (and their physics counterparts) to derive the units for **Table I**; $[M]$ represents units of mass, $[L]$ units of length, $[T]$ units of *physical* time, and $[\tau]$ units of *diffusion/alchemical* time. We begin with the forward diffusion equation, eq. (3):

$$dx_t = f(x_t,t)dt + g(t)dw_t.$$

where $x_t$ represents a point in Euclidean space $\mathbb{R}^n$ and thus has units of length $[L]$. More generally, $x_t$ can be a point on an arbitrary Riemannian manifold; every such manifold has a Riemannian metric, a measure of distance between two points, so units of length still broadly apply. The Wiener term $dw_t$ is sampled from a Gaussian with variance $dt$, defined in terms of alchemical/diffusion time, meaning its units must be $[\tau]^{1/2}$, the square root of the unit of the variance. The units of the functions $f$ and $g$ can thus be inferred:

$$dx_t \sim [L], \qquad dt \sim [\tau], \qquad dw_t \sim [\tau]^{1/2},$$

$$f(x_t,t) \sim \frac{[L]}{[\tau]}, \qquad \text{“Drift”}$$

$$g(t) \sim \frac{[L]}{[\tau]^{1/2}}. \qquad \text{“Diffusion Coefficient (AI)”}$$

The reverse diffusion equation, eq. (5),

$$dx_t = \left[f(x_t,t) - g(t)^2\nabla_x \log p(x_t,t)\right]dt + g(t)dw_t$$

introduces the marginal probability $p(x_t,t)$, which is unitless, and reveals the units of $g(t)^2$ and $\nabla_x \log p(x_t,t)$:

$$p(x_t,t) \sim [1],$$

$$g(t)^2 \sim \frac{[L]^2}{[\tau]}, \qquad \text{“Diffusion Coefficient (Physics)”}$$

$$\nabla_x \log p(x_t,t) \sim \frac{1}{[L]}. \qquad \text{“Score”}$$

Equations (11) and (12) connect the marginal probability to energy:

$$p(x_t,t) = \frac{e^{-\beta E_t(x_t)}}{Z};$$
$$\log p(x_t,t) = -\beta E_t(x_t) - \log Z.$$

These equations tell us $\beta E_t(x_t)$ and $Z$ must be unitless, and the dimensions of energy are defined by physics:

$$Z \sim [1], \quad E_t(x_t) \sim \frac{[M][L]^2}{[T]^2}, \quad \beta \sim \frac{[T]^2}{[M][L]^2},$$

$$-\beta^{-1}\log p(x_t,t) \sim \frac{[M][L]^2}{[T]^2}. \qquad \text{“Free Energy”}$$

Finally, equation (14) connects diffusion score and physics:

$$s_\theta(x_t,t) \approx \nabla_x \log p(x_t,t) = -\beta\nabla_x E_t(x_t),$$

introducing learned score $s_\theta(x_t,t)$ and force $-\nabla_x E_t(x_t)$:

$$s_\theta(x_t,t) \sim \frac{1}{[L]}, \qquad \text{“Learned Score”}$$

$$\nabla_x E_t(x_t) = -\beta^{-1}\nabla_x \log p(x_t,t) \sim \frac{[M][L]}{[T]^2}. \qquad \text{“Force”}$$

For completeness, we also include dimensional analysis of our integral formulations. For eq. (20):

$$\Delta \log p = \int_0^1 \left(\nabla_x \log p(x_t,t)\cdot\frac{\partial x_t}{\partial t} + \frac{\partial \log p(x_t,t)}{\partial t}\right)dt,$$

$$\frac{\partial x_t}{\partial t} \sim \frac{[L]}{[\tau]}, \qquad \frac{\partial \log p(x_t,t)}{\partial t} \sim \frac{[1]}{[\tau]}.$$

And eq. (21):

$$\frac{\partial \log p(x_t,t)}{\partial t} = \frac{1}{2}g(t)^2\Big(\nabla_x\cdot\nabla_x \log p(x_t,t)\Big) + \frac{1}{2}g(t)^2\|\nabla_x \log p(x_t,t)\|^2,$$

$$\frac{1}{2}g(t)^2\Big(\nabla_x\cdot\nabla_x \log p(x_t,t)\Big) \sim \frac{[L]^2}{[\tau]}\cdot\frac{[1]}{[L]^2} = \frac{[1]}{[\tau]},$$

$$\frac{1}{2}g(t)^2\|\nabla_x \log p(x_t,t)\|^2 \sim \frac{[L]^2}{[\tau]}\cdot\frac{[1]}{[L]^2} = \frac{[1]}{[\tau]}.$$

### B. Analysis of Prior Accuracy and Noise

A major assumption in diffusion modeling is that by the time horizon $t = 1$, sufficient noise has been added to the data that the distribution of the marginal $p(x_1, 1)$ is determined solely by the noise schedule. Specifically, we assume that $x_1 \sim p(x_1, 1) = p_1(x_1) = \mathcal{N}(x_1; \mu = 0, \sigma = \sigma_1)$. However, the accuracy of this assumption is highly dependent on both the magnitude of the added noise and the scale and variance of the true data distribution. As we rely on the prior as the single source of absolute probability along our integrated trajectories, any discrepancy between the assumed prior and the true marginal at $t = 1$ could have significant effects on both sampling and likelihood recovery.

For our trimodal Gaussian toy model, we initially chose the noise schedule hyperparameters ($\sigma_0 = 0.1$ and $\sigma_1 = 30$) to match those of DFMDock. This choice, however, turned out to yield a prior with significant inaccuracies, as will be demonstrated below. Although the inaccuracy of the prior is difficult to measure on real systems, we can easily compute the true marginal at time $t = 1$ for our toy Gaussian model. Recall that applying the variance-exploding forward diffusion process from time $t = 0$ to $t = 1$ to the data distribution has the effect of adding white noise with mean 0 and variance $\sigma^2_{\text{noise}} = \sigma_1^2 - \sigma_0^2$ to the data distribution (eq. (4)). Because the sum of two Gaussian random variables is itself Gaussian, adding this white noise to our mixture-of-Gaussians toy model produces a mixture-of-Gaussians $t = 1$ marginal distribution with variances $b^2_{i_{\text{noised}}} = b_i^2 + \sigma_1^2 - \sigma_0^2$, means $\mu_i$, and weights $w_i$ as defined in **Section VI.A**.

Plotting this ground-truth $t = 1$ marginal and assumed $t = 1$ prior distributions with $\sigma_0 = 0.1$ and $\sigma_1 = 30$ reveals that the true marginal is far from Gaussian in shape and is highly divergent from the assumed prior, with a KL divergence of 0.11 (**Supp. Fig. S3a**, left panel). As expected, this also leads to inaccurately recovered likelihoods (**Supp. Fig. S3a**, right two panels, red). To illustrate the error caused by an inaccurate prior, we computed the learned likelihoods using the true prior instead of the assumed Gaussian prior (**Supp. Fig. S3a**, right two panels, light blue). That is, we used the same $\Delta \log p$, but instead of computing $\log p_1(x_1)$ from the assumed Gaussian, we used the true value of the marginal distribution at time $t = 1$. As expected, likelihoods computed from an accurate marginal distribution are significantly more accurate to the ground truth distribution (dashed black line) and the sample KDE (blue line).

A surprising result of this comparison is that diffusion trajectories improve differently than flow trajectories when computed with a more accurate prior. Assumed-prior flow likelihoods (**Supp. Fig. S3a**, right panel) produce a significantly narrower and taller distribution than true-marginal flow trajectories, while retaining continuity and a similar overall shape. This neatly mirrors the way the assumed prior is narrower and taller than the true $t = 1$ marginal. By contrast, assumed-prior diffusion likelihoods (**Supp. Fig. S3a**, center panel) do not exhibit this narrowing and are instead excessively noisy. To estimate the bias of assumed-prior diffusion probabilities, we plot a binned average (red line), which overestimates the true probability and sample KDE on the left and right modes but is largely accurate on the central mode.

The key reason behind the different effects of prior inaccuracy on flow and diffusion trajectories is the stochasticity - and thus non-continuity - of the relationship between the endpoints of diffusion trajectory paths in the data distribution and the prior distribution. In other words, since diffusion paths are stochastic, very nearby or even identical $t = 0$ samples can be the result of diffusion trajectories starting from wildly different points in the prior, as illustrated in **Fig. 1a**. As the difference between the true and assumed prior distribution depends on $x_1$, this causes adjacent samples to carry very different prior error, producing noise. On the other hand, since flow trajectories are non-crossing ODEs, prior error continuously *biases* the result without adding noise as the error is continuously sampled.

In practical situations, we cannot compute the true data marginal, as we typically only have access to a biased set of discrete observations from the full data space. Thus, accurate likelihood estimation requires decreasing the discrepancy between the assumed prior and the true marginal. Recall that the assumed prior, a Gaussian with standard deviation $\sigma_{\text{prior}} = \sigma_1$, is derived by assuming that sufficient Gaussian noise is added to the data that the result is also Gaussian. As the true $t = 1$ marginal with $\sigma_1 = 30$ is clearly non-Gaussian, a simple solution is to increase the value of $\sigma_1$ and retrain the network. Increasing $\sigma_1$ from 30 to 70 produced a much more Gaussian-like prior (**Supp. Fig. S3b**, left panel), with less noisy diffusion likelihoods (**Supp. Fig. S3b**, center panel) and less biased flow trajectories (**Supp. Fig. S3b**, right panel). However, there still remains some discrepancy between the assumed and true prior (KL = 0.031), which makes assumed-prior diffusion and flow results respectively noisier and more biased than their true-prior counterparts.

To further improve the accuracy of the prior, recall that the assumed prior has mean 0 and standard deviation $\sigma_{\text{prior}} = \sigma_1$. These values are derived from assumptions about our data distribution: $\mu_{\text{data}} = 0$, and $\sigma_{\text{data}} = \sigma_0$. While our toy dataset is indeed approximately centered at 0, the standard deviation of our training samples—approximately 29.975—is significantly greater than our original $\sigma_0$ of 0.1. Increasing $\sigma_0$ to 30 significantly increased the accuracy of approximating the true marginal with the assumed prior, with a KL-divergence of $6.0 \cdot 10^{-5}$ (**Supp. Fig. S3c**, left panel). Retraining the model with the new noise schedule and integrating from the assumed prior yielded significantly improved likelihoods, and recomputing using the true $t = 1$ marginal now barely affects the result (**Supp. Fig. S3b**, right panels, light blue covering red). We therefore used the noise schedule $\sigma_0 = 30$ and $\sigma_1 = 70$ for our toy model throughout the main text.

Clearly, determination of both $\sigma_0$ and the prior mean rely heavily on accurate estimates of $\mu_{\text{data}}$ and $\sigma_{\text{data}}$, and the error caused by incorrect estimates can be significant. However, there can be cases where these estimates are infeasible in practice, such as in high-dimensional, sparse data regimes where the model is expected to generalize. Moreover, for conditional diffusion, it is unclear exactly what mean or variance these estimates should reflect; after all, the desired data distribution is by definition going to vary based on condition, which seemingly implies the need for a different $\sigma_0$ for each condition. Conveniently, one benefit of inference-time likelihood computation is that one can explore multiple choices of prior without changing the noise schedule or retraining the model, as we have done by computing likelihoods with the true prior for our toy model. While the true $t = 1$ marginal is not generally feasible to compute, perhaps other estimated priors could be?

Accordingly, we present a proof-of-concept for estimating the $t = 1$ marginal using a different prior than the standard choice. We use the original model trained with a noise schedule of $\sigma_0 = 0.1$ and $\sigma_1 = 30$, for which we know the $\sigma_0$ value is an improper estimate of the true data standard deviation. Recall the formula for $\sigma^2_{\text{prior}} = \sigma_1^2 - \sigma_0^2 + \sigma^2_{\text{data}}$; if we do not make the assumption that $\sigma_{\text{data}} = \sigma_0$, and instead use the knowledge that $\sigma_{\text{data}} \approx 29.975$, we can compute a more accurate estimate for $\sigma_{\text{prior}}$: $\sqrt{30^2 - 0.1^2 + 29.975^2} \approx 42.4$. **Supp. Fig. S3d**, left panel plots this estimated prior distribution with $\mu = 0$ and $\sigma = 42.4$ against the true $t = 1$ marginal for the $\sigma_0 = 0.1$, $\sigma_1 = 30$ diffusion process, which much more closely matches the spread of the marginal (though the marginal is still non-gaussian) with a KL divergence of 0.0016. The recovered likelihoods for both diffusion and flow are much more accurate than either of the assumed prior likelihoods with $\sigma_0 = 0.1$ (**Supp. Fig. S3a,b**), though not quite as accurate as the $\sigma_0 = 30$, $\sigma_1 = 70$ model due to the prior inaccuracy caused by the non-gaussian nature of the $t = 1$ marginal.

These results demonstrate the feasibility and potential of estimating the $t = 1$ marginal for models which are either costly to retrain (such as large models with proprietary datasets and closed-source training code) or are limited in their ability to estimate the true data mean and variance (such as low-data conditional diffusion models). Protein structure diffusion models commonly fall into both of these categories, so we believe there is significant potential in accurate estimation of the potentially complex marginal distributions for these models, both for improved sampling and for accurate likelihood computation. Finally, we note that even with perfect priors, diffusion likelihoods still have noise and flow likelihoods (can) still have bias. We believe this phenomenon is due to score model inaccuracy/non-conservativity, which we discuss at the end of **Section V**.

#### C. Derivation of Logarithmic Fokker Planck Equation

The Fokker-Planck equation is a partial differential equation (PDE) that describes the time-evolution of the marginal probability distribution $p(x_t, t)$ associated with a given stochastic differential equation (SDE). It has many variants depending on the form of the SDE; for $dx_t = f(x_t, t)dt + g(t)dw_t$ the Fokker-Planck equation is [52]

$$\begin{aligned}\frac{\partial p(x_t,t)}{\partial t} &= -\sum_{i=0}^{n} \frac{\partial}{\partial [x_t]_i}\Big([f(x_t,t)]_i \, p(x_t,t)\Big) \\ &\quad + \frac{1}{2} g(t)^2 \sum_{i=0}^{n} \frac{\partial^2}{\partial [x_t]_i^2} p(x_t,t), \end{aligned} \tag{S-1}$$

where $n$ is the dimensionality of $x_t$, $[x_t]_i$ is the $i$-th component of $x_t$, and $[f(x_t, t)]_i$ is the $i$-th component of the drift term $f(x_t, t)$.

The Fokker-Planck equation is useful, but it is dependent on the value and spatial gradients of probability $p(x_t, t)$, neither of which we have; rather, our score model is trained to predict the spatial gradients of the log probability. We can convert between the two using the chain rule:

$$\begin{aligned}\frac{\partial \log f(x)}{\partial x_i} &= \frac{d \log f(x)}{df(x)} \frac{\partial f(x)}{\partial x_i} \\ &= \frac{1}{|f(x)|} \frac{\partial f(x)}{\partial x_i}.\end{aligned}$$

Since $p(x_t, t) > 0$,

$$\frac{\partial \log p(x_t,t)}{\partial t} = \frac{1}{p(x_t,t)} \frac{\partial p(x_t,t)}{\partial t}. \tag{S-2}$$

Thus, we can rewrite the Fokker-Planck equation (S-1) in a logarithmic form like so:

$$\begin{aligned}\frac{\partial \log p(x_t,t)}{\partial t} &= \frac{1}{p(x_t,t)}\frac{\partial p(x_t,t)}{\partial t} \\ &= -\frac{1}{p(x_t,t)}\sum_{i=1}^{n}\frac{\partial}{\partial [x_t]_i}\Big([f(x_t,t)]_i\, p(x_t,t)\Big) \\ &+ \frac{1}{p(x_t,t)}\frac{1}{2}g(t)^2\sum_{i=1}^{n}\frac{\partial^2}{\partial [x_t]_i^2}p(x_t,t). \end{aligned} \tag{S-3}$$

This equation is still dependent on derivatives of $p(x_t,t)$, but we can convert them into logarithmic derivatives as well by multiplying by $\frac{1}{p(x_t,t)}$. The first-order term is relatively straightforward; first, we expand the derivative using the product rule:

$$\begin{aligned}\frac{\partial \log p(x_t,t)}{\partial t} &= -\frac{1}{p(x_t,t)}\sum_{i=1}^{n}p(x_t,t)\left(\frac{\partial}{\partial [x_t]_i}[f(x_t,t)]_i\right) \\ &- \frac{1}{p(x_t,t)}\sum_{i=1}^{n}[f(x_t,t)]_i\left(\frac{\partial}{\partial [x_t]_i}p(x_t,t)\right) \\ &+ \frac{1}{p(x_t,t)}\frac{1}{2}g(t)^2\sum_{i=1}^{n}\frac{\partial^2}{\partial [x_t]_i^2}\left(p(x_t,t)\right). \end{aligned} \tag{S-4}$$

In the first term, $p(x_t,t)$ cancels, and in the second term, the $p(x_t,t)$ derivative becomes logarithmic by eq. (S-2):

$$\begin{aligned}\frac{\partial \log p(x_t,t)}{\partial t} &= -\sum_{i=1}^{n}\frac{\partial}{\partial [x_t]_i}[f(x_t,t)]_i \\ &- \sum_{i=1}^{n}[f(x_t,t)]_i\left(\frac{\partial}{\partial [x_t]_i}\log p(x_t,t)\right) \\ &+ \frac{1}{p(x_t,t)}\frac{1}{2}g(t)^2\sum_{i=1}^{n}\frac{\partial^2}{\partial [x_t]_i^2}p(x_t,t). \end{aligned} \tag{S-5}$$

The second-order derivative in the third term requires more work. We can again use eq. (S-2) to derive an identity for logarithmic second derivatives:

$$\begin{aligned}\frac{\partial^2}{\partial [x_t]_i^2}p(x_t,t) &= \frac{\partial}{\partial [x_t]_i}\left(\frac{\partial}{\partial [x_t]_i}p(x_t,t)\right) \\ &= \frac{\partial}{\partial [x_t]_i}\left(\left(\frac{\partial}{\partial [x_t]_i}\log p(x_t,t)\right)p(x_t,t)\right) \\ &= \left(\frac{\partial^2}{\partial [x_t]_i^2}\log p(x_t,t)\right)p(x_t,t) + \left(\frac{\partial}{\partial [x_t]_i}\log p(x_t,t)\right)\left(\frac{\partial}{\partial [x_t]_i}p(x_t,t)\right) \\ &= \left(\frac{\partial^2}{\partial [x_t]_i^2}\log p(x_t,t)\right)p(x_t,t) + \left(\frac{\partial}{\partial [x_t]_i}\log p(x_t,t)\right)\left(\frac{\partial}{\partial [x_t]_i}\log p(x_t,t)\right)p(x_t,t) \\ &= \left(\frac{\partial^2}{\partial [x_t]_i^2}\log p(x_t,t)\right)p(x_t,t) + \left(\frac{\partial}{\partial [x_t]_i}\log p(x_t,t)\right)^2 p(x_t,t). \end{aligned} \tag{S-6}$$

Substituting this back into equation (S-5), the problematic term splits into a second-order logarithmic term and a squared first-order term, and the final remaining $p(x_t, t)$ coefficients cancel:

$$\begin{aligned}\frac{\partial \log p(x_t,t)}{\partial t} &= -\sum_{i=1}^{n} \frac{\partial}{\partial [x_t]_i}[f(x_t,t)]_i \\ &- \sum_{i=1}^{n} [f(x_t,t)]_i \left(\frac{\partial}{\partial [x_t]_i} \log p(x_t,t)\right) \\ &+ \frac{1}{2} g(t)^2 \sum_{i=1}^{n} \frac{\partial^2}{\partial [x_t]_i^2} \log p(x_t,t) \\ &+ \frac{1}{2} g(t)^2 \sum_{i=1}^{n} \left(\frac{\partial}{\partial [x_t]_i} \log p(x_t,t)\right)^2, \end{aligned} \tag{S-7}$$

giving us a form of the logarithmic Fokker-Planck equation that we can evaluate using only a trained score model. These sums can also be written using vector notation:

$$\begin{aligned}\frac{\partial \log p(x_t,t)}{\partial t} &= -\nabla_x \cdot f(x_t,t) \\ &- f(x_t,t) \cdot \nabla_x \log p(x_t,t) \\ &+ \frac{1}{2} g(t)^2 \nabla_x \cdot \nabla_x \log p(x_t,t) \\ &+ \frac{1}{2} g(t)^2 \|\nabla_x \log p(x_t,t)\|^2. \end{aligned} \tag{S-8}$$

Finally, since our models are variance-exploding, their drift term $f(x_t, t) = 0$, simplifying to eq. (21) in the main text:

$$\begin{aligned}\frac{\partial \log p(x_t,t)}{\partial t} &= \frac{1}{2} g(t)^2 \Big(\nabla_x \cdot \nabla_x \log p(x_t,t)\Big) \\ &+ \frac{1}{2} g(t)^2 \|\nabla_x \log p(x_t,t)\|^2. \end{aligned} \tag{21}$$

#### C.1. Reverse-Time Fokker-Planck

Technically, the process whose marginal distribution we seek is the *reverse* diffusion process, eq. (5):

$$dx_t = [f(x_t,t) - g(t)^2 \nabla_x \log p(x_t,t)]dt + g(t) d\overline{w}_t. \tag{5}$$

As mentioned earlier, the reverse diffusion process is carefully chosen such that its marginal distribution matches that of the forward process, and therefore the Fokker-Planck equations should be identical for both (in fact, Anderson (1982) [50] derived the reverse SDE precisely by matching the Fokker-Planck for the forward and backward equations). For thoroughness, we will also derive the marginal distribution for the reverse process.

The most important thing to keep track of is the direction of time and the sign of its change. The Fokker-Planck equation assumes a positive $dt$ and a forward-time Wiener process $dw_t$, but the reverse SDE involves a negative $dt$ and a reverse-time Wiener process $d\overline{w}_t$. To adjust, we perform change-of-variables on $t$ to produce an equivalent forward-time process in the variable $s$; by replacing $t$ in eq. (5) with $1 - s$, changing the sign of the first term to account for replacing $dt < 0$ with $ds > 0$, and using the forward-time Wiener term:

$$dx_{1-s} = [-f(x_{1-s}, 1-s) + g(1-s)^2 \nabla_x \log p(x_{1-s}, 1-s)]ds + g(1-s) dw_s. \tag{S-9}$$

The logarithmic Fokker-Planck equation for this SDE then describes the change in the reverse marginal probability, $\overline{p}$, with respect to $s$:

$$
\begin{aligned}
\frac{\partial \log \overline{p}(x_{1-s}, 1-s)}{\partial s} &= -\nabla_x \cdot [-f(x_{1-s}, 1-s) + g(1-s)^2 \nabla_x \log p(x_{1-s}, 1-s)] \\
&- [-f(x_{1-s}, 1-s) + g(1-s)^2 \nabla_x \log p(x_{1-s}, 1-s)] \cdot \nabla_x \log \overline{p}(x_{1-s}, 1-s) \\
&+ \frac{1}{2} g(1-s)^2 \nabla_x \cdot \nabla_x \log \overline{p}(x_{1-s}, 1-s) \\
&+ \frac{1}{2} g(1-s)^2 \|\nabla_x \log \overline{p}(x_{1-s}, 1-s)\|^2 \\
\\
&= \nabla_x \cdot f(x_{1-s}, 1-s) - g(1-s)^2 \nabla_x \cdot \nabla_x \log p(x_{1-s}, 1-s) \\
&+ f(x_{1-s}, 1-s) \cdot \nabla_x \log \overline{p}(x_{1-s}, 1-s) - g(1-s)^2 \nabla_x \log p(x_{1-s}, 1-s) \cdot \nabla_x \log \overline{p}(x_{1-s}, 1-s) \\
&+ \frac{1}{2} g(1-s)^2 \nabla_x \cdot \nabla_x \log \overline{p}(x_{1-s}, 1-s) \\
&+ \frac{1}{2} g(1-s)^2 \|\nabla_x \log \overline{p}(x_{1-s}, 1-s)\|^2 \\
\\
&= \nabla_x \cdot f(x_{1-s}, 1-s) \\
&+ f(x_{1-s}, 1-s) \cdot \nabla_x \log \overline{p}(x_{1-s}, 1-s) \\
&+ g(1-s)^2 \nabla_x \cdot \nabla_x \left( \frac{1}{2} \log p(x_{1-s}, 1-s) - \log \overline{p}(x_{1-s}, 1-s) \right) \\
&+ g(1-s)^2 \nabla_x \log \overline{p}(x_{1-s}, 1-s) \cdot \nabla_x \left( \frac{1}{2} \log p(x_{1-s}, 1-s) - \log \overline{p}(x_{1-s}, 1-s) \right). \qquad \text{(S-10)}
\end{aligned}
$$

To compare with the Fokker-Planck for the forward process, eq. (S-8), we'd like the derivative with respect to $t$, not $s$. Re-substituting $1-s$ with $t$ and using the chain rule:

$$
\begin{aligned}
\frac{\partial \log \overline{p}(x_{1-s}, 1-s)}{\partial s} &= \frac{\partial \log \overline{p}(x_t, t)}{\partial (1-t)} \\
&= -\frac{\partial \log \overline{p}(x_t, t)}{\partial t}, \qquad \text{(S-11)}
\end{aligned}
$$

which means the temporal gradient of the reverse process in terms of $t$ is:

$$
\begin{aligned}
\frac{\partial \log \overline{p}(x_t, t)}{\partial t} &= -\frac{\partial \log \overline{p}(x_{1-s}, 1-s)}{\partial s} \\
&= -\nabla_x \cdot f(x_t, t) \\
&- f(x_t, t) \cdot \nabla_x \log \overline{p}(x_t, t) \\
&+ g(t)^2 \nabla_x \cdot \nabla_x \left( \log \overline{p}(x_t, t) - \frac{1}{2} \log p(x_t, t) \right) \\
&+ g(t)^2 \nabla_x \log \overline{p}(x_t, t) \cdot \nabla_x \left( \log \overline{p}(x_t, t) - \frac{1}{2} \log p(x_t, t) \right). \qquad \text{(S-12)}
\end{aligned}
$$

To show that $p(x_t,t)=\overline{p}(x_t,t)$, then, it is sufficient to show that $p(x_t,t)$ is a solution to eq. (S-12) and that both functions satisfy the same initial value problems. That $p(x_t,t)$ satisfies eq. (S-12) is easy to show by substituting $p(x_t,t)$ for $\overline{p}(x_t,t)$ and observing that it simplifies to the forward logarithmic Fokker-Planck equation, eq. (S-8):

$$\begin{aligned}\frac{\partial \log \overline{p}(x_t,t)}{\partial t} &= -\nabla_x \cdot f(x_t,t)\\ &- f(x_t,t)\cdot \nabla_x \log p(x_t,t)\\ &+ g(t)^2 \nabla_x \cdot \nabla_x \left(\log p(x_t,t) - \frac{1}{2}\log p(x_t,t)\right)\\ &+ g(t)^2 \nabla_x \log p(x_t,t)\cdot \nabla_x \left(\log p(x_t,t) - \frac{1}{2}\log p(x_t,t)\right) \end{aligned} \tag{S-13}$$

$$\begin{aligned} &= -\nabla_x \cdot f(x_t,t)\\ &- f(x_t,t)\cdot \nabla_x \log p(x_t,t)\\ &+ \frac{1}{2}g(t)^2 \nabla_x \cdot \nabla_x \log p(x_t,t)\\ &+ \frac{1}{2}g(t)^2 \|\nabla_x \log p(x_t,t)\|^2, \end{aligned} \tag{S-8}$$

and the initial condition, $\overline{p}(x_1,1)=p_1(x_1)$, is satisfied by our assumption that we add sufficient noise for $p(x_1,1)=p_1(x_1)$. Since $p(x_t,t)$ satisfies the differential equation for $\overline{p}(x_t,t)$ with the same initial conditions, we conclude that $p(x_t,t)=\overline{p}(x_t,t)$ as expected. To save on this algebraic manipulation in the main body of the paper, we always use the Fokker-Planck of the forward equation and assume $dt>0$.

### D. Derivation of Flow Trajectory Integral

Applying eq. (26) to the flow ODE path, we can simplify the equation analytically by subbing in the path's $\frac{dx_t}{dt}$ as defined by the ODE, eq. (29):

$$\begin{aligned}\frac{dx_t}{dt} &= -\frac{1}{2}g(t)^2 \nabla_x \log p(x_t,t)\\ &\approx -\frac{1}{2}g(t)^2 s_\theta(x_t,t). \end{aligned} \tag{29}$$

Substituting this into eq. (26):

$$\log p_0(x_0) = \log p_1(x_1) - \int_0^1 \left(s_\theta(x_t,t)\cdot \frac{dx_t}{dt} + \frac{1}{2}g(t)^2 \mathrm{Tr}\Big(\nabla_x s_\theta(x_t,t)\Big) + \frac{1}{2}g(t)^2\|s_\theta(x_t,t)\|^2\right)dt \tag{26}$$

$$= \log p_1(x_1) - \int_0^1 \left(s_\theta(x_t,t)\cdot \left(-\frac{1}{2}g(t)^2 s_\theta(x_t,t)\right) + \frac{1}{2}g(t)^2 \mathrm{Tr}\Big(\nabla_x s_\theta(x_t,t)\Big) + \frac{1}{2}g(t)^2\|s_\theta(x_t,t)\|^2\right)dt$$

$$= \log p_1(x_1) - \int_0^1 \left(-\cancel{\frac{1}{2}g(t)^2\|s_\theta(x_t,t)\|^2} + \frac{1}{2}g(t)^2 \mathrm{Tr}\Big(\nabla_x s_\theta(x_t,t)\Big) + \cancel{\frac{1}{2}g(t)^2\|s_\theta(x_t,t)\|^2}\right)dt$$

$$= \log p_1(x_1) - \int_0^1 \frac{1}{2}g(t)^2 \mathrm{Tr}\Big(\nabla_x s_\theta(x_t,t)\Big)dt. \tag{30}$$

### E. Pseudocodes for Log Likelihood Integrals

These are pseudocode algorithms for integrating over discrete diffusion trajectories by forwards Euler integration (**Algorithm 1**), backward Euler integration (**Algorithm 2**), the trapezoid rule (**Algorithm 3**), interpolated discrete integration (**Algorithm 4**; compatible with Euler or trapezoidal methods), and piecewise ODE integration (**Algorithm 5**), and over flow trajectories using black-box ODE solvers (**Algorithm 6**).

**Algorithm 1** Forward Euler Integration over Diffusion Trajectories

**Input:** Diffusion trajectory samples $\{(x_{t_0}, t_0), (x_{t_1}, t_1), \ldots, (x_{t_n}, t_n)\}$ where $t_i \in [0, 1], t_0 = 0, t_n = 1$; score model $s_\theta(x_t, t)$; noise schedule $g(t)$
**Output:** Log-likelihood estimate $\log p_0(x_0) = \log p_1(x_1) - \Delta \log p$
Initialize $\Delta \log p \leftarrow 0$
**for** $i = 0$ to $n - 1$ **do**
  $\Delta x_i \leftarrow x_{t_{i+1}} - x_{t_i}$
  $\Delta t_i \leftarrow t_{i+1} - t_i$
  $\Delta \log p \leftarrow \Delta \log p + s_\theta(x_{t_i}, t_i) \cdot \Delta x + \frac{1}{2} g(t_i)^2 \mathrm{Tr}\Big(\nabla_x s_\theta(x_{t_i}, t_i)\Big) + \frac{1}{2} g(t_i)^2 \|s_\theta(x_{t_i}, t_i)\|^2$
**end for**
Compute $\log p_1(x_1) \sim \mathcal{N}(0, \sigma_1^2 I)$ ▷ Gaussian prior
$\log p_0(x_0) \leftarrow \log p_1(x_1) - \Delta \log p$
**return** $\log p_0(x_0)$

**Algorithm 2** Backward Euler Integration over Diffusion Trajectories

**Input:** Diffusion trajectory samples $\{(x_{t_0}, t_0), (x_{t_1}, t_1), \ldots, (x_{t_n}, t_n)\}$ where $t_i \in [0, 1], t_0 = 0, t_n = 1$; score model $s_\theta(x_t, t)$; noise schedule $g(t)$
**Output:** Log-likelihood estimate $\log p_0(x_0) = \log p_1(x_1) - \Delta \log p$
Initialize $\Delta \log p \leftarrow 0$
**for** $i = 0$ to $n - 1$ **do**
  $\Delta x_i \leftarrow x_{t_{i+1}} - x_{t_i}$
  $\Delta t_i \leftarrow t_{i+1} - t_i$
  $\Delta \log p \leftarrow \Delta \log p + s_\theta(x_{t_{i+1}}, t_{i+1}) \cdot \Delta x + \frac{1}{2} g(t_{i+1})^2 \mathrm{Tr}\Big(\nabla_x s_\theta(x_{t_{i+1}}, t_{i+1})\Big) + \frac{1}{2} g(t_{i+1})^2 \|s_\theta(x_{t_{i+1}}, t_{i+1})\|^2$
**end for**
Compute $\log p_1(x_1) \sim \mathcal{N}(0, \sigma_1^2 I)$ ▷ Gaussian prior
$\log p_0(x_0) \leftarrow \log p_1(x_1) - \Delta \log p$
**return** $\log p_0(x_0)$

**Algorithm 3** Trapezoidal Integration over Diffusion Trajectories

**Input:** Diffusion trajectory samples $\{(x_{t_0}, t_0), (x_{t_1}, t_1), \ldots, (x_{t_n}, t_n)\}$ where $t_i \in [0, 1], t_0 = 0, t_n = 1$; score model $s_\theta(x_t, t)$; noise schedule $g(t)$
**Output:** Log-likelihood estimate $\log p_0(x_0) = \log p_1(x_1) - \Delta \log p$
**function** INTEGRAND($x_t$, $t$,$\Delta x$, $\Delta t$)
  **return** $s_\theta(x_t, t) \cdot \Delta x + \frac{1}{2} g(t)^2 \mathrm{Tr}\Big(\nabla_x s_\theta(x_t, t)\Big)\Delta t + \frac{1}{2} g(t)^2 \|s_\theta(x_t, t)\|^2 \Delta t$
**end function**
Initialize $\Delta \log p \leftarrow 0$
**for** $i = 0$ to $n - 1$ **do**
  $\Delta x_i \leftarrow x_{t_{i+1}} - x_{t_i}$
  $\Delta t_i \leftarrow t_{i+1} - t_i$
  $f_a \leftarrow$ INTEGRAND$(x_{t_i}, t_i, \Delta x_i, \Delta t_i)$
  $f_b \leftarrow$ INTEGRAND$(x_{t_{i+1}}, t_{i+1}, \Delta x_i, \Delta t_i)$
  $\Delta \log p \leftarrow \Delta \log p + \frac{1}{2}(f_a + f_b)$
**end for**
Compute $\log p_1(x_1) \sim \mathcal{N}(0, \sigma_1^2 I)$ ▷ Gaussian prior
$\log p_0(x_0) \leftarrow \log p_1(x_1) - \Delta \log p$
**return** $\log p_0(x_0)$

**Algorithm 4** Interpolated Discrete Integration over Diffusion Paths

**Input:** Diffusion trajectory samples $\{(x_{t_0}, t_0), (x_{t_1}, t_1), \ldots, (x_{t_n}, t_n)\}$ where $t_i \in [0, 1], t_0 = 0, t_n = 1$; score model $s_\theta(x_t, t)$; noise schedule $g(t)$; Interpolation factor $m$, Discrete integration function $F(\{(x_{t_0}, t_0), \ldots\}, s_\theta(x_t, t), g(t))$
**Output:** Log-likelihood $\log p_0(x_0)$
$s_{\text{interp}} \leftarrow \{(x_{t_0}, t_0)\}$ ▷ Initialize interpolated trajectory
**for** $i = 0$ to $n - 1$ **do**
    **for** $k = 1$ to $m$ **do**
        $\alpha \leftarrow k/m$
        $x \leftarrow \alpha x_{t_i} + (1 - \alpha) x_{t_{i+1}}$
        $t \leftarrow \alpha t_i + (1 - \alpha) t_{i+1}$
        APPEND$(s_{\text{interp}}, (x, t))$
    **end for**
**end for**
**return** $F(s_{\text{interp}}, s_\theta(x_t, t), g(t))$ ▷ Perform discrete integration (trapezoidal, euler, etc) on interpolated path

**Algorithm 5** Piecewise ODE Integration over Diffusion Paths

**Input:** Diffusion trajectory samples $\{(x_{t_0}, t_0), (x_{t_1}, t_1), \ldots, (x_{t_n}, t_n)\}$ where $t_i \in [0, 1], t_0 = 0, t_n = 1$; score model $s_\theta(x_t, t)$; noise schedule $g(t)$
**Output:** Log-likelihood $\log p_0(x_0)$
**function** INTEGRAND$(x_t, t, \frac{dx}{dt})$
    **return** $s_\theta(x_t, t) \cdot \frac{dx}{dt} + \frac{1}{2} g(t)^2 \mathrm{Tr}\Big(\nabla_x s_\theta(x_t, t)\Big) + \frac{1}{2} g(t)^2 \|s_\theta(x_t, t)\|^2$
**end function**
Initialize $\Delta \log p \leftarrow 0$
**for** $i = 0$ to $n - 1$ **do**
    $\frac{dx}{dt} \leftarrow \frac{x_{t_{i+1}} - x_{t_i}}{t_{i+1} - t_i}$
    Solve the following ODEs from $t = 0$ to $t = 1$ using an ODE solver:

- $\frac{dx_t}{dt} = \frac{dx}{dt}$ with initial condition $x_0$
- $\frac{d(\mathtt{logp})}{dt} = $ INTEGRAND$(x_t, t, \frac{dx}{dt})$ with initial value 0
- **return** $\{x_t|_{t=0}^{1}\}, \Delta_i \log p$

    $\Delta \log p \leftarrow \Delta \log p + \Delta_i \log p$
**end for**
Compute $\log p_1(x_1) \sim \mathcal{N}(0, \sigma_1^2 I)$ ▷ Gaussian prior
$\log p_0(x_0) \leftarrow \log p_1(x_1) - \Delta \log p$
**return** $\log p_0(x_0)$

**Algorithm 6** ODE Integration over Flow Paths

**Input:** Diffusion sample $x_0$; score model $s_\theta(x_t, t)$; noise schedule $g(t)$
**Output:** Log-likelihood $\log p_0(x_0)$
**function** DRIFT$(x_t, t)$
    **return** $-g(t)^2 \cdot s_\theta(x_t, t)$
**end function**
**function** TRACEJACOBIAN$(x_t, t)$
    **return** $\frac{1}{2} g(t)^2 \mathrm{Tr}\Big(\nabla_x s_\theta(x_t, t)\Big)$
**end function**
Solve the following ODEs from $t = 0$ to $t = 1$ using an ODE solver:

- $\frac{dx_t}{dt} = $ DRIFT$(x_t, t)$ with initial condition $x_0$
- $\frac{d(\mathtt{logp})}{dt} = $ TRACEJACOBIAN$(x_t, t)$ with initial value 0
- **return** $\{x_t|_{t=0}^{1}\}, \Delta \log p$

Compute $\log p_1(x_1) \sim \mathcal{N}(0, \sigma_1^2 I)$ ▷ Gaussian prior
$\log p_0(x_0) \leftarrow \log p_1(x_1) - \Delta \log p$
**return** $\log p_0(x_0)$

#### F. Analysis of Numerical Integration Techniques

Here we present the accuracy of likelihood recovery on discrete diffusion trajectories by various formulations of numerical integration. **Supp. Fig. S1** summarizes the results on our trimodal Gaussian mixture toy model. The simplest algorithms for discrete integration are the forward (**Algorithm 1**) and backwards (**Algorithm 2**) Euler methods, which compute eq. (26) using the score computed at either the beginning or the end respectively of each diffusion step. Forward Euler integration (**Supp. Fig. S1a**) consistently underestimates the true probability, and backward Euler integration (**Supp. Fig. S1c**) consistently overestimates it.

One potential explanation of this error is that by taking large discrete jumps, the diffusion trajectory skips over regions of significant and relevant probability change without sampling their gradients. To explore this, we linearly interpolated diffusion trajectories by adding 9 additional points per step, spread equally in space and time between each pair $(x_i, t_i), (x_{t_{i+1}}, t_{i+1})$ of the original trajectory (**Algorithm 4**). **Supp. Fig. S1b** and **Supp. Fig. S1d** integrate these interpolated trajectories using the forwards and backward Euler algorithms respectively, and demonstrate significantly improved likelihood recovery. However, even with interpolation, forward Euler consistently underestimates likelihoods, and backward Euler consistently overestimates.

Trapezoidal integration (**Algorithm 3**, **Supp. Fig. S1e**) provides a significantly less biased and less noisy estimate of the true distribution without sampling at additional points. Interestingly, unlike the Euler integration methods, interpolation of trapezoidal-integrated paths only marginally increases accuracy and remains nearly as noisy, especially compared to flow trajectories (**Supp. Fig. S1f**).

As trapezoidal integration and interpolation are still coarse approximations, we further tested whether using a black-box integrator capable of high-order approximations and intelligent subsampling could allow us to attain a similar confidence in our integration as flow trajectories. These solvers require an integrand indexed by a continuous parameter $t$, while our discrete diffusion path is defined only at discrete points $t_i$; this is easy enough to remedy by connecting each pair of points with a straight line, defining a continuous path akin to that sampled by interpolating. A larger issue is that our continuous integrand, eq. 26, depends on $\frac{\partial x_t}{\partial t}$, requiring the path to be differentiable; however, this straight-line path is not differentiable at its vertices. While it is possible to define a smooth path through the discrete trajectory using splines, this is not necessary, as a function does not need to be defined everywhere to be integrable so long as it is only missing countably many points. That is, we only need to integrate along the countably many differentiable sections of the path - each straight line segment - and sum the resulting $\Delta_i \log p$ values:

$$\log p_0(x_0) = \log p_1(x_1) - \sum_{i=0}^{n-1} \int_{t_i}^{t_{i+1}} \left( s_\theta(x_t, t) \cdot \frac{\Delta x_{t_i}}{\Delta t_i} + \frac{1}{2} g(t)^2 \mathrm{Tr}\Big(\nabla_x s_\theta(x_t, t)\Big) + \frac{1}{2} g(t)^2 \|s_\theta(x_t, t)\|^2 \right) dt, \tag{S-14}$$

where $\Delta x_{t_i} = x_{t_{i+1}} - x_{t_i}$ and $\Delta t_i = t_{i+1} - t_i$ as in eq. (28). This integral is also described in **Algorithm 5**.

Piecewise integration elevates the accuracy of integrating arbitrary discrete trajectories like diffusion to that of integrating ODE trajectories like flow. However, when we apply it to diffusion trajectories from our Gaussian toy model, the likelihood recovery is essentially unchanged from basic trapezoidal integration (**Supp. Fig. S1g**). We can measure this convergence quantitatively by plotting the per-sample correlation between likelihoods from pairs of integration methods (**Supp. Fig. S1i-m**), which confirms that trapezoidal integration of toy model diffusion trajectories attains similar numerical precision to interpolated trapezoidal and piecewise ODE integration.

We also investigate the numerical accuracy of our DFMDock integration methods. Although we have no ground truth distribution with which to compare recovered likelihoods, we can again use per-sample correlation between integration methods which we know to have progressively better precision to measure convergence towards some consensus value. For this analysis, we combined the results from 120 DFMDock generated poses for each of the 25 complexes in our validation set. We started with flow trajectory ODE integration, as again, we have no ground truth with which to verify like we did for our Gaussian flow trajectories. For ODE integration, we used the Runge-Kutta integration method of order 4 and the $\frac{3}{8}$ rule (see **Section VI.C**), which uses a user-defined step size and subsamples each step by a fixed amount. A simple choice of step size is $\frac{1}{40}$, which produces 40 steps from $t = 0$ to $t = 1$, matching the number of diffusion steps taken during model inference. Increase precision to 80 steps yielded no significant difference in recovered energy $-\log p_0$ (**Supp. Fig. S2a**), though there are a small number of outliers which we believe stem from dynamically-generated flow paths occasionally following a different trajectory with the higher precision. We therefore used 40-step ODE integration for integration over DFMDock flow trajectories in our calculations.

We performed a similar analysis for DFMDock diffusion trajectory integration, both by discrete methods and piecewise ODE methods. We compared each method in turn with the highest precision method we tested: piecewise ODE integration with 120 steps (**Supp. Fig. S2b-e**). As diffusion trajectories have 40 steps, a step size of $\frac{1}{120}$ means each short segment has 3 total steps, 80 steps means 2 ODE steps per diffusion step, and 40 steps means only a single ODE step each. Unlike the Gaussian trajectories, the correlation between trapezoidal integration and piecewise ODE integration displays significant noise (**Supp. Fig. S2b**), motivating the use of higher precision methods. Interpolated trapezoidal integration aligns much better with piecewise integration (**Supp. Fig. S2c**). However, the computational cost of increased accuracy through interpolation grows more quickly than ODE integration due to the lower accuracy of the trapezoidal integration method. 40-step piecewise ODE integration achieves slightly worse 120-step piecewise ODE correlation than interpolated trapezoidal, but at a fraction of the computational cost, needing roughly $\frac{1}{3}$ the compute time (**Supp. Fig. S2d**). Increasing to 80 steps marginally increases precision as well (**Supp. Fig. S2e**). We plotted learned energies from trapezoidal, interpolated trapezoidal, and 40-step piecewise ODE integration against DockQ, interface RMSD, and Rosetta interface energy (**Supp. Fig. S4-S9, S15-S9**) and observed a marked decrease in noise by interpolated trapezoidal and piecewise ODE integration over standard trapezoidal, but little difference between interpolated trapezoidal and piecewise ODE integration. We therefore used 40-step piecewise ODE integration for our DFMDock diffusion trajectory results.

With these experiments, we can be reasonably confident that our integration methods achieve sufficient numerical accuracy on diffusion trajectories for both of our models. Despite this, we still find diffusion trajectory likelihoods to be much noisier than flow trajectory likelihoods. If neither numerical precision nor sampling resolution of the score is responsible for the significant noise compared to flow, where does it come from? We elaborate on one source in **Supplementary Material B** and discuss other sources in **Section V**.

### G. Additional Supplementary Figures

| **Supp. Fig.** | **Description** |
|---|---|
| **Supp. Fig. S1** | Trimodal Gaussian Recovered Probabilities by Numerical Integration Methods |
| **Supp. Fig. S2** | Comparison of DFMDock Numerical Integration Methods |
| **Supp. Fig. S3** | Noise Schedule and Assumed Prior Effects on Recovered Likelihoods |

| **Supp. Fig.** | $y$**-axis** | $x$**-axis** |
|---|---|---|
| **Supp. Fig. S4** | Diffusion Trajectory Learned Energy (Trapezoidal Integration) | Interface RMSD |
| **Supp. Fig. S5** | Diffusion Trajectory Learned Energy (Trapezoidal Integration) | DockQ |
| **Supp. Fig. S6** | Diffusion Trajectory Learned Energy (Interpolated Trapezoidal Integration) | Interface RMSD |
| **Supp. Fig. S7** | Diffusion Trajectory Learned Energy (Interpolated Trapezoidal Integration) | DockQ |
| **Supp. Fig. S8** | Diffusion Trajectory Learned Energy (Piecewise ODE Integration) | Interface RMSD |
| **Supp. Fig. S9** | Diffusion Trajectory Learned Energy (Piecewise ODE Integration) | DockQ |
| **Supp. Fig. S10** | Flow Trajectory Learned Energy | Interface RMSD |
| **Supp. Fig. S11** | Flow Trajectory Learned Energy | DockQ |
| **Supp. Fig. S12** | Rosetta Energy | Interface RMSD |
| **Supp. Fig. S13** | Rosetta Energy | DockQ |
| **Supp. Fig. S14** | Flow Trajectory Learned Energy | Rosetta Energy |
| **Supp. Fig. S15** | Diffusion Trajectory Learned Energy (Trapezoidal Integration) | Rosetta Energy |
| **Supp. Fig. S16** | Diffusion Trajectory Learned Energy (Interpolated Trapezoidal Integration) | Rosetta Energy |
| **Supp. Fig. S17** | Diffusion Trajectory Learned Energy (Piecewise ODE Integration) | Rosetta Energy |
| **Supp. Fig. S18** | Diffusion Trajectory Learned Energy (Trapezoidal Integration) | Flow Trajectory Learned Energy |
| **Supp. Fig. S19** | Diffusion Trajectory Learned Energy (Interpolated Trapezoidal Integration) | Flow Trajectory Learned Energy |
| **Supp. Fig. S20** | Diffusion Trajectory Learned Energy (Piecewise ODE Integration) | Flow Trajectory Learned Energy |

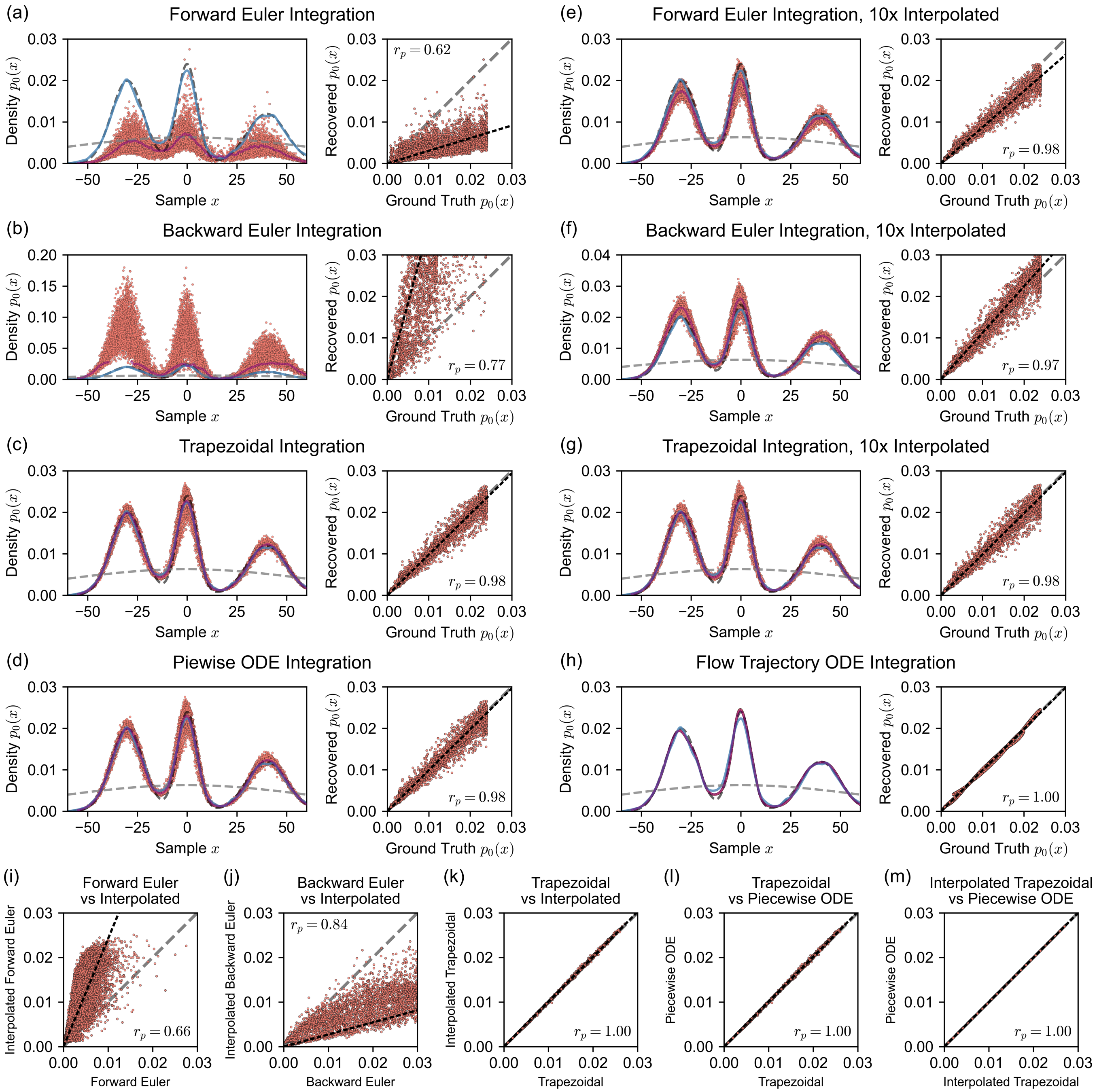


FIG. S1. Recovered probabilities $p_0(x_0)$ from our toy 1D Gaussian mixture model by integrating diffusion trajectories with different numerical integration methods. a) Forward Euler integration, b) interpolated forward Euler integration, c) backward Euler integration, d) interpolated backward Euler integration, e) trapezoidal integration, f) interpolated trapezoidal integration, and f) piecewise ODE integration. Interpolated integration (panels b, d, and f) was performed by first adding 9 additional points linearly spaced along each step of the original trajectory (thus multiplying the number of steps tenfold) and integrating the resulting path by the corresponding method. Likelihood recovery using ODE integration over flow trajectories (h) is included as a reference. i-m) direct per-sample comparison of learned energies recovered by various methods; i-k) display forward Euler, backward Euler, and trapezoidal integration on the $x$ axis plotted against the interpolated variant of each method on the $y$ axis. l) Trapezoidal integration versus piecewise ODE integration; m) interpolated trapezoidal integration versus piecewise ODE integration.

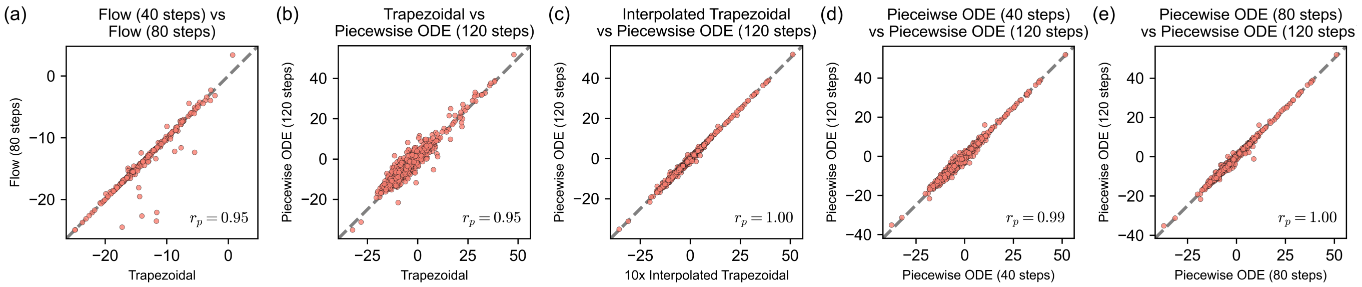


FIG. S2. Comparison of DFMDock integration methods. Plots show per-sample correlation of DFMDock learned negative log-likelihood. a) Flow ODE integration with 40 steps vs 80 steps. b-e) Diffusion trajectory integration by: b) trapezoidal vs 120-step piecewise ODE integration, c) interpolated trapezoidal vs 120-step piecewise ODE integration, d) 40-step piecewise ODE vs 120-step piecewise ODE integration, and e) 80-step piecewise ODE vs 120-step piecewise ODE integration.

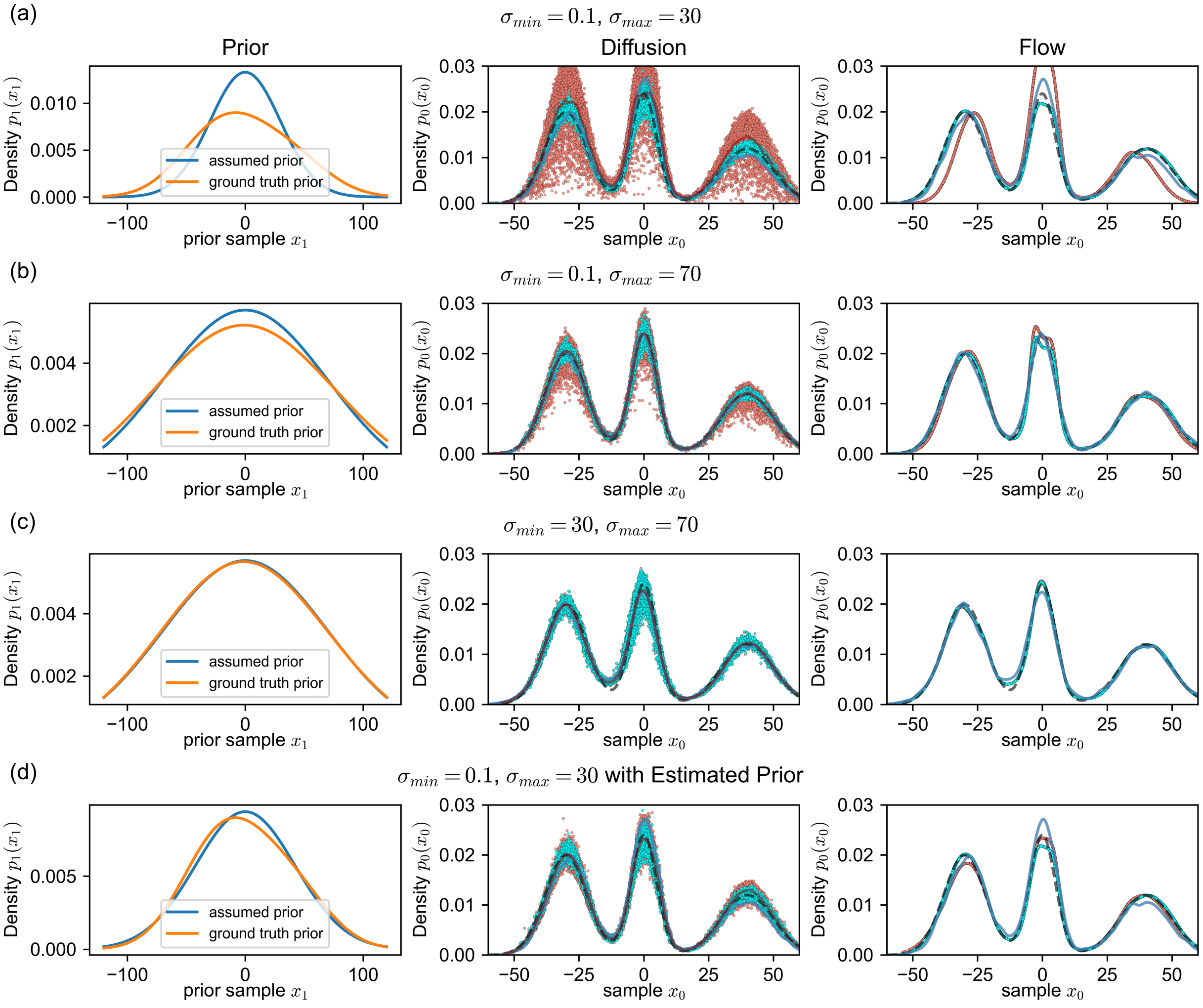


FIG. S3. Comparison of prior accuracy and likelihood recovery with different noise schedule hyperparameters $\sigma_0$ and $\sigma_1$. Row a), $\sigma_0 = 0.1$, $\sigma_1 = 30$; row b, $\sigma_0 = 0.1$, $\sigma_{=}70$; row c), $\sigma_0 = 30$, $\sigma_1 = 70$; row d), $\sigma_0 = 0.1$, $\sigma_1 = 30$, with estimated data deviation of $\sigma_{\text{data}} = 29.975$. The left panel in each row plots the assumed prior (blue) vs the ground truth prior (orange): for rows a-c, $p_1(x_1) = \mathcal{N}(x_1; \mu = 0, \sigma = \sigma_1)$; for row d, $p_1(x_1) = \mathcal{N}\left(x_1; \mu = 0, \sigma = \sqrt{\sigma_{\text{data}}^2 + (\sigma_1^2 - \sigma_0^2)}\right)$. The ground truth marginals (orange line) in rows a) and d) are identical in value with different axis scaling. The right two panels in each row plot recovered likelihoods along diffusion and flow trajectories respectively. As in **Fig. 3**, the black dashed line is the ground truth data distribution and the dark blue line is the sample KDE. Red dots represent recovered likelihoods from the assumed prior, $p_0(x_0) = p_1^{Assumed}(x_1) + \Delta \log p$, and cyan dots represent recovered likelihoods from the ground truth prior, $p_0(x_0) = p(x_1, 1) + \Delta \log p$.

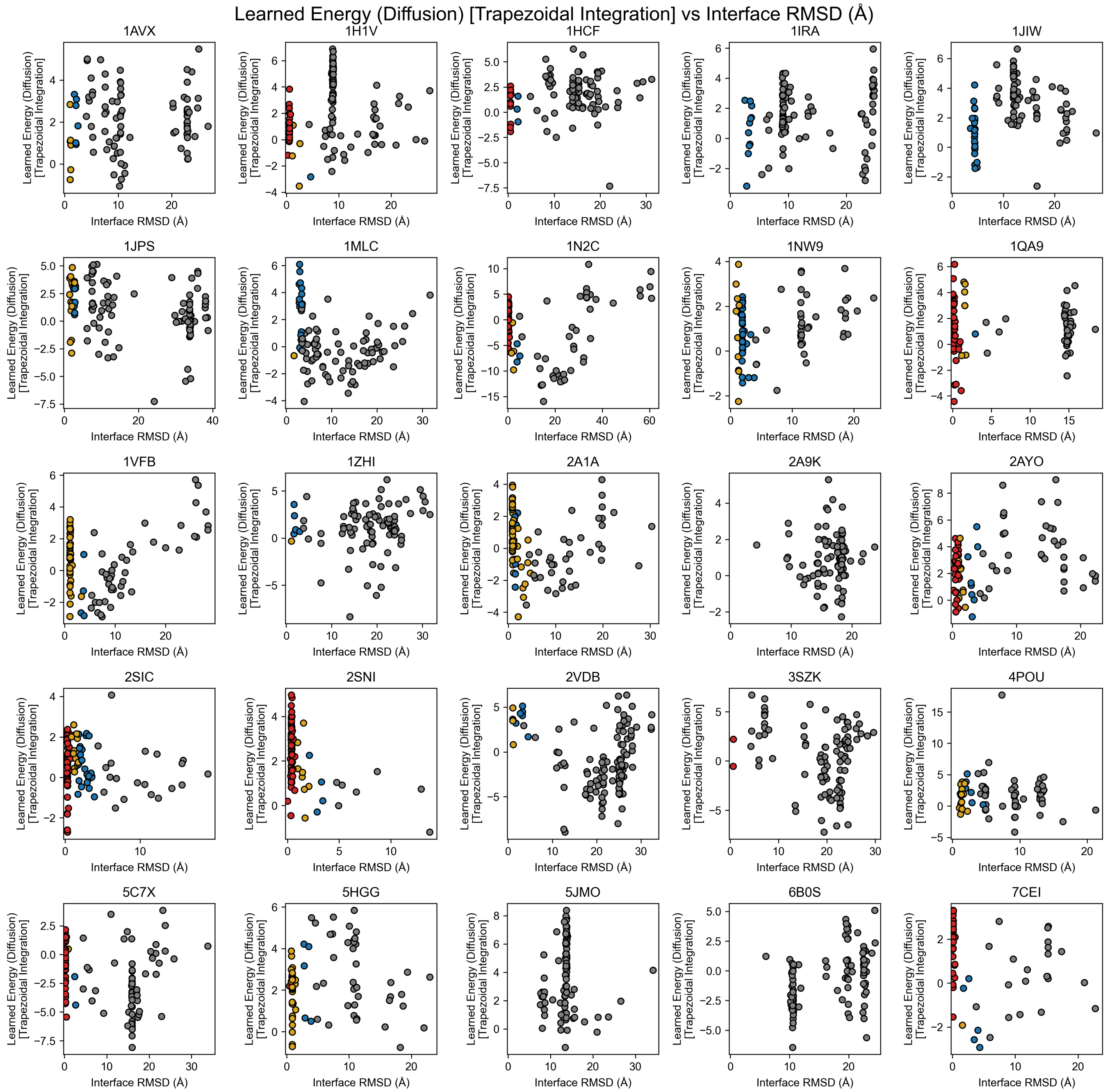


FIG. S4. Learned energy computed from integrating over diffusion trajectories with the trapezoidal method, plotted against interface RMSD of 120 docking poses generated from DFMDock for 25 targets in the DB5.5 dataset. Individual points are colored by their docking quality based on the CAPRI classification (incorrect: gray, acceptable: blue, medium: gold, high: red).

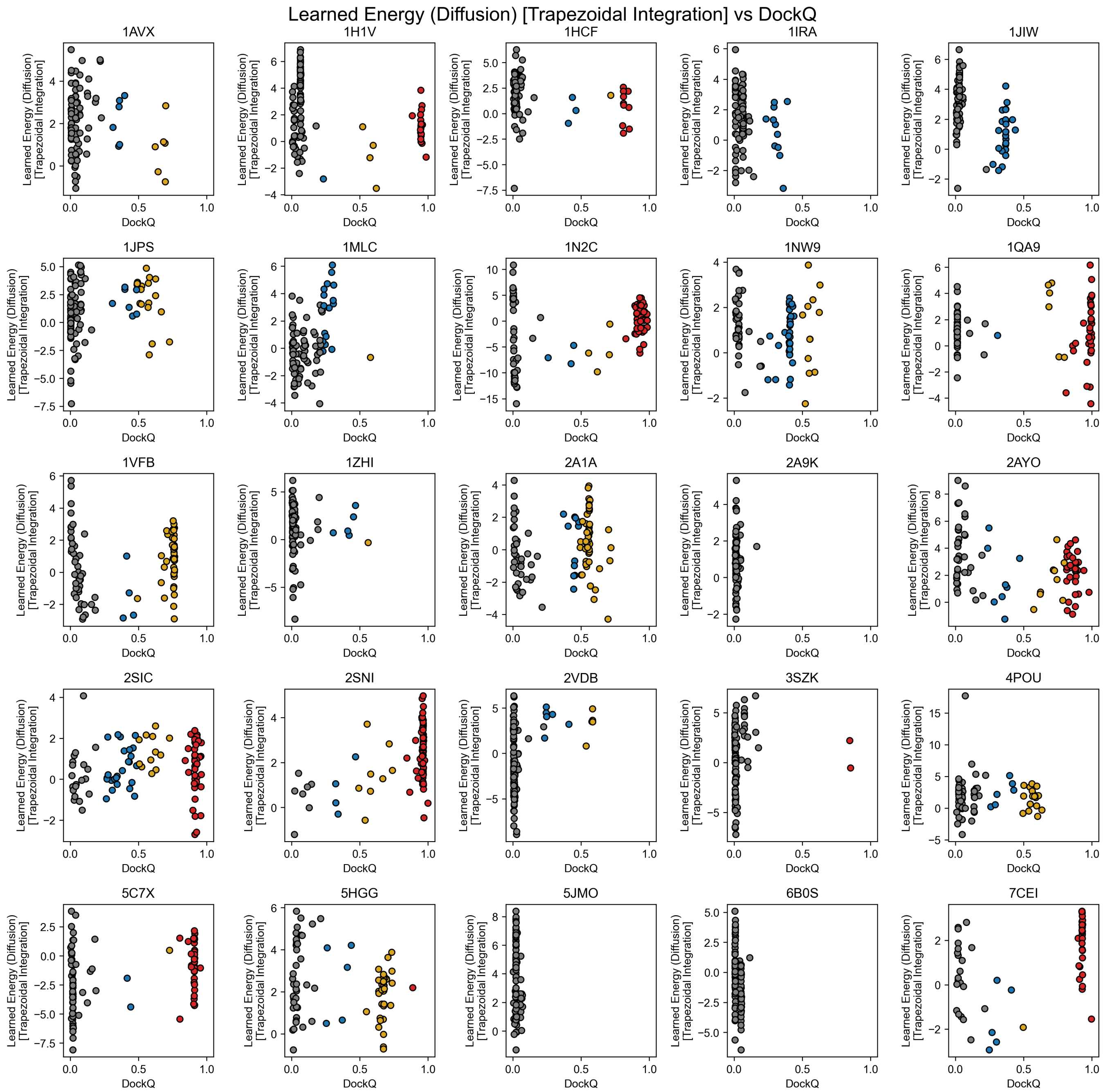


FIG. S5. Learned energy computed from integrating over diffusion trajectories with the trapezoidal method, plotted against DockQ of 120 docking poses generated from DFMDock for 25 targets in the DB5.5 dataset. Individual points are colored by their docking quality based on the CAPRI classification (incorrect: gray, acceptable: blue, medium: gold, high: red).

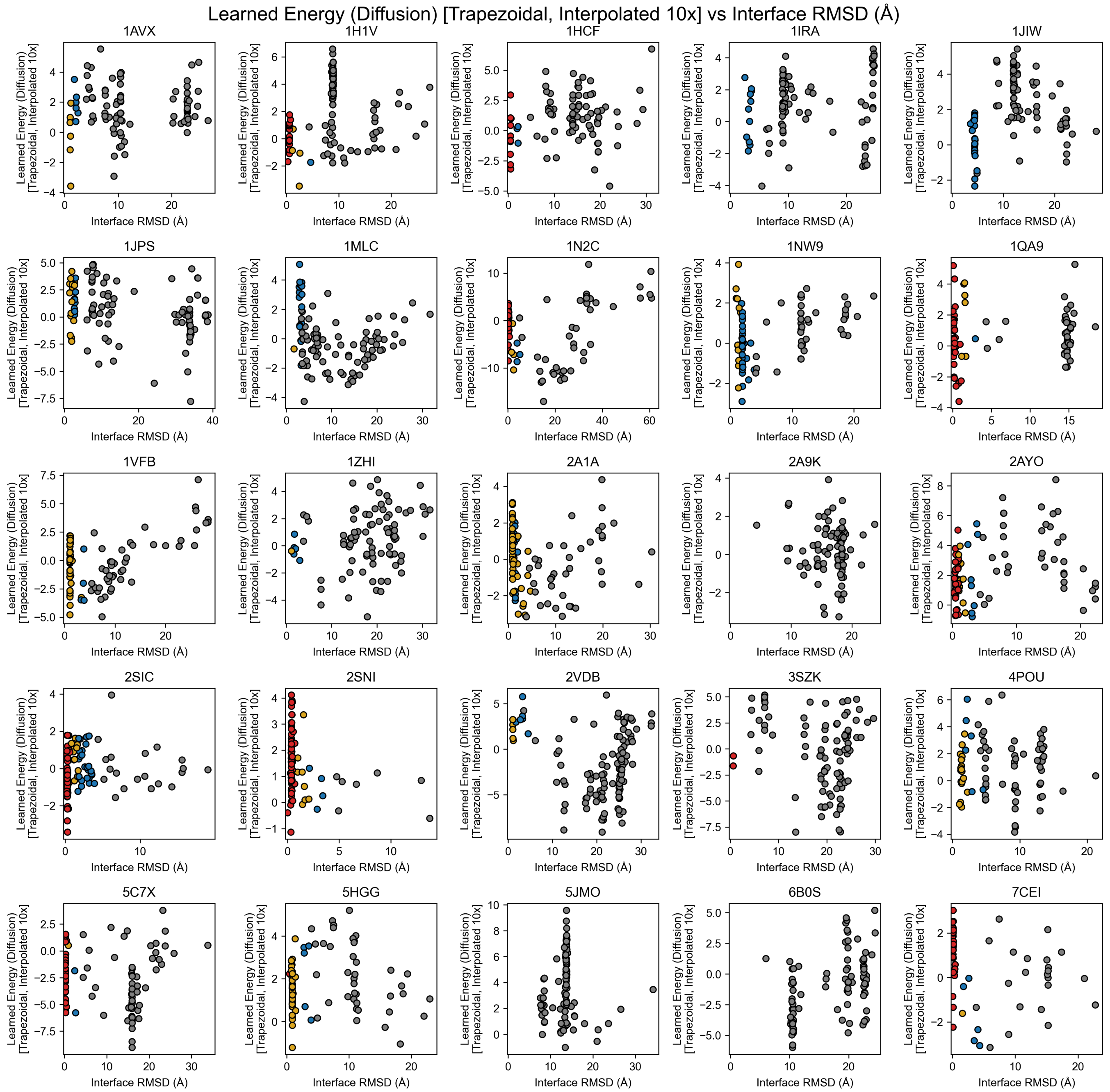


FIG. S6. Learned energy computed from integrating over diffusion trajectories with the interpolated trapezoidal method, plotted against interface RMSD of 120 docking poses generated from DFMDock for 25 targets in the DB5.5 dataset. Individual points are colored by their docking quality based on the CAPRI classification (incorrect: gray, acceptable: blue, medium: gold, high: red).

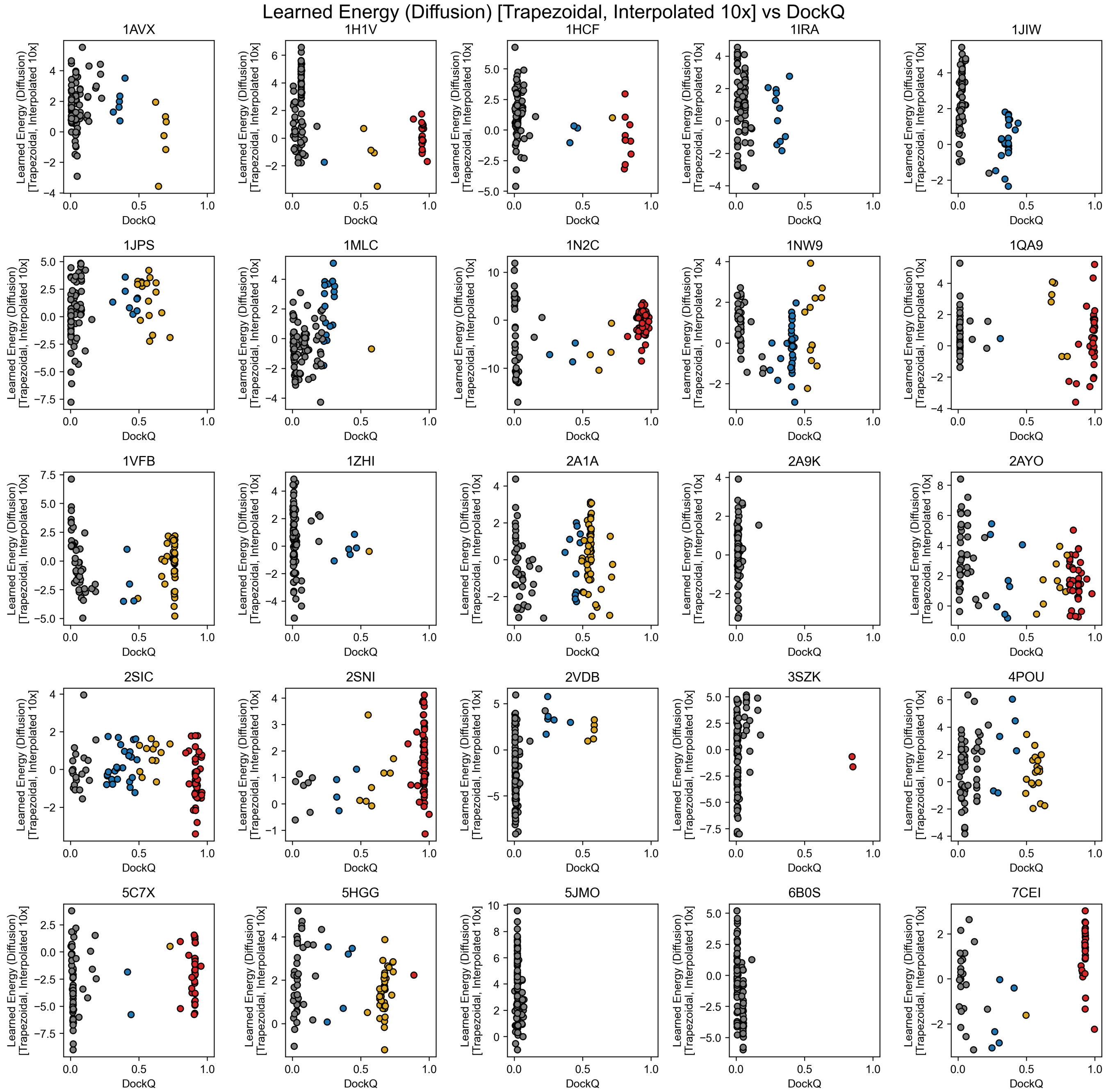


FIG. S7. Learned energy computed from integrating over diffusion trajectories with the interpolated trapezoidal method, plotted against DockQ of 120 docking poses generated from DFMDock for 25 targets in the DB5.5 dataset. Individual points are colored by their docking quality based on the CAPRI classification (incorrect: gray, acceptable: blue, medium: gold, high: red).

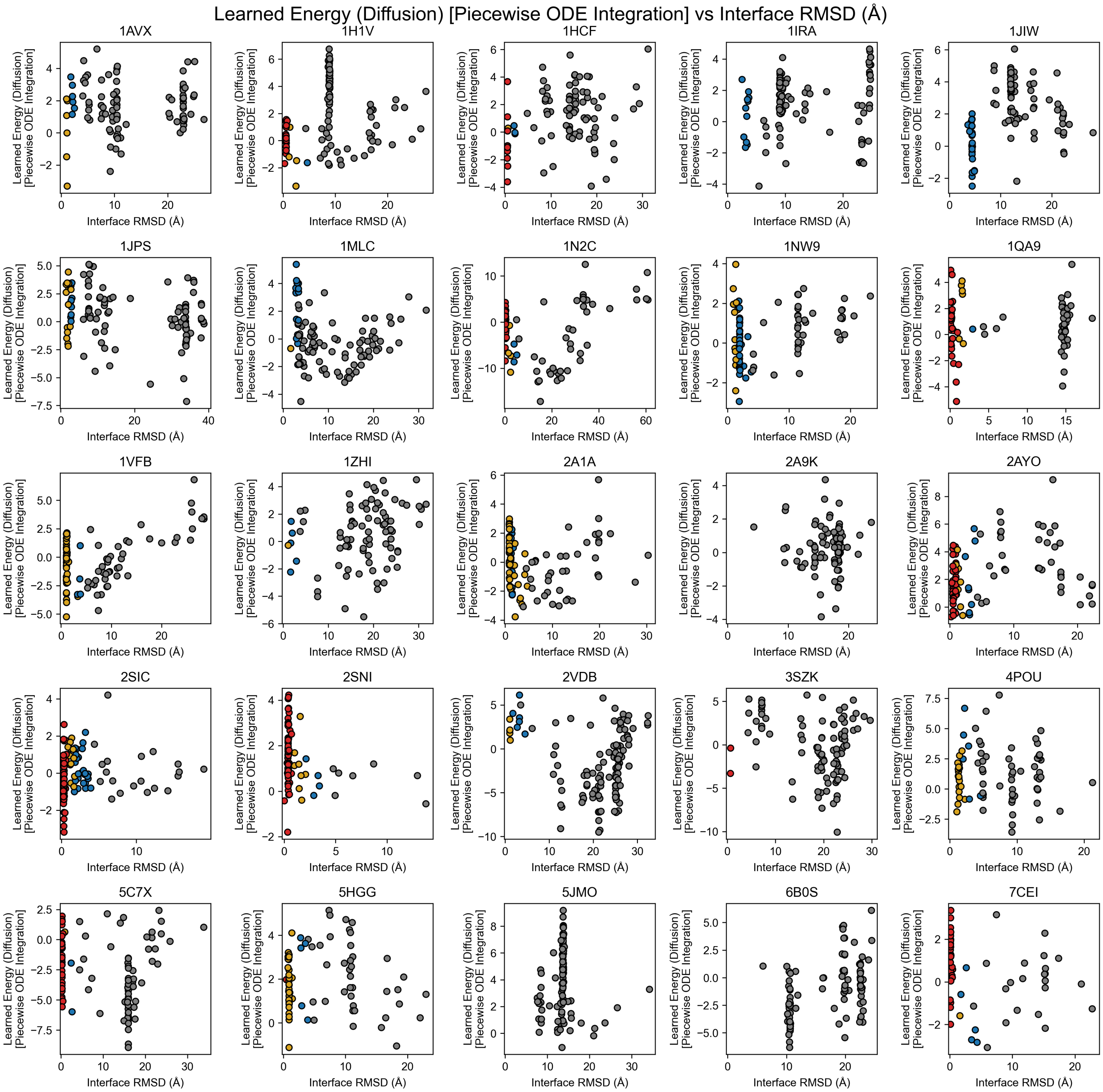


FIG. S8. Learned energy computed from integrating over diffusion trajectories with the piecewise ODE method, plotted against interface RMSD of 120 docking poses generated from DFMDock for 25 targets in the DB5.5 dataset. Individual points are colored by their docking quality based on the CAPRI classification (incorrect: gray, acceptable: blue, medium: gold, high: red).

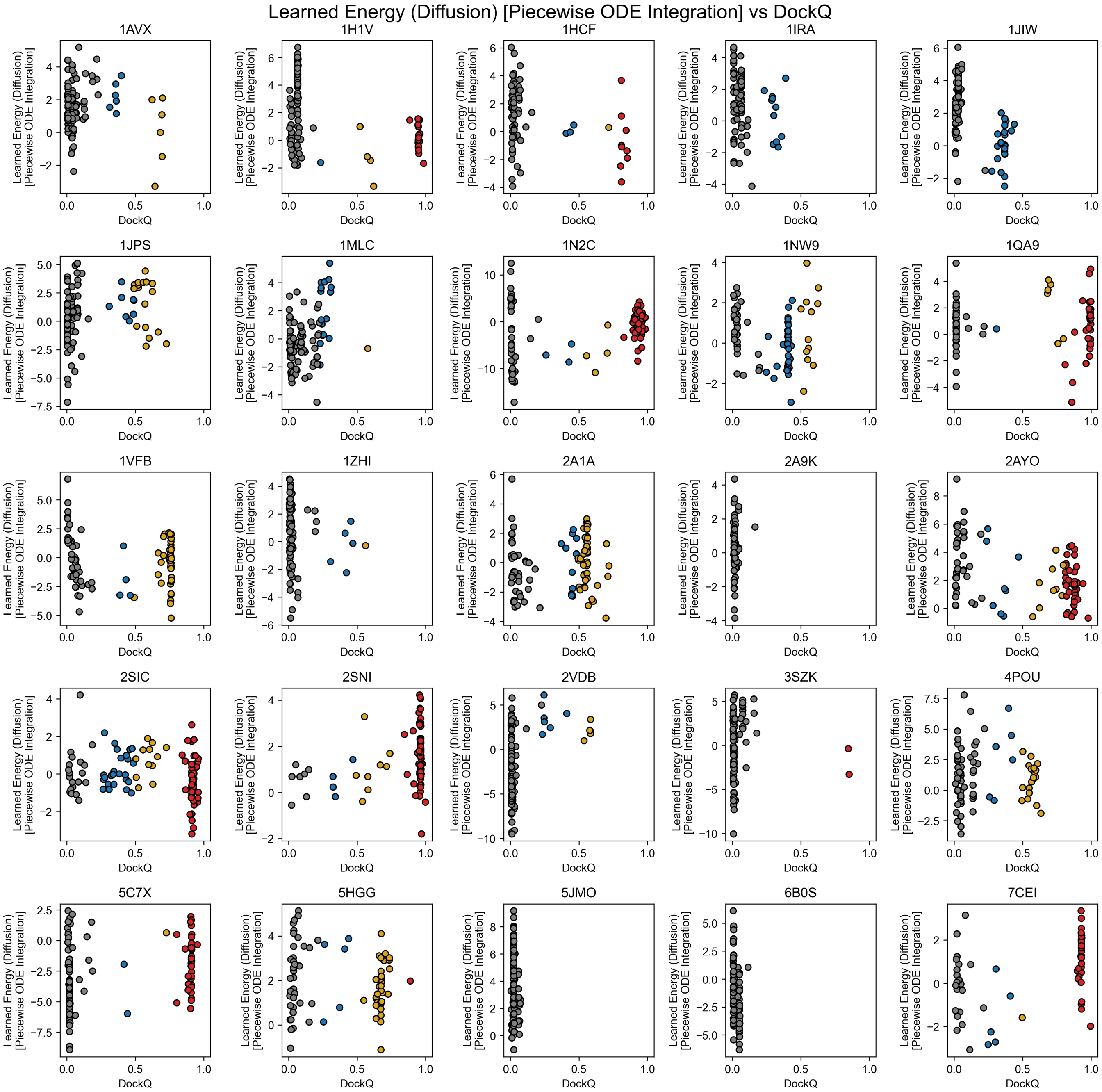


FIG. S9. Learned energy computed from integrating over diffusion trajectories with the piecewise ODE method, plotted against DockQ of 120 docking poses generated from DFMDock for 25 targets in the DB5.5 dataset. Individual points are colored by their docking quality based on the CAPRI classification (incorrect: gray, acceptable: blue, medium: gold, high: red).

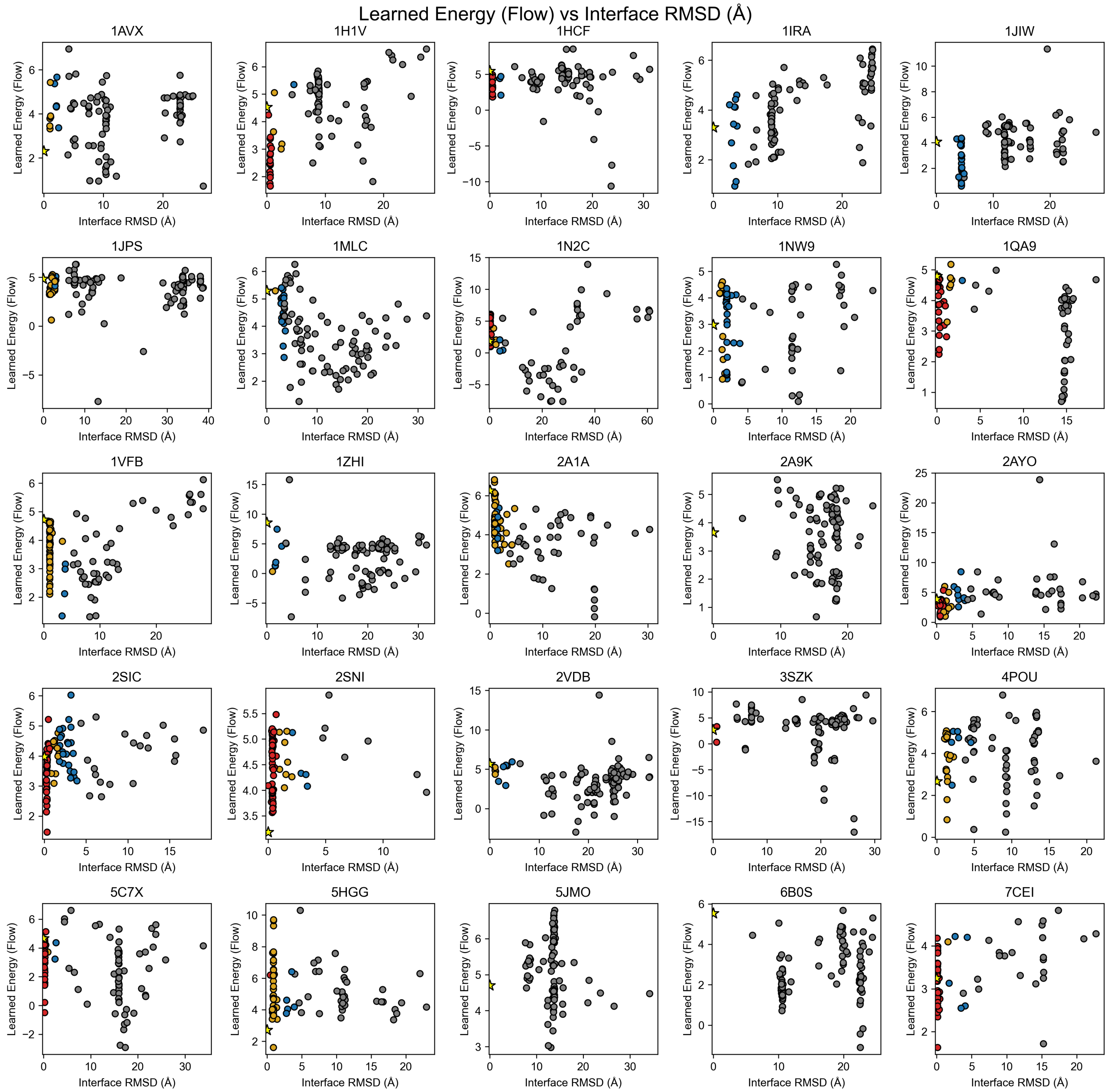


FIG. S10. Learned energy computed from integrating over flow trajectories, plotted against interface RMSD of 120 docking poses generated from DFMDock for 25 targets in the DB5.5 dataset. Individual points are colored by their docking quality based on the CAPRI classification (incorrect: gray, acceptable: blue, medium: gold, high: red). Energy of the ground truth structure is shown as a yellow star.

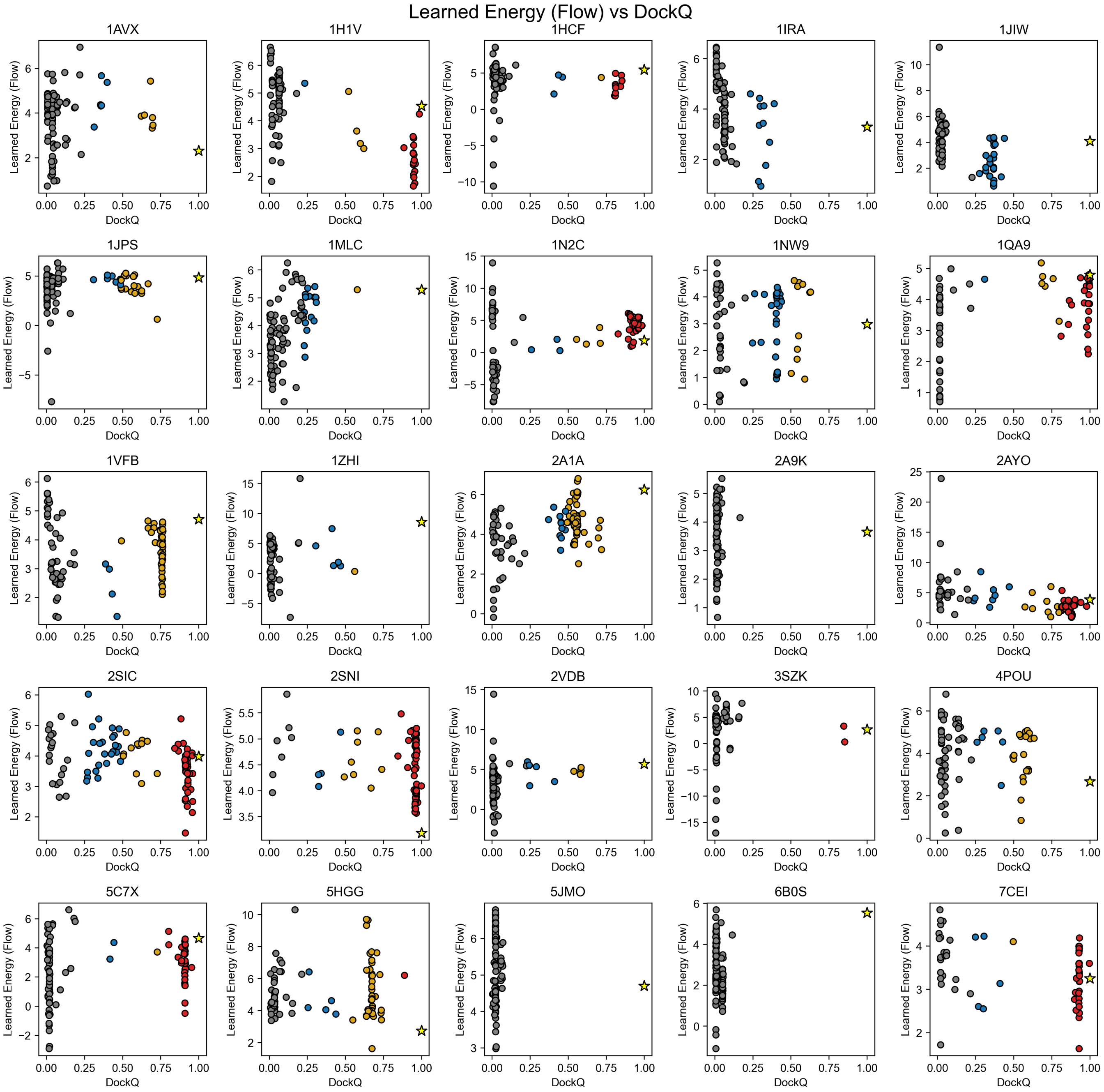


FIG. S11. Learned energy computed from integrating over flow trajectories, plotted against DockQ of 120 docking poses generated from DFMDock for 25 targets in the DB5.5 dataset. Individual points are colored by their docking quality based on the CAPRI classification (incorrect: gray, acceptable: blue, medium: gold, high: red). Energy of the ground truth structure is shown as a yellow star.

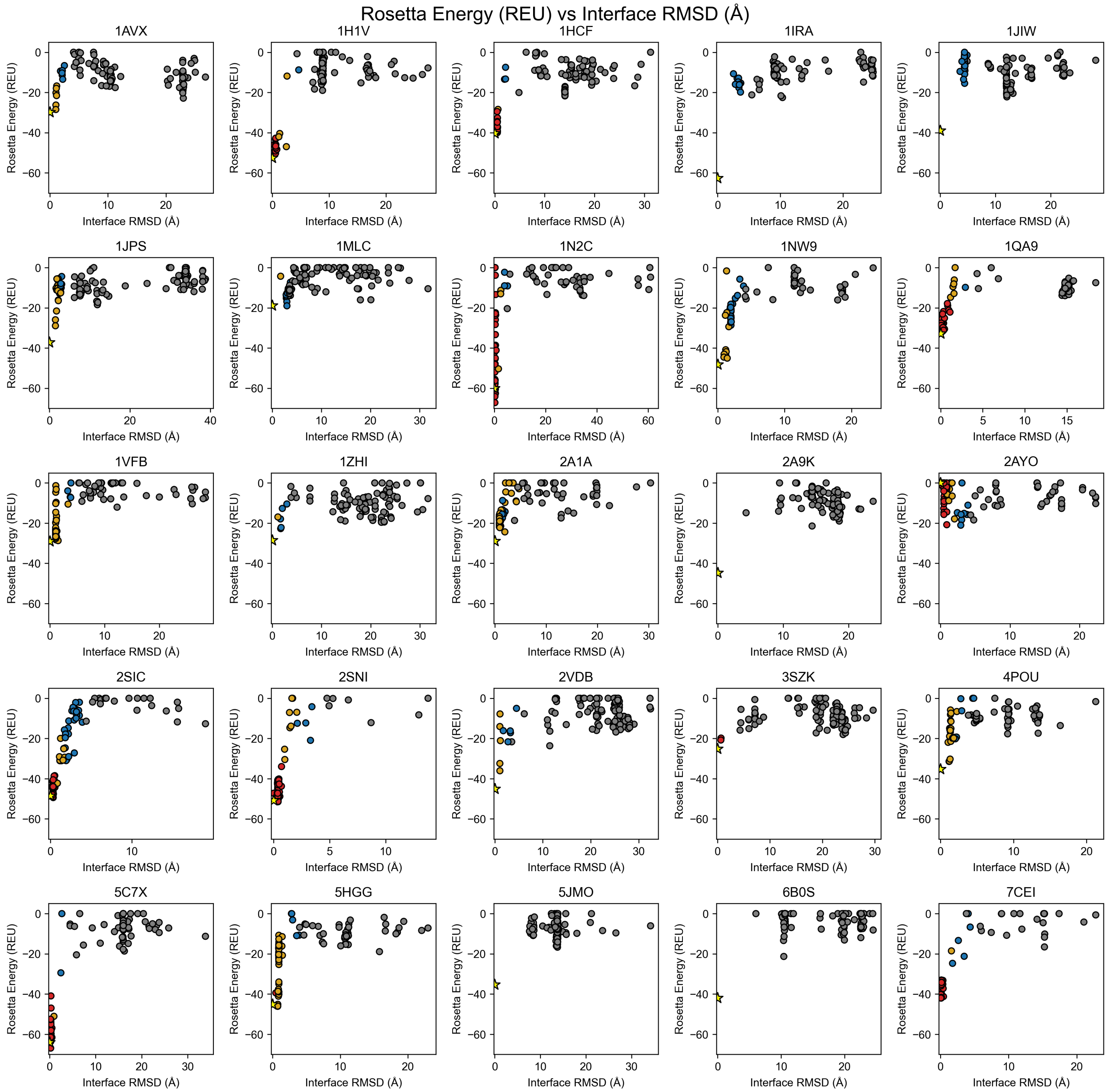


FIG. S12. Rosetta energy plotted against interface RMSD of 120 docking poses generated from DFMDock for 25 targets in the DB5.5 dataset. Individual points are colored by their docking quality based on the CAPRI classification (incorrect: gray, acceptable: blue, medium: gold, high: red). Energy of the ground truth structure is shown as a yellow star.

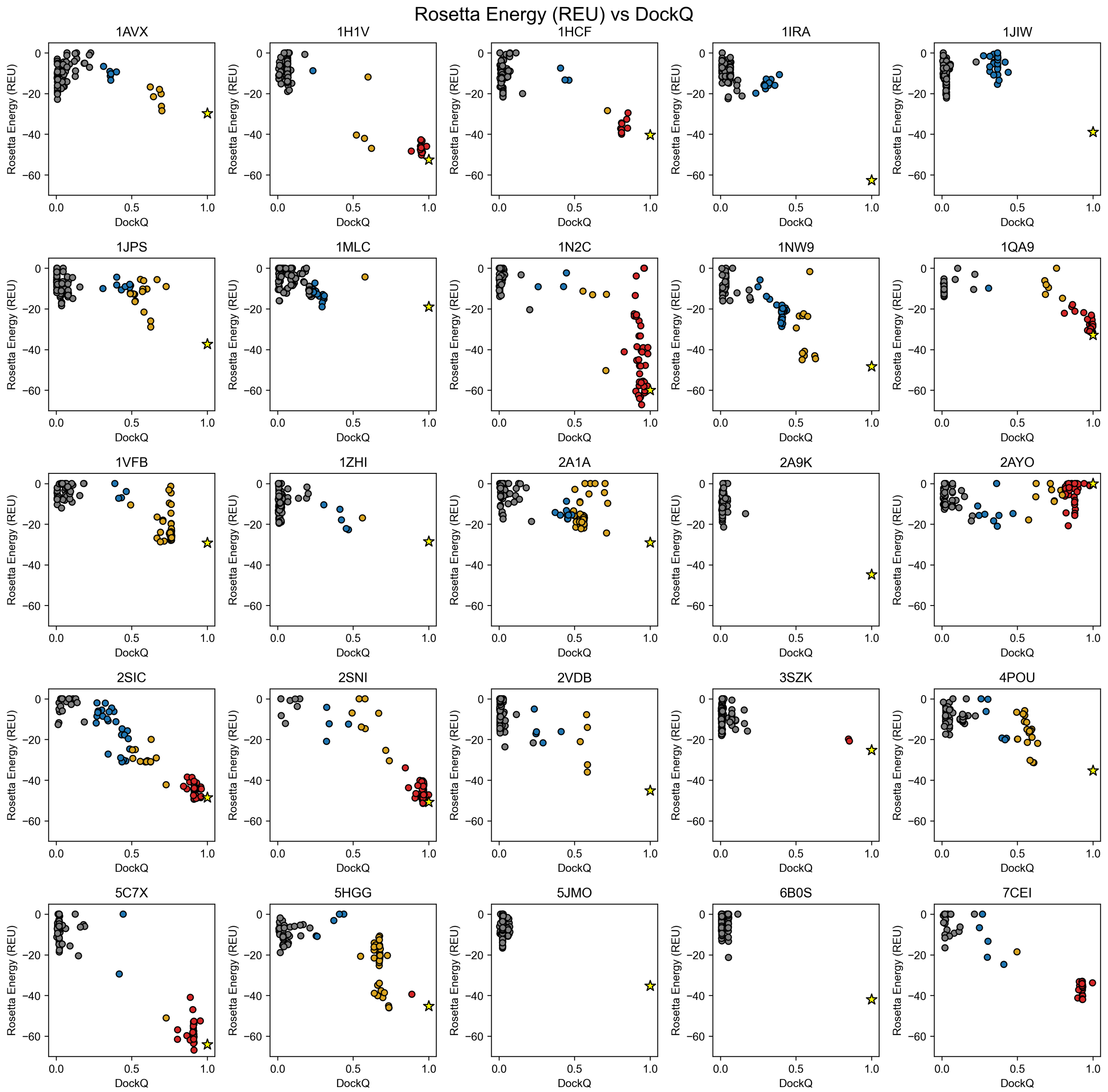


FIG. S13. Rosetta energy plotted against DockQ of 120 docking poses generated from DFMDock for 25 targets in the DB5.5 dataset. Individual points are colored by their docking quality based on the CAPRI classification (incorrect: gray, acceptable: blue, medium: gold, high: red). Energy of the ground truth structure is shown as a yellow star.

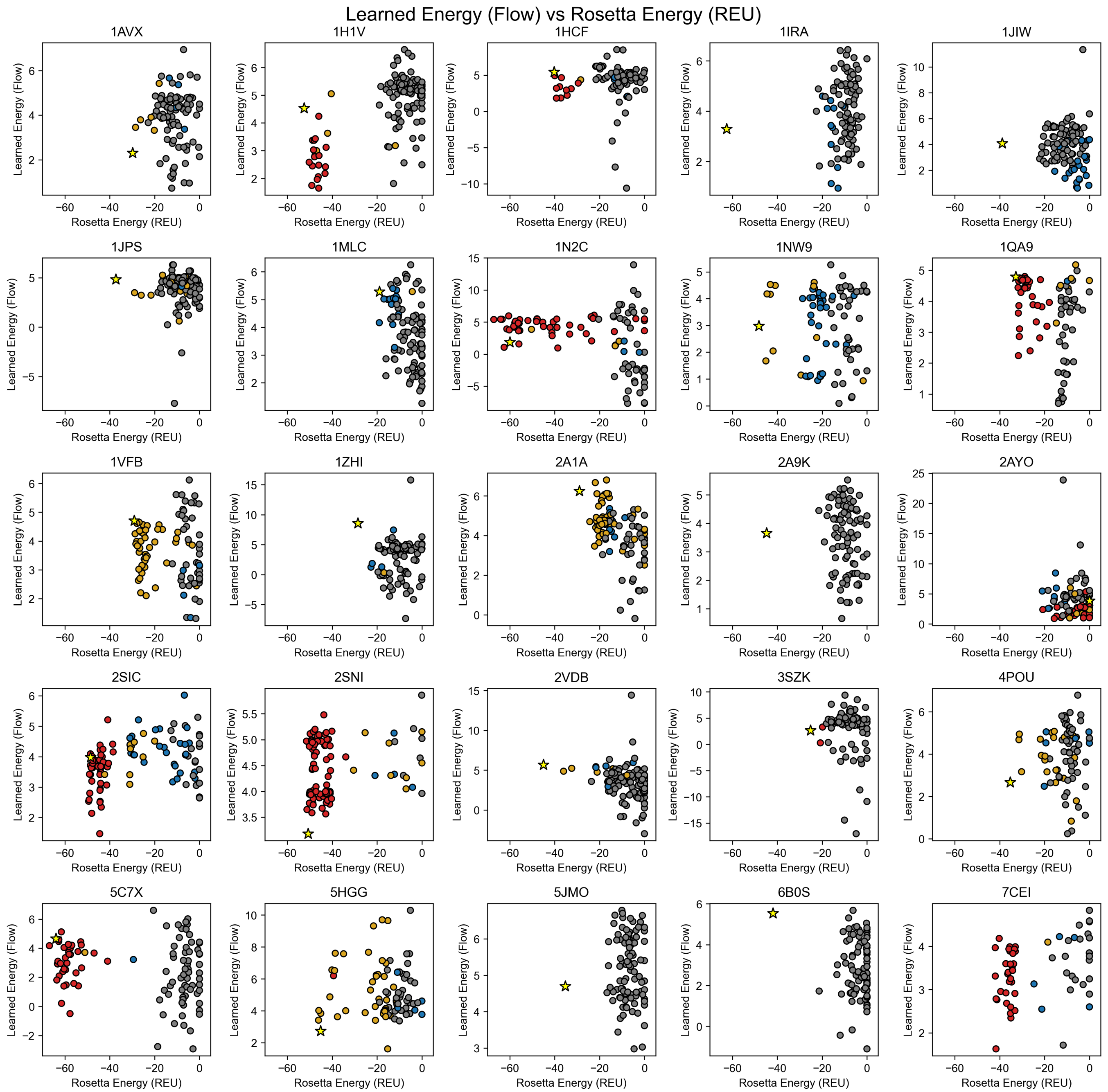


FIG. S14. Learned energy computed from integrating over flow trajectories, plotted against Rosetta energy of 120 docking poses generated from DFMDock for 25 targets in the DB5.5 dataset. Individual points are colored by their docking quality based on the CAPRI classification (incorrect: gray, acceptable: blue, medium: gold, high: red). Energy of the ground truth structure is shown as a yellow star.

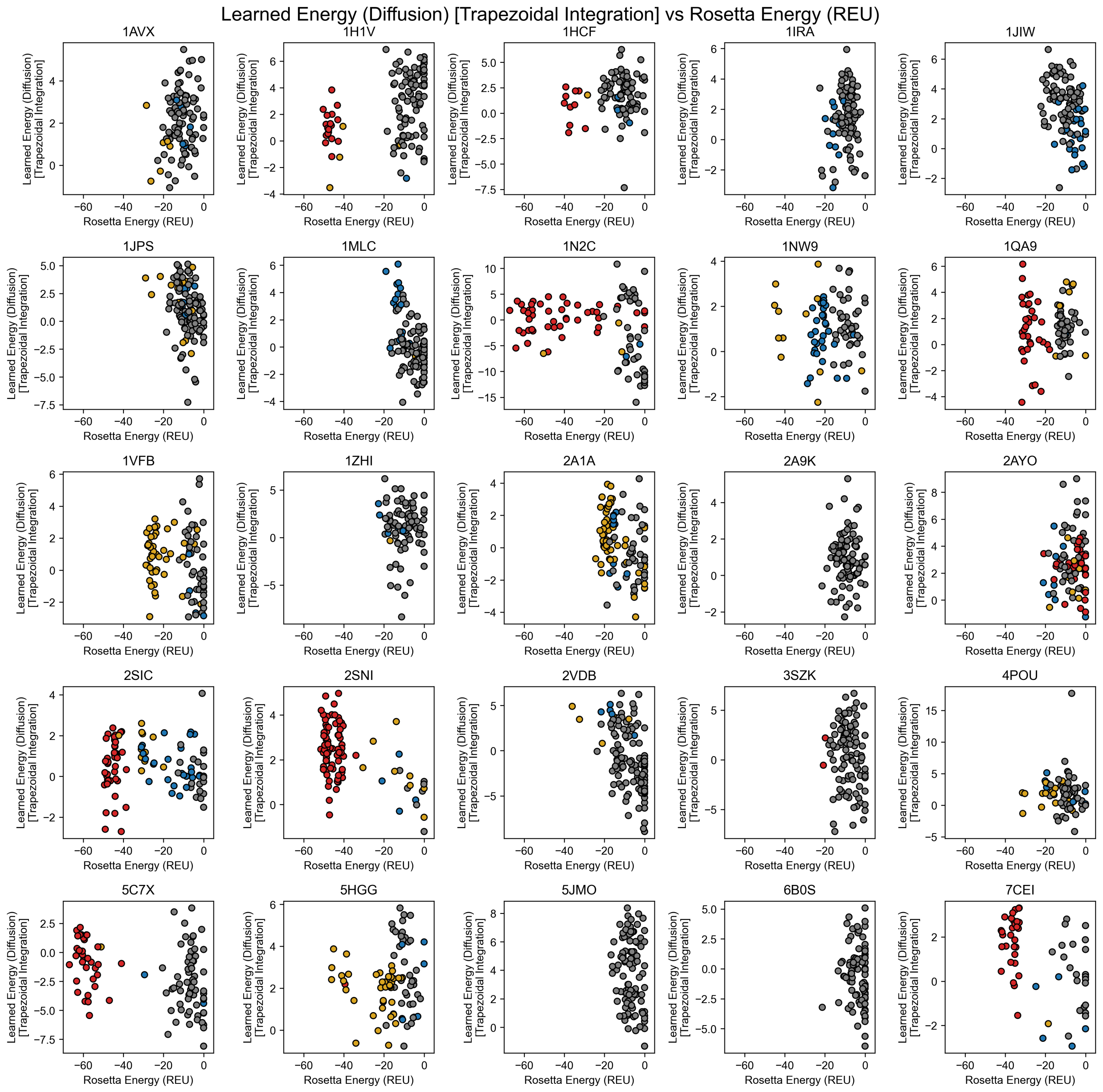


FIG. S15. Learned energy computed from integrating over diffusion trajectories with the trapezoidal method, plotted against Rosetta energy of 120 docking poses generated from DFMDock for 25 targets in the DB5.5 dataset. Individual points are colored by their docking quality based on the CAPRI classification (incorrect: gray, acceptable: blue, medium: gold, high: red).

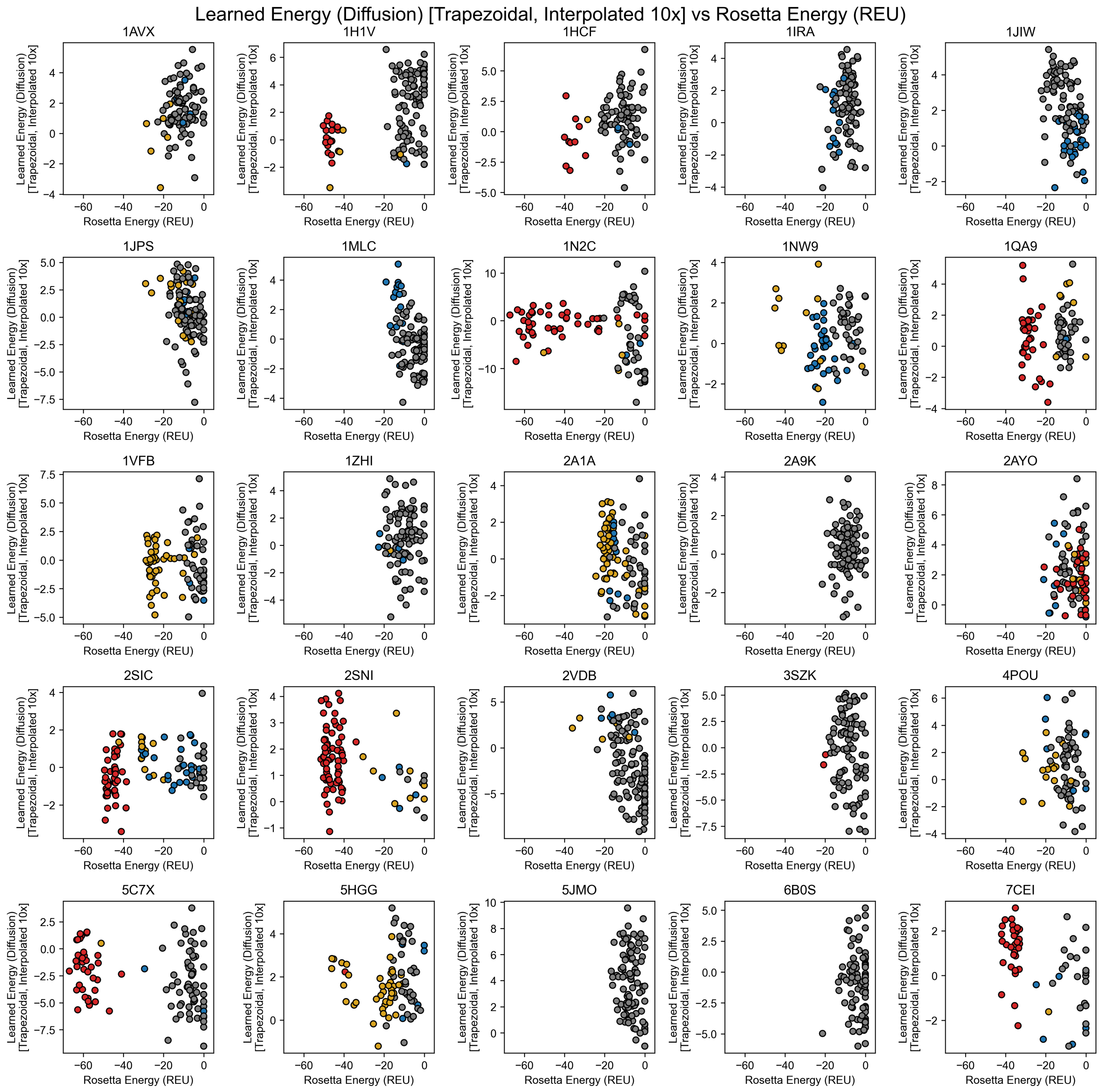


FIG. S16. Learned energy computed from integrating over diffusion trajectories with the interpolated trapezoidal method, plotted against Rosetta energy of 120 docking poses generated from DFMDock for 25 targets in the DB5.5 dataset. Individual points are colored by their docking quality based on the CAPRI classification (incorrect: gray, acceptable: blue, medium: gold, high: red).

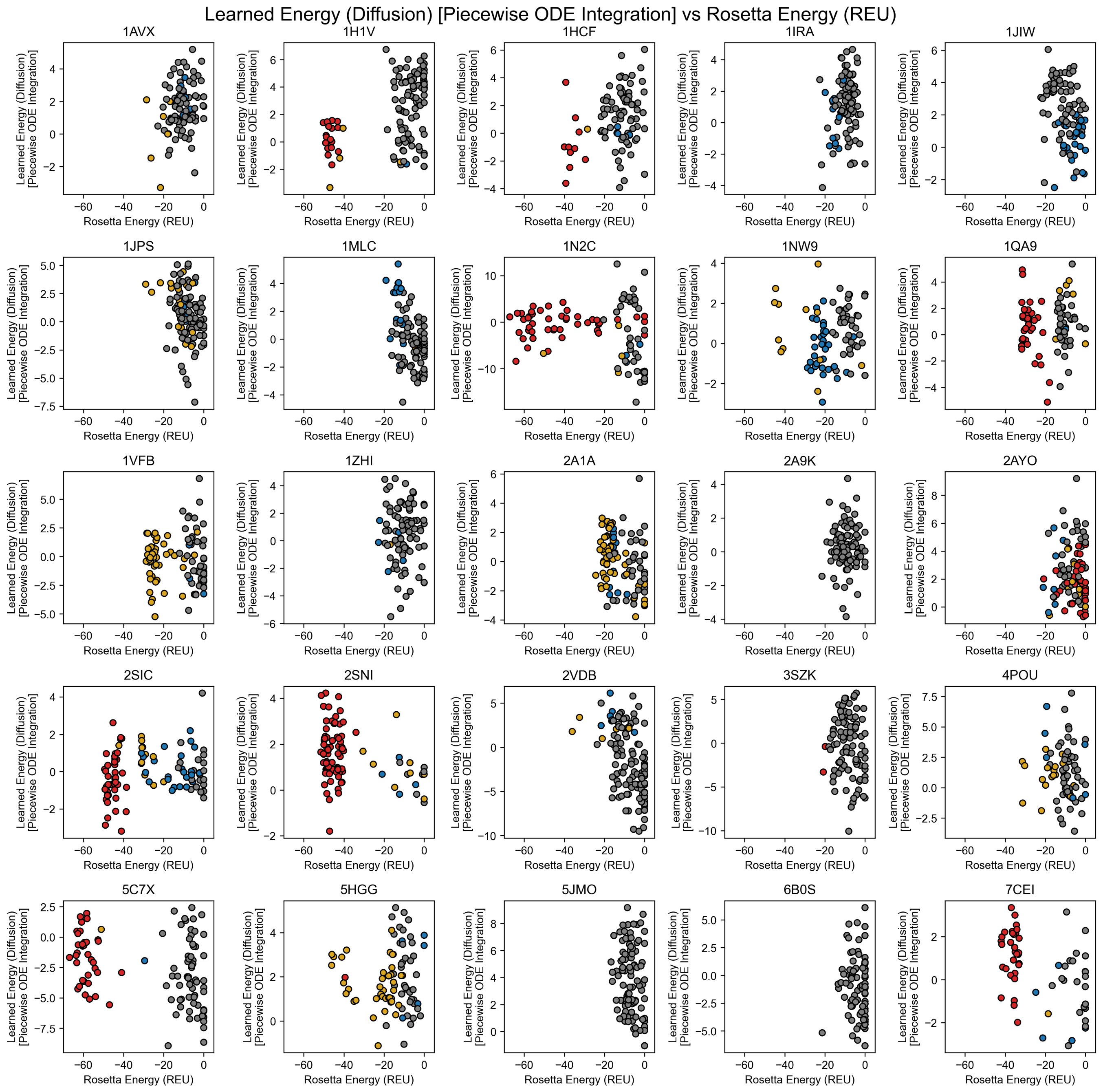


FIG. S17. Learned energy computed from integrating over diffusion trajectories with the piecewise ODE method, plotted against Rosetta energy of 120 docking poses generated from DFMDock for 25 targets in the DB5.5 dataset. Individual points are colored by their docking quality based on the CAPRI classification (incorrect: gray, acceptable: blue, medium: gold, high: red).

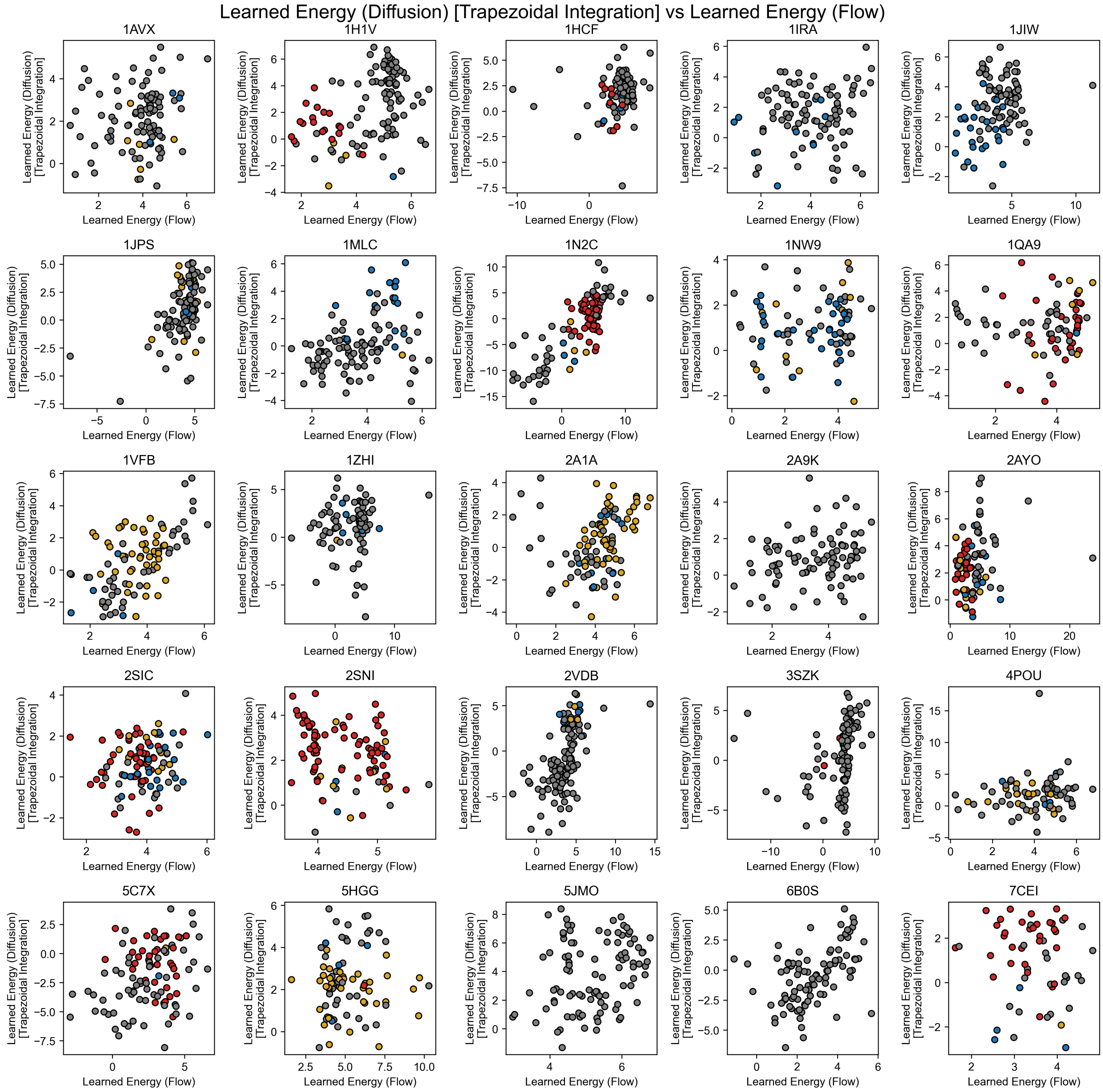


FIG. S18. Learned energy computed from integrating over diffusion trajectories with the trapezoidal method, plotted against integrating over flow trajectories generated from DFMDock for 25 targets in the DB5.5 dataset. Individual points are colored by their docking quality based on the CAPRI classification (incorrect: gray, acceptable: blue, medium: gold, high: red).

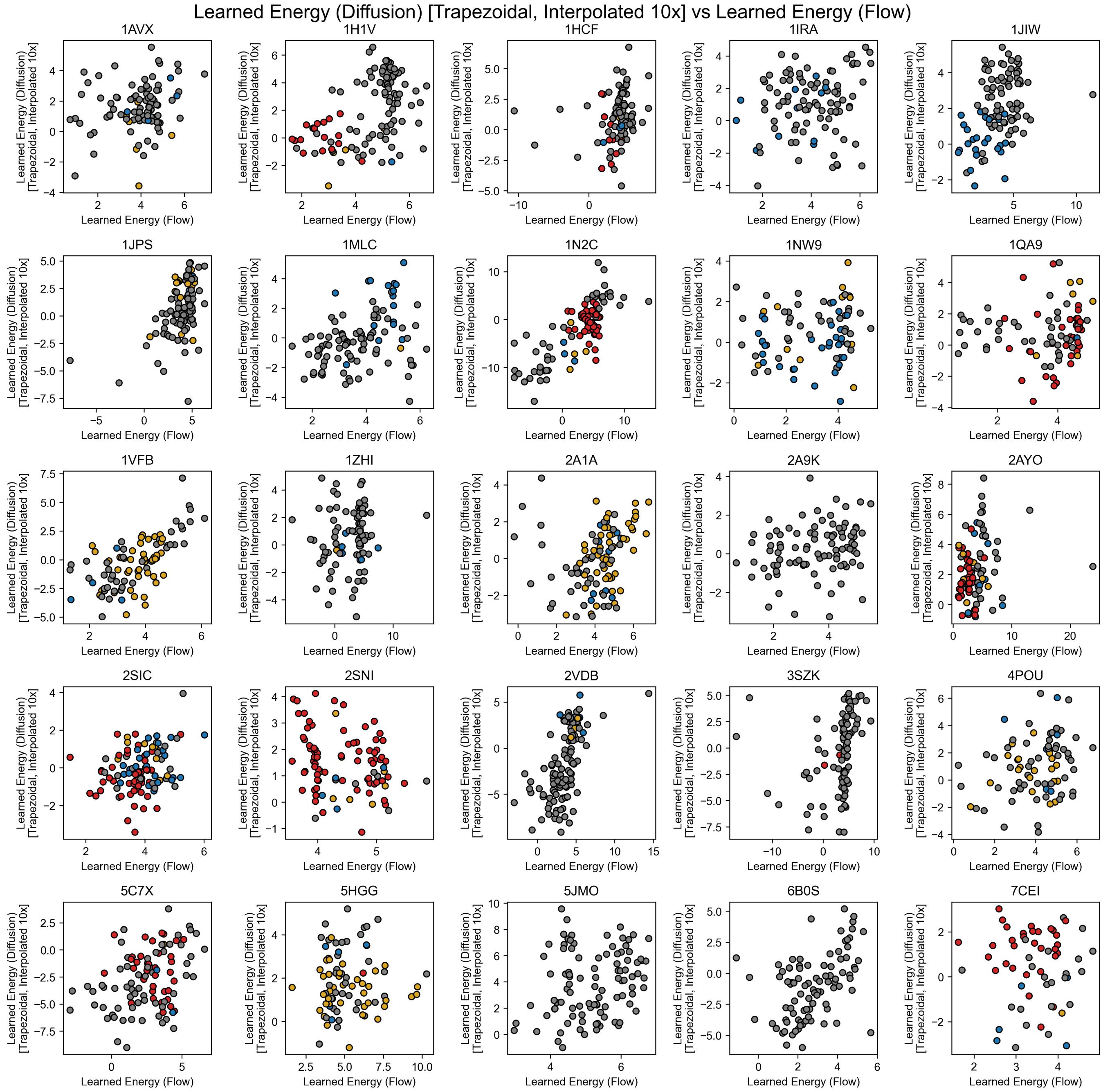


FIG. S19. Learned energy computed from integrating over diffusion trajectories with the interpolated trapezoidal method, plotted against integrating over flow trajectories generated from DFMDock for 25 targets in the DB5.5 dataset. Individual points are colored by their docking quality based on the CAPRI classification (incorrect: gray, acceptable: blue, medium: gold, high: red).

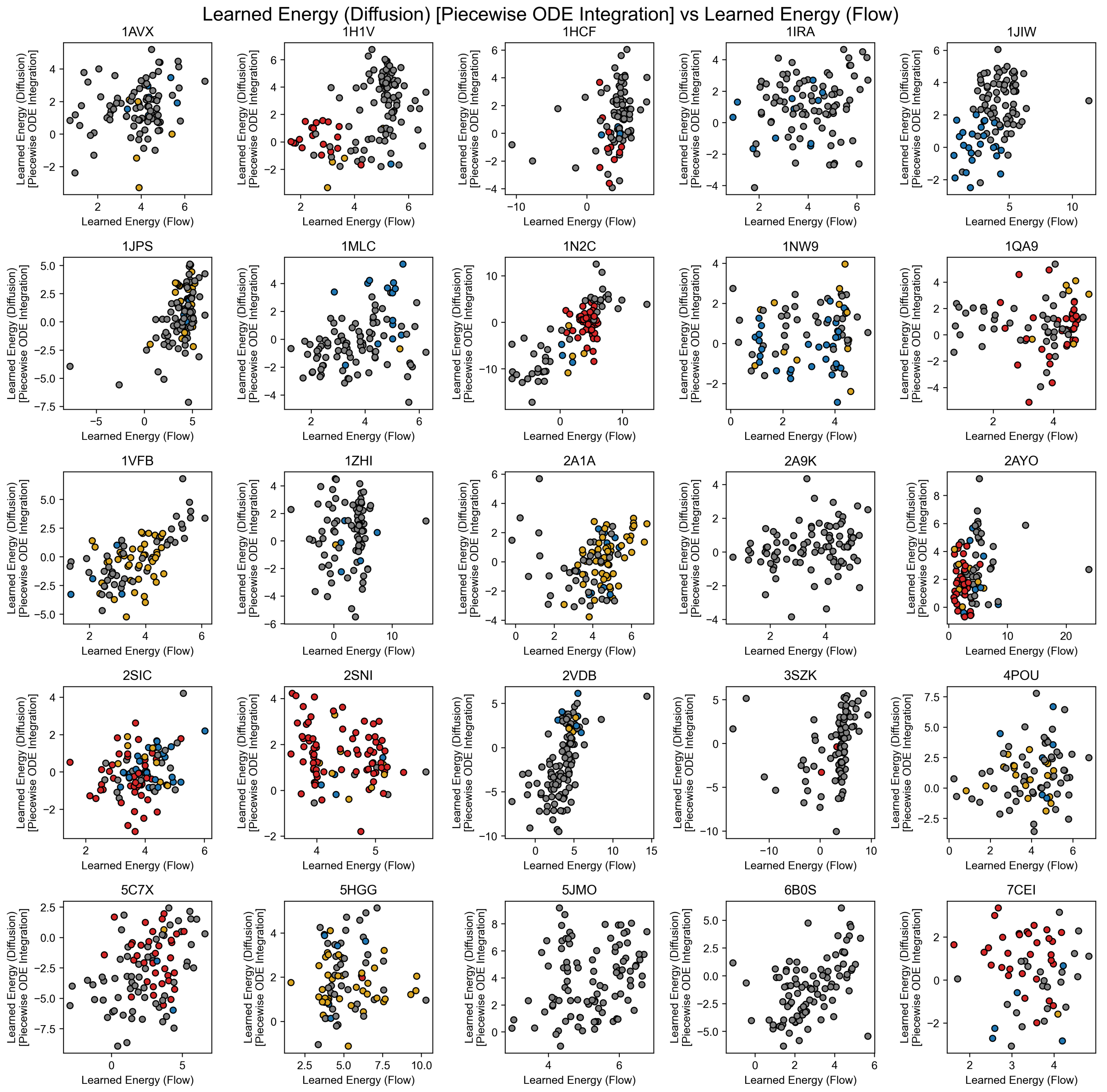


FIG. S20. Learned energy computed from integrating over diffusion trajectories with the piecewise ODE method, plotted against integrating over flow trajectories generated from DFMDock for 25 targets in the DB5.5 dataset. Individual points are colored by their docking quality based on the CAPRI classification (incorrect: gray, acceptable: blue, medium: gold, high: red).
